# Chromatin-dependent motif syntax defines differentiation trajectories

**DOI:** 10.1101/2024.08.05.606702

**Authors:** Sevi Durdu, Murat Iskar, Luke Isbel, Leslie Hoerner, Christiane Wirbelauer, Lukas Burger, Daniel Hess, Vytautas Iesmantavicius, Dirk Schübeler

## Abstract

Transcription factors recognizing short DNA sequences within gene regulatory regions are crucial drivers of cell identity. Despite recent advances, their specificity remains incompletely understood. Here, we address this by contrasting two TFs, NGN2 and MyoD1, which recognize ubiquitous E-box motifs yet instigate distinct cell fates—neurons and muscles, respectively. Following controlled induction in embryonic stem cells, we monitor binding across differentiation trajectories, employing an interpretable machine-learning approach integrating pre-existing DNA accessibility data. This reveals a chromatin-dependent motif syntax, delineating both common and factor-specific binding and predicting genome engagement with high precision. Shared binding sites reside in open chromatin, locally influenced by nucleosomes. In contrast, factor-specific binding in closed chromatin involves NGN2 and MyoD1 acting as pioneer-factors, influenced by multi-motifs, rotational spacing, flanking sequences, and specific interaction partners, accounting for subsequent lineage divergence. Extending our methodology to other models demonstrates how such combination of opportunistic-binding and context-specific chromatin opening underpin transcription factor specificity driving differentiation trajectories.

## INTRODUCTION

Eukaryotic transcription factors (TFs) play an important role in creating cell-type diversity in multicellular organisms, mediating selective activation of genomic regions. Unlike their prokaryotic counterparts, most eukaryotic TFs typically bind to DNA sequence motifs that are low in information content, due to being short and degenerate, set against the backdrop of a vast genome ^1^. This presents a paradox: among the millions of potential binding sites, only a few thousand are actually occupied by TFs. This indicates that the mere presence of a cognate motif is not a reliable predictor of eukaryotic TF binding. Yet these TFs that recognize short and highly similar cognate motifs interact with the genome with high precision and direct diverse cellular outcomes. This is exemplified by the basic-helix-loop-helix (bHLH) TF family with over 100 members that dimerize through bHLH domains and bind to the CANNTG motif known as E-box. E-box motifs occur more than 10 million times in mammalian genomes. Despite the low information content of the motif, the bHLH TFs drive distinct cellular differentiation programs such as neurogenesis by Neurogenin 2 (NGN2) and myogenesis by MyoD1 ^2–4^. The DNA binding preferences of bHLH-TFs have been shown not to be sufficient to account for the bound subset among the vast number of potential sites ^5^ nor to explain the differences in binding patterns between them^6^. While it is generally assumed that chromatin provides a barrier that masks the millions of unbound sites, it remains largely unclear at the level of individual loci what determines specificity in binding. If these abundant motifs are all masked by chromatin, how would such open chromatin dependent, and thus opportunistic binding enable the activation of the correct set of silent genes which seems to be a requirement to drive cell fates?

These observed specific binding patterns of the TFs are prevalently described as ’context-dependent’, and potentially driven by two primary mechanisms: direct recruitment through protein-protein interactions and indirect recruitment via chromatin that is made permissive by other TFs ^7^. Although direct interactions can enhance the combinatorial assembly of TFs at specific genomic locations, they only account for a minor portion of TF localization due to the considerable distances between co-occurring motifs ^8^. The second model proposes that “pioneer factors” or chromatin “insensitive” factors, capable of nucleosome binding, modify the chromatin structure to facilitate access for other TFs, thus influencing their targeting in a cell-type specific manner. Although these factors can open and modify a subset of their potential target-sites upon expression, contrary to expectations, pioneer factors do not consistently bind to identical sites across different cell types nor do they occupy or activate all their potential cognate motifs ^9–12^ This variability points to a context-dependence that remains poorly understood, highlighting a significant gap in our understanding of genomic targeting processes ^13^. To gain insights into the underlying mechanisms of TF targeting, it is necessary to implement systems that allow for controlled modulation of TF activity and monitoring binding as a function of chromatin state prior to the TF’s presence. While lineage-specifying TFs have been examined in dynamic conditions, such as developing embryos ^14^, the inherent heterogeneity of these systems complicates the precise control of TF timing and the assessment of chromatin states both prior to and following TF activation. Given that TF binding both influences and is influenced by the chromatin state, temporal resolution in modulating TF activity appears crucial. In response to these limitations, several studies have implemented induced TF expression and explored the concepts of TF binding preferences, pioneering and chromatin opening activity, albeit at later states of cellular transition ^6,15–18^.

With these challenges in mind, we established homogenous and comparable differentiation models in mouse embryonic stem cells driven by controlled induction of two bHLH TFs, NGN2 and MyoD1, resulting in the distinct generation of neurons and myocytes, respectively. This allows the investigation of the time-resolved genomic binding and subsequent responses of two similar short-motif recognizing TFs in relation to the DNA sequence and chromatin state. By measuring the immediate genomic binding of each factor starting from the same cellular origin, we identified the main determinants of chromatin engagement. These determinants underlie the specific binding patterns of NGN2 and MyoD1 among millions of their potential binding sites. By employing machine learning techniques, we uncovered a chromatin-dependent motif syntax with high predictive value that is composed of pre-existing DNA accessibility, motif variations including flanking bases, motif occurrence, and their relative positions. NGN2 and MyoD1 open chromatin depending on single base-pair differences in their motifs, with patterns that surprisingly differ from their mere binding strength. Cellular and in vitro assays reveal that other transcription factors, as well as NGN2 and MyoD1 dimerization-partners, differentially interact with these motif variants. The specific activities of NGN2 and MyoD1 on different motif variants led to downstream expression of distinct sets of TFs that further change genomic accessibility and in turn their binding patterns. Together, these insights provide a model explaining how short motif binding-TFs regulate chromatin with predictable specificity, underlined by a generalizable motif syntax. This uncovered motif syntax accounts for both pioneering and chromatin sensitive binding as well as cooperative regulatory actions at factor-specific versus shared target sites.

## RESULTS

### Controlled induction of NGN2 and MyoD1 in mouse embryonic stem cells drives reproducible and synchronous differentiation

To address how TFs with highly abundant and similar sequence motifs can distinctly guide specific lineages, we established a system in mouse embryonic stem cells (mESCs) allowing for the controlled induction of individual transcription factors.

Specifically, we employed NGN2, which directs mESCs towards a neuronal lineage **(Figure S1A-D)**, and contrasted it with MyoD1, one of the earliest identified lineage-driving TFs known to induce myocyte formation ^19–21^ **(Figure 1A)**. Both factors employ bHLH domains for DNA binding and recognize highly similar motifs.

**Figure 1.**
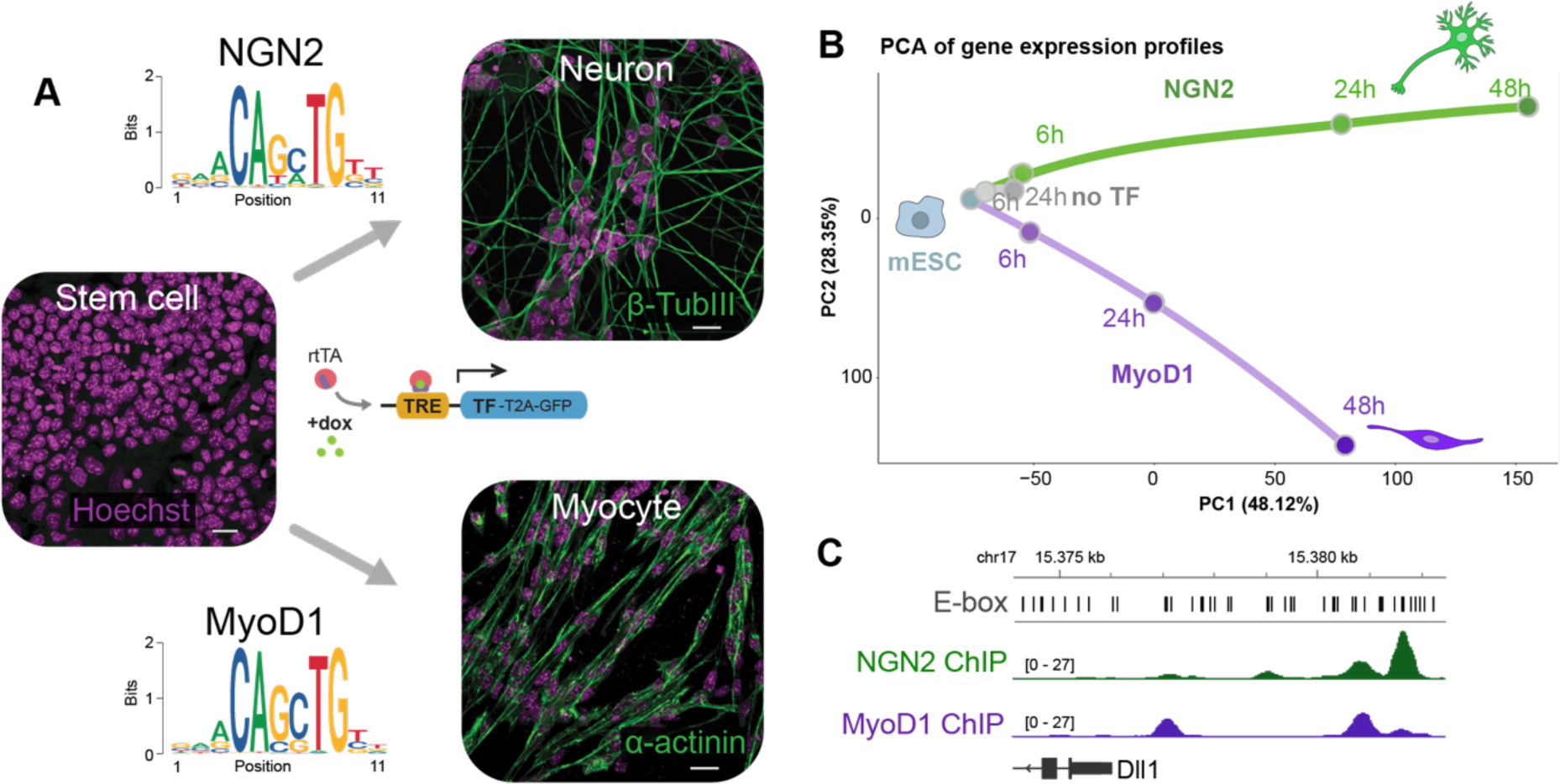
NGN2 and MyoD1 induces neurogenesis and myogenesis respectively, when expressed in mESCs, despite recognizing similar short DNA sequence motifs **(A)** Immunofluorescence images of mESCs prior and two-days post-induction of NGN2 and MyoD1, resulting in distinct cell types. Nuclei labeled with Hoechst in magenta, neuronal marker TubIII and myocyte marker α-Actinin in green. Scale bar, 20 µm. The motif logos represent the E-box motif that is de-novo identified by HOMER, considering the 500 most enriched ChIP-seq peaks (Figure S2A and B) **(B)** Differentiation trajectories illustrated by principal component analysis (PCA) of the transcriptomes along TF induction. mESCs are labeled in blue, NGN2 induction in green, MyoD1 in purple and no-TF control induction in gray (h: hours). **(C)** Genome browser tracks of the Dll1 gene locus illustrating shared and factor-specific binding by NGN2 and MyoD1 shown as ChIP-seq reads at 6h induction. E-box motifs are labeled in black bars.

For controlled induction, we engineered mESCs with the Tet-On system, incorporating stable transactivator expression and a single-locus insertion of the inducible epitope-tagged TFs followed by an in-frame cleaved GFP, serving as an indicator of expression (see Methods, ^22^). Doxycycline induced GFP expression alone without an additional TF expression (noTF) served as the induction control **(Figure S1E-H).** Upon NGN2 and MyoD1 induction, mESCs underwent synchronous differentiation into neurons or myocytes, within less than three days, as documented by time-lapse imaging. Within two days post induction, cell division ceased, and the characteristic neural or muscle-specific morphologies and cytoskeletal markers were detected (TUBB3 and α-Actinin) **(Figure 1A, Supplementary Video, Figure S1H and I).** Transcriptome analysis similarly revealed distinct gene expression trajectories, aligned with bona fide myogenic or neuronal differentiation **(Figure 1B, Figure S1J-L)**. Having established this controlled system, we next examined the initial genomic binding of NGN2 and MyoD1.

### Genomic engagement is a function of prior accessibility, motif frequency and motif sequence

To investigate genome-wide binding following TF induction, we profiled NGN2 using chromatin immunoprecipitation sequencing (ChIP-seq) at six hours after induction (6h) **(Figure 1C),** which represents the earliest time-point when robust levels of NGN2 are detected in most cells. A mutant form of NGN2 (NGN2mut) deficient in DNA binding, and GFP alone served as controls **(Figure S2A, Figure S1G).** The resulting genome-wide binding data revealed specific NGN2 peaks, including at expected targets like Neurod1 **(Figure S2A).** Combining multiple independent replicates identified 14116 consensus peaks as sites of NGN2’s initial chromatin engagement (**Figure S2C**). Among these, the typical E-box motif is highly enriched, with specific base pair preferences, especially in the central nucleotides. The most frequent central nucleotides of these NGN2 cognate motifs were GA, GC, TA, and GG **(Figure S2E,** see Methods ^23,24^). Notably, 95% of the peaks had an exact matching sequence to one of these motifs. In parallel we similarly profiled MyoD1 following its induction, which identified 13,266 bound consensus sites, similarly enriched for E-box motifs featuring frequent occurrences of GC and GG as central nucleotides **(Figure S2B, D and F)**.

The presence of these cognate motif sequences alone, despite their high enrichment in ChIP-seq peaks, falls short of explaining the specific binding patterns of NGN2 or MyoD1. As expected for a low complexity sequence and in contrast to the low number of bound sites, there are over a million occurrences of the NGN2 cognate motif in the mouse genome, leaving 99% unoccupied. We hypothesized that NGN2 and MyoD1 might preferentially bind sites within open chromatin. To explore this, we contrasted binding with genome-wide DNA accessibility prior to TF induction using DNase I hypersensitivity (DHS) mapping ^25^ **(Figure 2A** and **B).** This revealed that motifs residing in open chromatin, as defined by high DHS signal, are indeed more likely to be bound than those residing in closed chromatin. On average 1% of motifs in closed chromatin will be bound and 40% within accessible chromatin ranging from 0.4% to to 61% **(Figure 2A** and **C).** A similar pattern emerged with MyoD1: 54% of its cognate motifs in highly accessible chromatin were bound, compared to less than 0.2% in less accessible chromatin **(Figure 2D).** These findings suggest that pre-existing chromatin accessibility is a key determinant in whether NGN2 and MyoD1 are likely to engage with their cognate motifs. While binding to closed chromatin is a relatively rare event, these sites nevertheless accumulate to one third of all bound sites **(Figure 2A).** This is a direct reflection of the large number of motif sites residing in the closed chromatin state prior to TF induction. A comparative analysis of NGN2 and MyoD1 binding on their cognate motifs revealed that a majority of these lowly accessible but bound sites contains multiple cognate motifs that reside in close proximity to each other, which is rarely observed for the unbound or highly accessible bound sites **(Figure 2E** and **F).** This suggests that the presence of multiple motifs increases the likelihood of engaging in less accessible chromatin, conceivably by enhancing binding strength to a genomic sequence.

**Figure 2.**
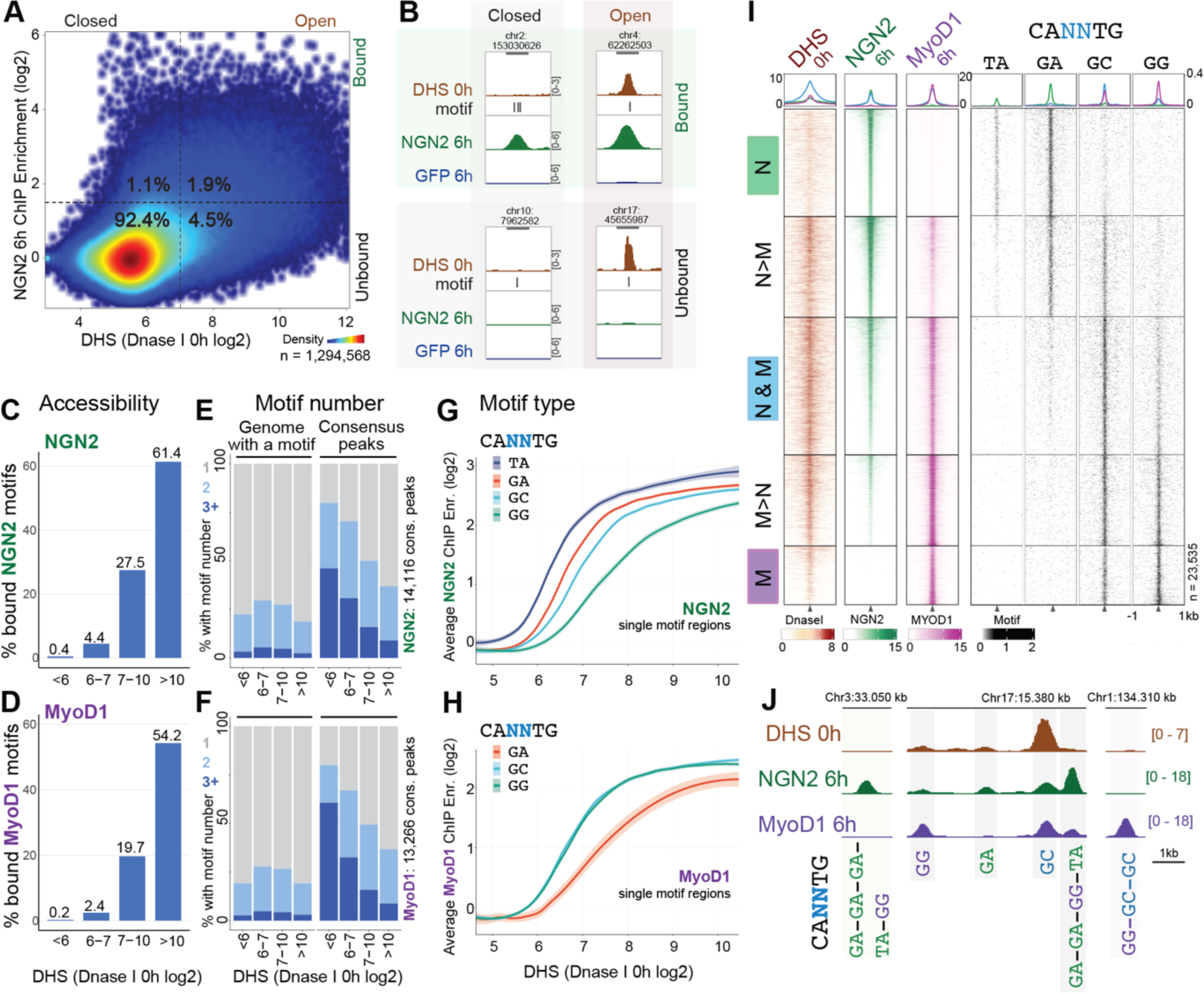
DNA accessibility and cognate motif occurrences are major factors underlying initial binding patterns **(A)** Density plot of all genomic regions that contain an NGN2 cognate motif with values corresponding to pre-existing DNase-seq (DHS 0h) counts and NGN2 ChIP-seq enrichment upon induction (NGN2 cognate motif sites:1.3 million 500bp genomic regions with at least one non-overlapping NGN2 motif described in FigureS2E). Numbers represent the percentage of total sites present within the indicated quarters. This shows that the majority of NGN2 cognate motifs reside in closed chromatin and are also not bound by NGN2 upon induction. Those heterocromatic regions that will be bound nevertheless make up one third of all NGN2 consensus peaks. **(B)** Example cognate motif sites of open-bound, closed-bound, open-unbound and closed-unbound categories are visualized as genomic tracks with values of 0h DNase-seq, NGN2 motifs in the 500 bp region, 6h NGN2 ChIP-seq and control GFP ChIP-seq. **(C** and **D)** Percentage of bound cognate motif sites (above enrichment 1.5) grouped in bins according to their accessibility prior to induction, showing higher likelihood of binding (>50%) in open regions and very low likelihood (<1%) in closed regions, indicating the impact of DNA accessibility. **(E** and **F)** Frequency of multiple motifs in consensus ChIP-seq peaks and rest of the cognate motif sites in the genome, grouped in bins according to their accessibility prior to induction (gray: one cognate motif, light blue: two and dark blue: three or more cognate motifs). For both NGN2 and MyoD1, closed but bound sites have a higher number of motifs. **(G** and **H)** Fitted line plots of average ChIP-seq enrichment versus DNA accessibility (DHS 0h) of cognate motif sites with a single motif (residing in consensus ChIP-seq peaks and same number of matched backgrounds), grouped by the most frequent central nucleotides: GA, GC, TA, GG variants for NGN2 and GC, GG, GA variants for MyoD1. The differences in the curves illustrate the difference in the likelihood to be bound of a motif variant depending on the accessibility of the region. **(I)** Combined NGN2 and MyoD1 peaks illustrated as a heatmap. Peaks strongly bound by both factors (N&M) in blue, peaks exclusively bound by one factor (N in green or M in purple), and peaks preferentially bound by one of the two (N>M or M>N). DNase-seq in red, NGN2 ChIP-seq in green MyoD1 ChIP-seq in purple and the presence of cognate motif variants depending on the central nucleotides (TA, GA, GC, GG) in black. **(J)** Genomic regions with exclusive or overlapping NGN2 and MyoD1 binding, exemplifying the effect of chromatin accessibility and occurrence of cognate motif variants present at the peak centers (motifs are represented by the two central nucleotides of the E-box). NGN2 preferred motifs are colored in green, MyoD1 preferred motifs in purple.

Next, we asked how the likelihood of engaging single motif occurrences relates to particular motif variants. To test this, we analyzed NGN2 and MyoD1 binding enrichments on the genomic regions with single motif occurrences as a function of their initial DNA accessibility. This revealed differences in binding profiles at E-box motif variants when grouped by their central nucleotides. While all the motif variants are comparably bound when residing in accessible chromatin, differences are apparent in less accessible chromatin **(Figure 2G** and **H).** This reflects NGN2’s ability to engage E-box motif variants, ranked from highest binding potential to lowest as: TA, GA, GC and GG central bases **(Figure 2G, Figure S2G)**. Such difference is not limited to NGN2 as it is similarly evident for MyoD1, which also shows comparable binding within open chromatin while in less accessible chromatin a strong preference is observed for GG and GC as central nucleotides compared to GA **(Figure 2H)**. These data argue that for both TFs, variations in the motifs are relatively less inconsequential for binding in open chromatin, while they appear to be critical for engaging closed chromatin.

We next addressed if other preexisting chromatin features such as histone modifications might influence binding ^10,26^. Contrasting well-characterized histone marks reveals that modifications that are hallmarks of open chromatin such as H3K4 methylation indeed correlate with binding. Importantly, however, none of them contribute any additional predictive information beyond that already provided by DHS **(Figure S2H and I).** This suggests that in the case of NGN2 and MyoD1, individual histone marks themselves are unlikely to account for global motif engagement independent of DNA accessibility.

If the genomic binding patterns of NGN2/MyoD1 are indeed a function of motif occurrence and DNA accessibility, then the pronounced contrasts in binding are expected to occur in genomic sites that are initially less accessible but have high occurrences of NGN2 and MyoD1’s individually preferred motifs. Conversely, the similarities should be driven by regions with higher accessibility, or where different preferred motifs are situated adjacent to each other, thus bound by both NGN2 and MyoD1. Contrasting the bound regions by grouping NGN2 and MyoD1 binding sites indeed illustrates a general overlap at highly accessible regions and distinct binding at sites with lower accessibility which are enriched for factor specific motif variants **(Figure 2I, Figure S2J).** These patterns can be observed at individual loci as illustrated by a low accessible region with multiple GA motifs (E-box motifs with GA central nucleotides) that is distinctly bound by NGN2 or a low accessible region, containing one GG and two GC motifs, that is only bound by MyoD1. In contrast, a single GC motif residing in accessible chromatin is bound by both factors **(Figure 2J, Figure S2K)**. These observations support a model in which chromatin accessibility explains the overlapping as well as the differential binding patterns between two lineage driving TFs as a local modulator **(Figure S2L).**

Although our analysis thus far identifies contributors to the distinct binding patterns of the two TFs, these components alone fail to account for many binding events, including those unbound motifs residing in accessible regions or with multiple motifs in proximity **(Figure 2B-F)**. This suggests the presence of additional features that our analysis has not captured, thus motivated us to use a self-learning approach with the aim of generating a model that can leverage the previously identified features, as well as potential additional features to predict binding.

### Convolutional neural networks uncover a chromatin-dependent motif syntax that underlies TF binding

To gain further insights into the underlying features accounting for TF binding, we adopted a machine learning approach, and built a convolutional neural network (CNN) model that uses DNA sequence combined with chromatin accessibility prior to TF expression as input **(Figure 3A)** (see Methods ^27,28^). Training this CNN model with NGN2 and MyoD1 ChIP-seq enrichments predicted the degree of binding remarkably well, with the correlation between prediction and experimental data being comparable to the correlation between experimental replicates **(Figure S3A and B, Figure S2C and D**).

**Figure 3.**
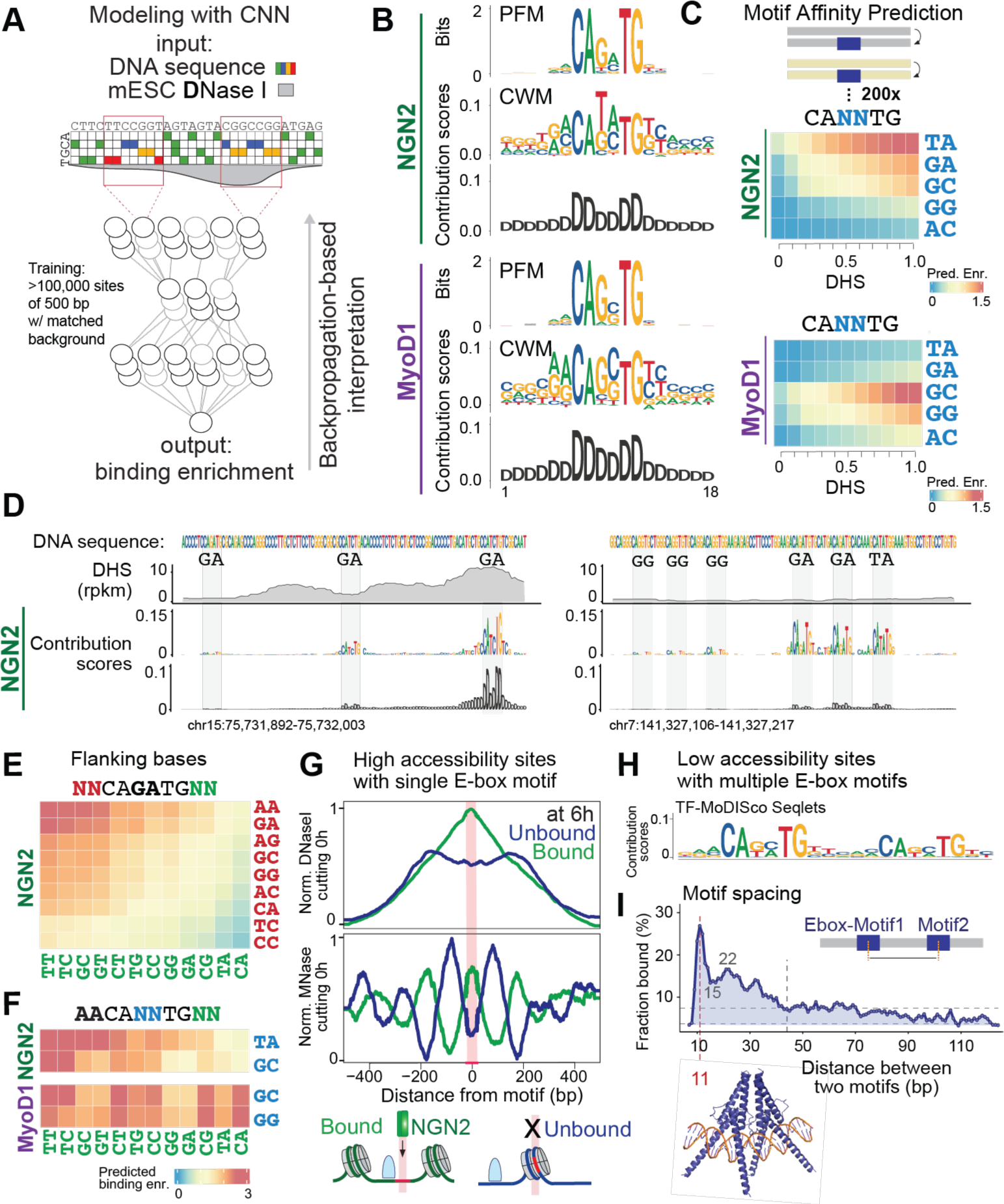
Identification of a chromatin-dependent motif syntax using a convolutional neural network (CNN) **(A)** Scheme of the applied CNN to predict binding of NGN2 and MyoD1 upon induction, using DNA sequence and continuous DNase I signal in mESCs as input. **(B)** Contrasting frequency-based motif representation (PFM) that informs about the frequency of the DNA bases in the enriched motifs in NGN2 peaks, and contribution weight matrices (CWM) as identified by the CNN model that informs about the contribution of the DNA bases to the degree of binding enrichments. DNA sequences are represented with the letters A,C,G,T and accessibility with the letter D. The differences in PFM and CWM highlight the binding strength contribution of flanking nucleotides and less frequent central nucleotides. The contribution of DNA accessibility to the binding prediction shows high contribution directly overlapping with the E-box motif and lower contribution of the surrounding. (n=26,128 E-box instances for NGN2 and 25,327 for MyoD1). **(C)** Calculation of NGN2 and MyoD1’s relative affinities to E-box motif variants independent of the genomic sequence context at any given accessibility using the CNN model on synthetic sequences (average gained binding strength upon placing the same motif across various backgrounds with simulated accessibility). The predicted binding strengths of NGN2 and MyoD1 to the motif variants across different initial accessibility are represented in colors from blue to red. Central nucleotides of the E-box motifs are colored in blue. **(D)** Single locus display of DNA sequence, DNase I accessibility profile and their deconvolved contribution scores for NGN2 binding prediction, reporting on the features accounting for the binding strength. **(E)** Predicted binding strength contribution of the flanking-bases (in red for 5’, in green for 3’) that are represented in the E-box motif summary models, shown for CAGATG motif. **(F)** Contribution of the flanking-bases to NGN2 and MyoD1 binding prediction at the E-box motifs with different central nucleotides. Central nucleotides are colored in blue and 3’ flanking nucleotides in green. **(G)** Comparison of pre-induction MNase and Dnase I cutting profiles at NGN2 cognate motif sites residing in open chromatin grouped as sites that are bound or not bound by NGN2 upon 6h induction. **(H)** DNA sequence motif (seqlet, identified by TF MoDISco) composed of two E-box sequences that is informative for NGN2 binding prediction at bound sites with low initial accessibility. **(I)** Fraction of NGN2 bound sites among initially less accessible regions with two NGN2 motifs at various distances (bp distance between motif centers). Inner panel: modeled NGN2 binding (AlphaFold 3) on two NGN2 motifs with 11 bp center to center distance, which gives high binding potential.

To identify features influencing transcription factor binding predictions ^29^, we utilized DeepLIFT ^27^ and quantified the contribution scores of the DNA sequences and chromatin accessibility. Based on these contribution scores, we further extracted DNA sequence motifs using TF-MoDISco ^30^ (see methods). Consistent with the previous analysis, this approach identified E-box motifs as the major DNA sequence component for the prediction, as well as other previously known motifs of synergizing TFs such as Pbx for MyoD1 **(Figure 3B, Figure S3C)** ^6,31^. In addition to the motif sequence, the model predicts that its local accessibility significantly contributes to the binding. This should enable the further use of the CNN model to estimate the chromatin binding strengths for individual E-box motif sequences (i.e. motif variants), isolated from the genomic sequence context. To do so, we averaged the predicted TF binding on E-box motifs embedded in hundreds of in silico generated background sequences (as in affinity distillation approach ^32^) at varying initial accessibilities (see Methods) (**Figure 3C, Figure S3D).** This is exemplified by examining the motif variants with only a few isolated binding instances such as CATATG and CAACTG motifs for NGN2 and MyoD1 respectively (**Figure 3B** and **C)**. This shows that the model can estimate the likelihood of TF binding to a given DNA sequence as a function of its initial accessibility, which is in line with the observations on the motif variant binding preferences in **Figure 2G** and **H**.

Visual inspection of the genomic tracks of the contribution scores calculated for DNA sequence and accessibility illustrates that the CNN model assigns different degrees of importance to E-box motifs depending on sequence variations and their local accessibility **(Figure 3D, Figure S3E)**. The identified sequence variations were not limited to the motif itself but also included flanking bases that in several cases appear to be even more relevant for binding than the central two bases. Calculating the binding strengths revealed that even a single base change at the flanks can impact the likelihood of being bound within a wide dynamic range **(Figure 3E)**. This impact of the flanking nucleotides varies between NGN2 and MyoD1 potentially further accounting for their differences in initial genomic binding. Interestingly, the flanking nucleotides’ influence depends on the central bases hinting at a potential crosstalk between these bases for binding preference **(Figure 3F)**.

Although binding occurs more frequently in open chromatin, many accessible sites remain nevertheless unbound **(Figure 2A-C)**. What enables the model to accurately predict the binding at accessible sites beyond variations in the motif sequence?

Intrigued by single locus inspections (**Figure 3D, Figure S3E**) we hypothesized that nucleosome positioning, which has previously been shown to influence TF binding also for bHLH factors, could be a contributing feature ^33^. The actual nucleosome phasing within open chromatin regions, as determined with our previous MNase measurements in mESCs ^34^, indeed differs between bound and unbound sites **(Figure 3G)**. Motifs that will not be bound by NGN2 upon induction tend to overlap more frequently with a nucleosome, whereas sites bound upon induction are more likely to reside between nucleosomes. As these sites lie within accessible regions, we presume that this phasing results from the activity of other TFs ^35,36^, and that the DNase I signal carries information about it ^37^. The resulting binding hindrance of E-boxes by phased nucleosomes is in line with the finding that bHLH proteins do not bind to the nucleosome core in vitro ^33^.

Within less accessible regions, the CNN model discerns a motif syntax of two E-box motifs in line with higher motif occurrences described above and in addition provides further insights into their relative position **(Figure 2E** and **F, Figure S3D, Figure 3H),** hinting to a potential synergy between motifs. To test this, we predicted the cooperativity of two motifs as a function of their distance, using in-silico generated sequences ^38^. This revealed a wide range of impact with higher cooperativity between two motifs that reside close to each other **(Figure S3F).** Comparing this prediction with the measured binding preferences confirmed that shorter distances are enriched for binding at less-accessible regions **(Figure S3G and H)**.

Interestingly, within these closely spaced motifs, also the exact spacing matters. Inter-motif distances shorter than 9 base pairs (bp) did not enhance binding potential, possibly due to a molecular clash at the two motifs. The highest cooperativity is predicted at 10 to 12 bp distance corresponding to one helical turn of the DNA, whereas 15 bp, corresponding to an opposite facing on the DNA helix, was less favorable **(Figure S3F).** The actual binding measurements confirm this prediction as they revealed a similar preference for shorter distances within 40 bp, up to 100 bp. It further supported the reduced binding likelihood at a motif distance below 9 bp, yet increased binding at 11 bp, and again a decrease at 15 bp **(Figure 3I, Figure S3I)** (modeled with AlphaFold3 ^39^). This argues that the spacing between motifs impacts binding potential, a feature critical at sites of lower accessibility to counteract the chromatin barrier.

Taken together, these results suggest that a motif syntax composed of motif variants including flanking bases, their local frequency and the underlying chromatin state accounts for the ability of a genomic region to facilitate TF binding. Motifs’ positions relative to phased nucleosomes further influence the likelihood to be bound within open chromatin while at less accessible regions higher motif numbers with optimal inter-motif spacing enable binding **(Figure S3J)**.

### The abilities of NGN2 and MyoD1 to open chromatin and activate genes differs depending on the motif variant

Having thus far focused on initial binding we next asked how NGN2 and MyoD1 themselves impact chromatin and transcriptional activity downstream of binding. More specifically, we measured accessibility by ATAC-seq, acetylation of lysine 27 of histone H3 (H3K27ac) and presence of RNA polymerase II (Pol2) by ChIP-seq 6h post-induction. For visualization, we employed dimensionality reduction (UMAP) of the relevant genomic regions, that are either bound by NGN2 or MyoD1 or regions that show changes in accessibility (see Methods) at any time point. **(Figure 4A)**. This positioned the genomic regions in an intuitive way, allowing inspection of shared regions and the regions exclusive for NGN2 or MyoD1.

**Figure 4.**
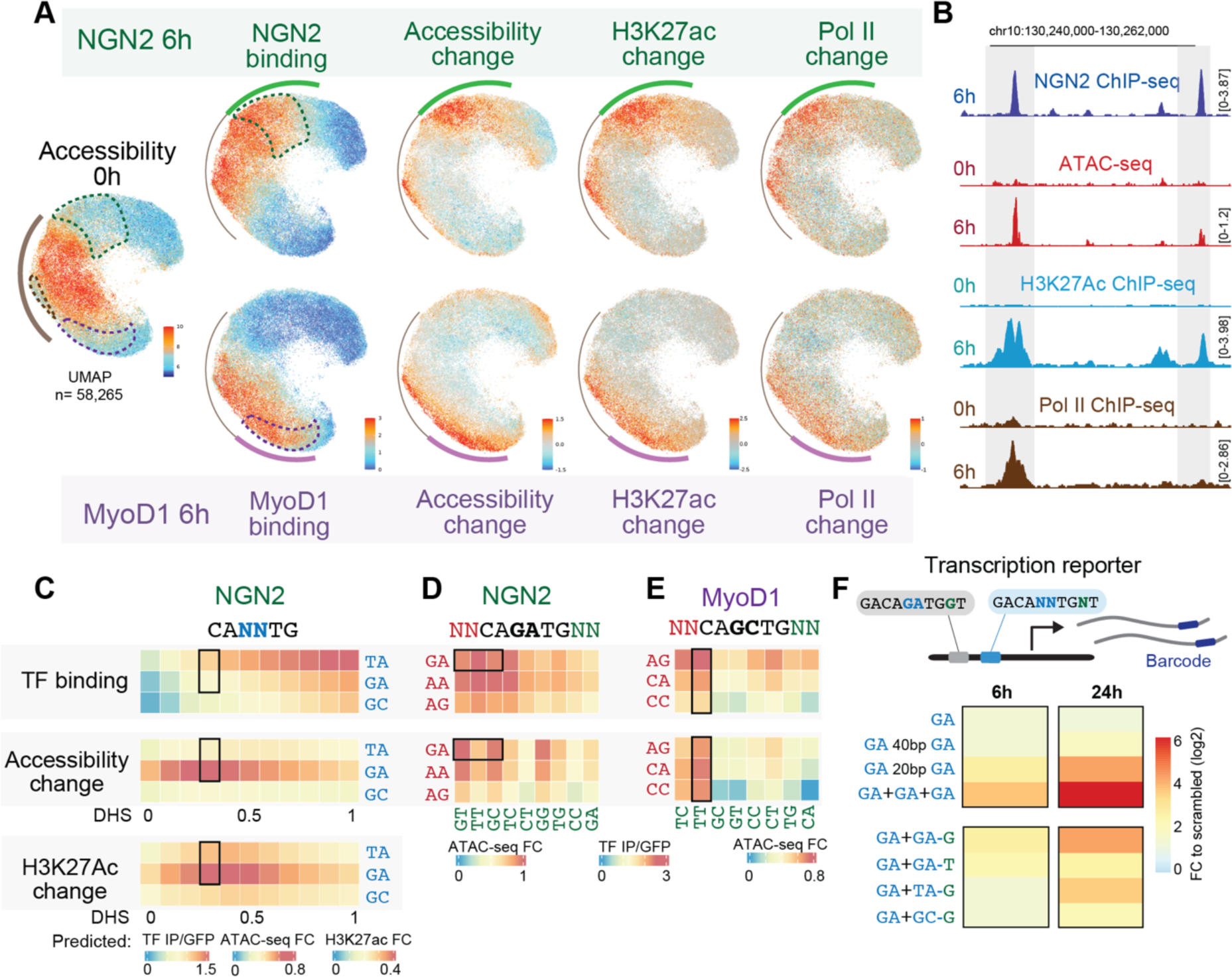
Motif variants lead to different chromatin and transcription responses **(A)** UMAP of relevant genomic regions (represented as single points that are positioned based on similarity of accessibility and NGN2/MyoD1 binding) displaying the measurements of initial accessibility, TF binding and fold change in ATAC-seq, H3K27ac ChIP-seq and Pol2 ChIP-seq, colored from blue to red. Bound regions with low initial accessibility or factor-specific binding are outlined with dashed lines. Curved lines highlight the common (brown), NGN2 specific (green) and MyoD1 specific (purple) binding. The patterns of binding and change in chromatin features reflect increased activity upon binding at initially low DHS sites, though with variable degrees (e.g. for NGN2, binding strength is more graded from left to right and chromatin opening from top to bottom at its TF-specific binding sites) **(B)** Genome browser tracks of an example genomic locus, showing regions with similar starting accessibility and similar NGN2 binding but with varying gain of activity upon 6h NGN2 induction. **(C)** Contrasting the binding and activity on the motif variants using CNN model predictions on synthetic sequences. Predicted binding, change in accessibility (ATAC-seq) and change in histone acetylation (H3K27ac ChIP-seq) upon NGN2 induction for NGN2 E-box motifs with TA, GA and GC central nucleotides across various starting accessibilities are shown as heatmap from blue to red. The outlined box at DHS 0.3 highlights the differences between the binding strength and its impact on chromatin for CAGATG and CATATG motifs (DHS 0.3, representing low initial accessibility and high dynamic range for gain of activity upon binding). **(D)** Contrasting predicted binding and accessibility change upon NGN2 induction at CAGATG motif (DHS 0.3) depending on the flanking bases. **(E)** Contrasting predicted binding and accessibility change upon MyoD1 induction at CAGCTG motif (DHS 0.3) depending on the flanking bases. **(F)** A library of transcriptional reporters inserted stably in a defined locus, read-out by the matched barcodes. Constructs designed to report the effect of NGN2 motif variations on gene expression upon NGN2 induction. GA: single GA motif, GA 40 GA: two GA motifs and 40 base-pairs between them. GA 20 GA: two GA motifs and 20 base-pairs between them. GA+GA+GA: three GA motifs. GA+NN: one GA motif and another motif with the indicated central nucleotide and three prime flanking nucleotide. In (C-F), central nucleotides of the E-box motif are colored in blue, flanking nucleotides in red for 5’, in green for 3’.

At the early 6h time-point, NGN2 and MyoD1 bound sites, which are initially less accessible, show an overall increase in accessibility, histone acetylation and RNA polymerase II **(Figure 4A)**, suggesting that NGN2 and MyoD1 primarily activate chromatin upon binding ^40^. Intriguingly, these activities vary between regions even when displaying similar binding enrichments for the same TF, hinting at a differential impact upon binding **(Figure 4B).** To investigate these differences further, we trained a CNN model on accessibility changes as measured by ATAC-seq, which identified the impact of initial accessibility and the occurrence of cognate motifs. While the binding potential for a genomic sequence is higher in more accessible chromatin (**Figure 4A**), further opening of chromatin as measured by ATAC-seq is primarily observed at sites of lower initial accessibility as might be expected **(Figure 4C)**. Like the binding potentials measured by ChIP-seq, this opening potential varied between motif variants but surprisingly not always in the same order as the strength of binding (**Figure 4C-E, Figure S4A-F**). For instance, while the CATATG motif had a higher binding potential, the CAGATG motif demonstrated a higher chromatin opening potential at the same initial accessibility **(Figure 4C, Figure S4A-C)**. Also, the flanking bases of the same motif variant exhibited different binding versus opening potentials, exemplified by the contrast between GT versus TT 3’ flanks of the CAGATG motif **(Figure 4D, Figure S4D, E and G)**. Similar differences were observed for MyoD1 activity on its cognate motifs **(Figure 4E)**.

To readily test whether these subtle single-base variations impact transcriptional output we generated over 20 barcoded reporter constructs for gene expression harboring different motif variations. These were inserted into a defined genomic locus in mESCs (see Methods ^22,41^) and expression measured following NGN2 induction **(Figure 4F, Figure S4H)**. This validated the expected increase of transcriptional output with higher motif density but also that the slight motif variations can differentially impact gene expression even when embedded in identical DNA sequence context, where G flanking CAGATG motif had the highest activity. This was in line with the enrichment of particular motif variants in NGN2 bound sites that are linked to the up-regulated neurogenesis-specific genes **(Figure S4I and J).**

Altogether, these results argue that subtle sequence differences have direct functional impact on chromatin and gene expression. Since these do not always correlate with the measured binding strength, we hypothesized that they could reflect binding by different TF complexes.

### Distinct TF combinations act differentially on the motif variants, increasing the regulation diversity by a single base difference

Many TFs can potentially engage with E-box motifs such as bHLH and zinc finger domain containing families ^4,42,43^. To ask which TFs might be interacting with the NGN2 preferred motif variants in our cellular system, we performed mass spectrometry using nuclear extracts from NGN2 induced cells, and double stranded 20bp DNA oligomers containing a single E-box motif as baits **(Figure S5A and B).** This detected multiple transcription factors including known bHLH domain containing E-box binders such as TCF proteins, MGA, MAX and TFAP4. Since dimerization is essential for bHLH domain-containing factors to interact with an E-box ^3,44^, we asked whether these proteins hetero-dimerize with NGN2 or MyoD1. Immunoprecipitation followed by mass-spectrometry identified TCF3 (TFE2/E2A), TCF4 (ITF2/SEF2) and TCF12 (HEB/HTF4) as potential protein partners for both TFs **(Figure 5A, Figure S5C)**. We hypothesized that the resulting dimers might have differential motif preferences, which we next tested in vitro to isolate the direct impact of the partnering. More specifically, we performed DNA affinity purification sequencing (DAP-seq ^45^). Recombinant purified TFs are incubated with sheared mESC genomic DNA, and bound DNA is subsequently sequenced (see Methods). To be comprehensive, we generated DAP-seq profiles for NGN2, MyoD1, TCF3E12, TCF3E47, TCF4 and TCF12 in all potential pairs that could be present in vivo **(Figure 5B)**. This identified combination-specific enrichment profiles for all 20 TF pairs, which clustered consistently according to protein dimers **(Figure S5D).** While the E-box motif was enriched among all tested protein dimers, we further uncovered differential motif variant enrichments, suggesting dimer-specific binding preferences **(Figure 5C)**. For example, the CATATG motif was preferentially bound by the NGN2 homodimer whereas the NGN2-TCF3E12 heterodimer showed preference for the CAGATG motif, in line with the observation that binding and opening potentials of these motif variants showed variability. Similarly, TCF homodimers preferred motifs starting with G flanks whereas TCF-MyoD1 combinations preferred A. This is in line with our previous observation that NGN2 and MyoD1 binding preferences were a function of both the central motif bases and flanking sequences, and that the influence of the flanks varied depending on the central bases **(Figure 3F).** It further suggests that these binding preferences might be driven by the differential interaction with distinct sets of protein partners.

**Figure 5.**
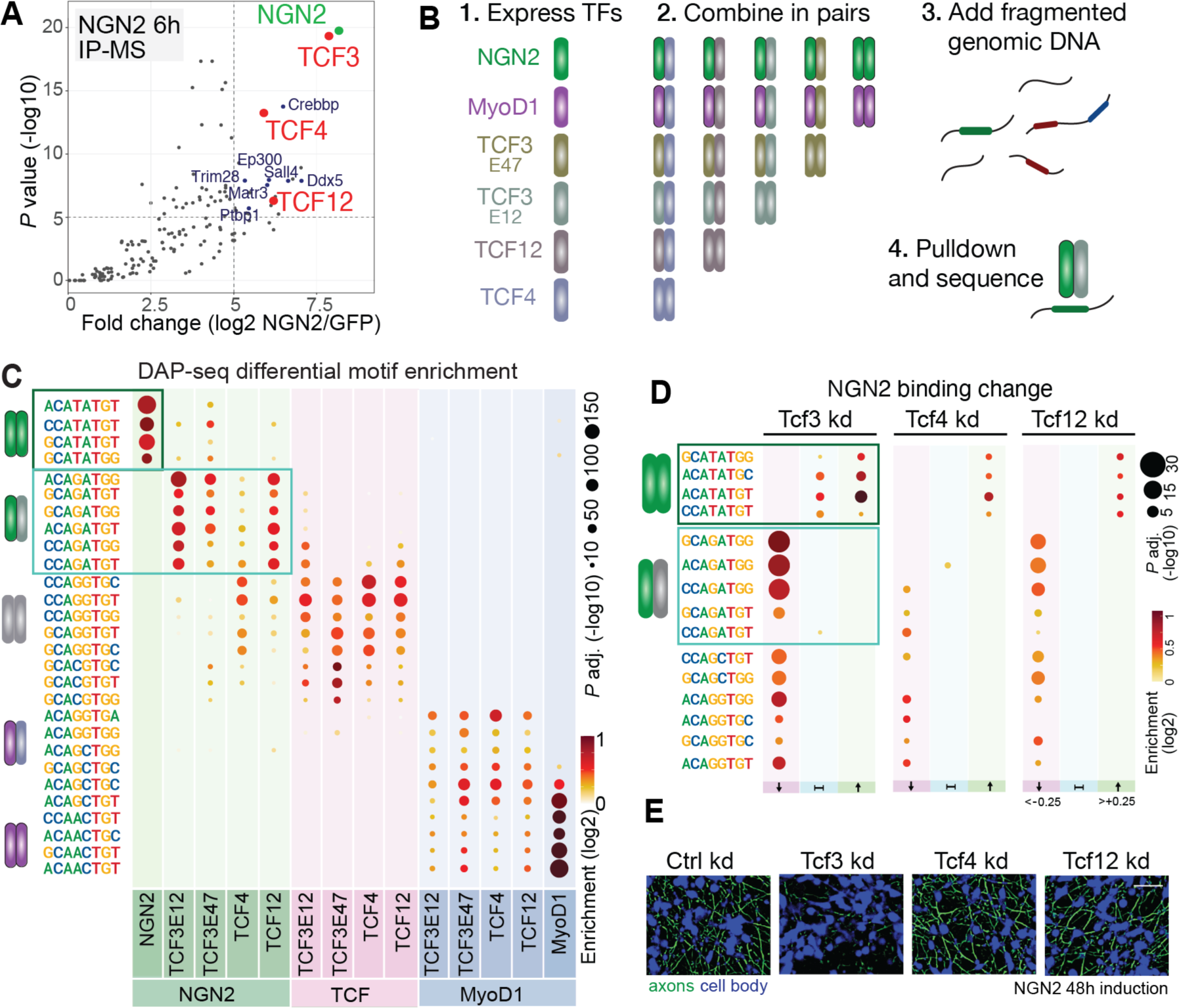
Homo-and hetero-dimer formation with other bHLH proteins increase the regulatory repertoire of NGN2 and MyoD1 **(A)** Mass spectrometric detection of proteins interacting with NGN2 (enrichment relative to GFP>5 with -log10(p value)>5). bHLH TFs are labeled in red, chromatin associated proteins in blue (Supplementary Table 8). **(B)** Scheme of the DAP-seq approach to probe genomic DNA binding preferences in vitro, testing NGN2 and MyoD1 alone and in combination with the putative bHLH partners: TCF3 (present as two main isoforms: E12 and E47), TCF4 and TCF12 **(C)** Differential E-box k-mer enrichment analysis (among top 5000 enriched genomic regions in DAP-seq) among NGN2, MyoD1 and TCF homodimers and heterodimers. **(D)** Differential E-box k-mer enrichment analysis among NGN2 ChIP-seq peaks grouped based on change in NGN2 binding at 6h upon decreasing TCF levels in comparison to control treatment. (kd: siRNA knockdown) **(E)** Images of 48h NGN2 induced cells for inspection of cellular morphology (cytoplasmic GFP signal segmented for axons in green and cell bodies in blue) treated with control (non-targeting siRNA), Tcf3, Tcf4 and Tcf12 siRNA. Scale bar, 50 µm.

To further test this hypothesis in the cellular context, we generated a tagged Tcf3 allele and profiled its genomic binding in mESCs which showed similar motif preferences to TCF homodimers **(Figure S5F-G)**. Subsequently, we decreased levels of TCF3, TCF4 or TCF12 prior to induction of NGN2 and measured the resulting impact on NGN2 binding **(Figure 5D, Figure S5H)**. Knock-down of TCFs reduced NGN2 binding specifically at the CAGATG motif, which is preferentially bound by the TCF-NGN2 heterodimers in vitro. However, it did not reduce NGN2 binding at the CATATG motif, which is preferentially bound by the NGN2 homodimer. In line with the in vitro binding, the effect differed depending on which TCF was reduced, whereby TCF3 had the highest impact on the CAGATGG motif, which is the most enriched motif for NGN2 induced neurogenic trajectory **(Figure S4J)**. The fact that decreasing cellular TCF3 levels causes failure of NGN2 induced neurogenesis further suggests that this dimerization is required for differentiation **(Figure 5E, Figure S5I).** Importantly, the observations on the initial binding strength and opening ability were in general correlated with these motif preferences of the protein partners **(Figure 4C** and **D, Figure S5E).**

Together, this demonstrates that different responses at motif variants reflect binding and activity of different sets of protein partners **(Figure S5J-K)**, by which single base-pair differences create significant regulatory diversity.

### Newly activated TFs reshape the accessibility landscape causing further divergence in NGN2 and MyoD1 binding patterns

While NGN2 and MyoD1 binding at the earliest timepoint is guided by chromatin their binding also modifies chromatin subsequently (**Figure 2-4**). To investigate whether their binding patterns further changes along the differentiation trajectory, we profiled binding and related chromatin features at later time-points. In addition to the factor-specific binding at initially less accessible regions, both NGN2 and MyoD1 gained new binding sites at 24 hours (24h). These are characterized by lower accessibility at earlier time-points yet become more accessible at 24h **(Figure 6A** and **B, Figure S6A and B)**. These new sites are specific to each factor, reflecting a divergence of the binding landscape that further reduced the overlap between MyoD1-bound and NGN2-bound sites. Contrasting the binding between 6h and 24h **(Figure 6C, Figure S6C and D)** revealed several motifs for transcription factors beyond NGN2 and MyoD1. For both factors, motifs specific to 6h were associated with the pluripotent cell state, such as ESRRB and POU(OCT)/SOX. At 24h however, EBF and ONECUT motifs characterized the newly formed NGN2 sites, whereas MEF2 and PITX/OTX motifs were enriched among newly formed MyoD1 sites. The TFs corresponding to these motifs exhibited correlated changes in their gene expression levels **(Figure S6E, Figure 6D, Figure S6I)**. For example, Ebf2 and Ebf3 genes were transcriptionally activated by NGN2. Contrasting the nuclear protein abundance between mESCs and NGN2 24h induction confirmed the increased presence of EBF2 and EBF3 as well as a decrease in pluripotency related TFs at 24h **(Figure S6F)**. In order to address how EBF2/3 affects NGN2 binding patterns, we compared the accessibility profiles of newly opened EBF sites that are either co-bound by NGN2 or remain unbound. Comparison of these genomic regions showed NGN2 binding and increased chromatin opening if there is an NGN2 motif within 100 bp distance around the EBF motif and thus below a nucleosomal repeat length **(Figure 6E** and **F, Figure S6G)**. In line with this, the CNN models trained for predicting the change in accessibility upon 24h of NGN2 and MyoD1 induction identified EBF and MEF/PBX motifs for NGN2 and MyoD1 respectively, where the gained binding occurred within the immediate range of accessibility created by these TFs **(Figure S6H, Figure S6H-L)**. This predicts that the motif syntax composed of both individual motif sequences and their relative positions impacts these TFs’ activity on chromatin.

**Figure 6.**
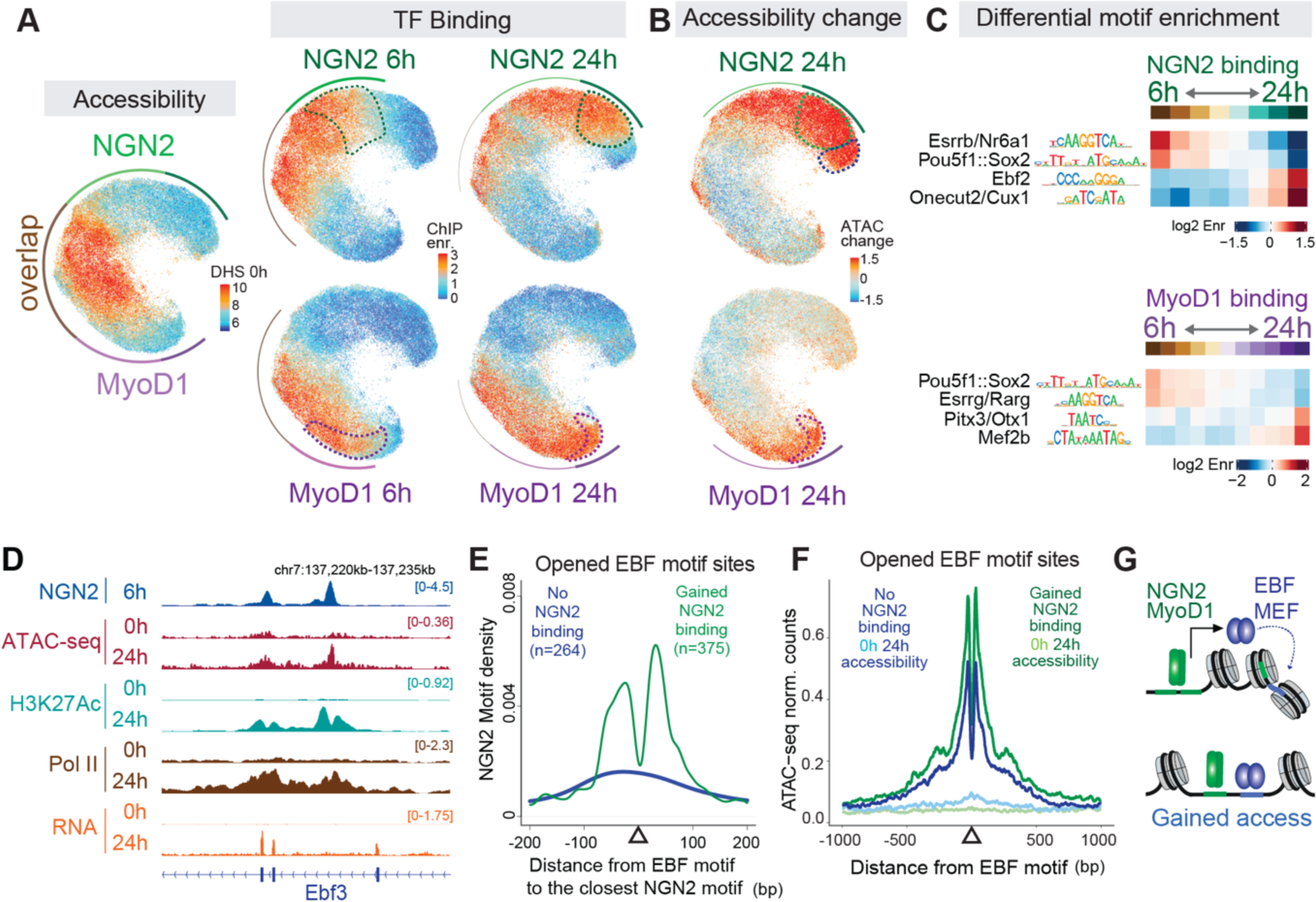
Reshaped accessibility landscape diverges NGN2 and MyoD1 genomic binding **(A)** UMAP of genomic regions (as in FIGURE 4A) displaying initial accessibility, NGN2 and MyoD1 binding at 6h and 24h. Dashed lines outline NGN2 and MyoD1 specific regions at 6h and new regions at 24h. The decrease in binding signal at the overlapping regions (higher 0h DHS) and gain in new regions (lower 0h DHS) show divergence of the binding patterns of NGN2 and MyoD1 through differentiation (h: hours). **(B)** Accessibility changes (ATAC-seq fold-change) upon 24h NGN2 and MyoD1 induction, displayed on the UMAP, show loss at overlapping high DHS regions and gain in new low DHS regions. Outlines indicate gained NGN2 binding sites at 24h in green, gained MyoD1 binding sites at 24h in purple and sites that gain accessibility but not NGN2 binding are outlined in blue. **(C)** Top two enriched unique motifs identified in differential motif enrichment analysis (JASPAR TF catalog) contrasting 6h vs 24h NGN2 and MyoD1 binding. Combined peaks of 6h and 24h are grouped according to changes in binding enrichment between 6h and 24h (FigureS6C-D). **(D)** Genome browser tracks of NGN2 ChIP-seq, ATAC-seq, H3K27Ac ChIP-seq, Pol2 ChIP-seq and RNA-seq at the Ebf3 locus. **(E)** NGN2 motif density around EBF motifs that are newly made accessible upon 24h NGN2 induction, sub-grouped as NGN2 bound and not bound at 24h (corresponding to the outlined regions in B, green vs blue). **(F)** Metaplots showing ATAC-seq signal pre and post NGN2 induction at 24h across genomic regions grouped as in (E). The increase in ATAC-seq signal shows gained accessibility in regions with an EBF motif upon induction, and the regions with an EBF motif and NGN2 motif (within 100 bp distance) gain NGN2 binding and even higher accessibility. **(G)** Schematic summary of dynamic NGN2/MyoD1 binding through differentiation.

We conclude that NGN2 and MyoD1 engagement to initially closed sites at 6h activates expression of other TFs. These TFs subsequently change the chromatin landscape, creating accessibility that generates new opportunities for binding by NGN2 and MyoD1, further diverging the activity patterns in a lineage specific fashion **(Figure 6G)**.

### The CNN derived chromatin-dependent motif syntax is transferable and predicts TF-chromatin interactions in different cellular contexts

Since the observed binding patterns for NGN2 and MyoD1 are dependent on the initial genomic accessibility, which itself reflects the activity of other TFs, we expect them to vary between cell types. If our predictive model for binding is generalizable, it should be transferable to different cell types as long as their starting accessibility is known. We tested this hypothesis using available ChIP-seq and ATAC-seq datasets from a widely used myogenesis model, C2C12 cells, where MyoD1 can be induced by changing culture conditions ^46–48^. Analyzing MyoD1 binding in this system identifies the same preferred binding motifs as in our cellular system, however the actual sites of MyoD1 binding in C2C12 cells were largely different from mESCs, with only 12.5% overlap among consensus binding sites (**Figure 7A-C, Figure S7A and B**). Differentially bound regions indeed had high accessibility in only one of the cell types whereas overlapping regions had varying degrees of starting accessibility.

**Figure 7.**
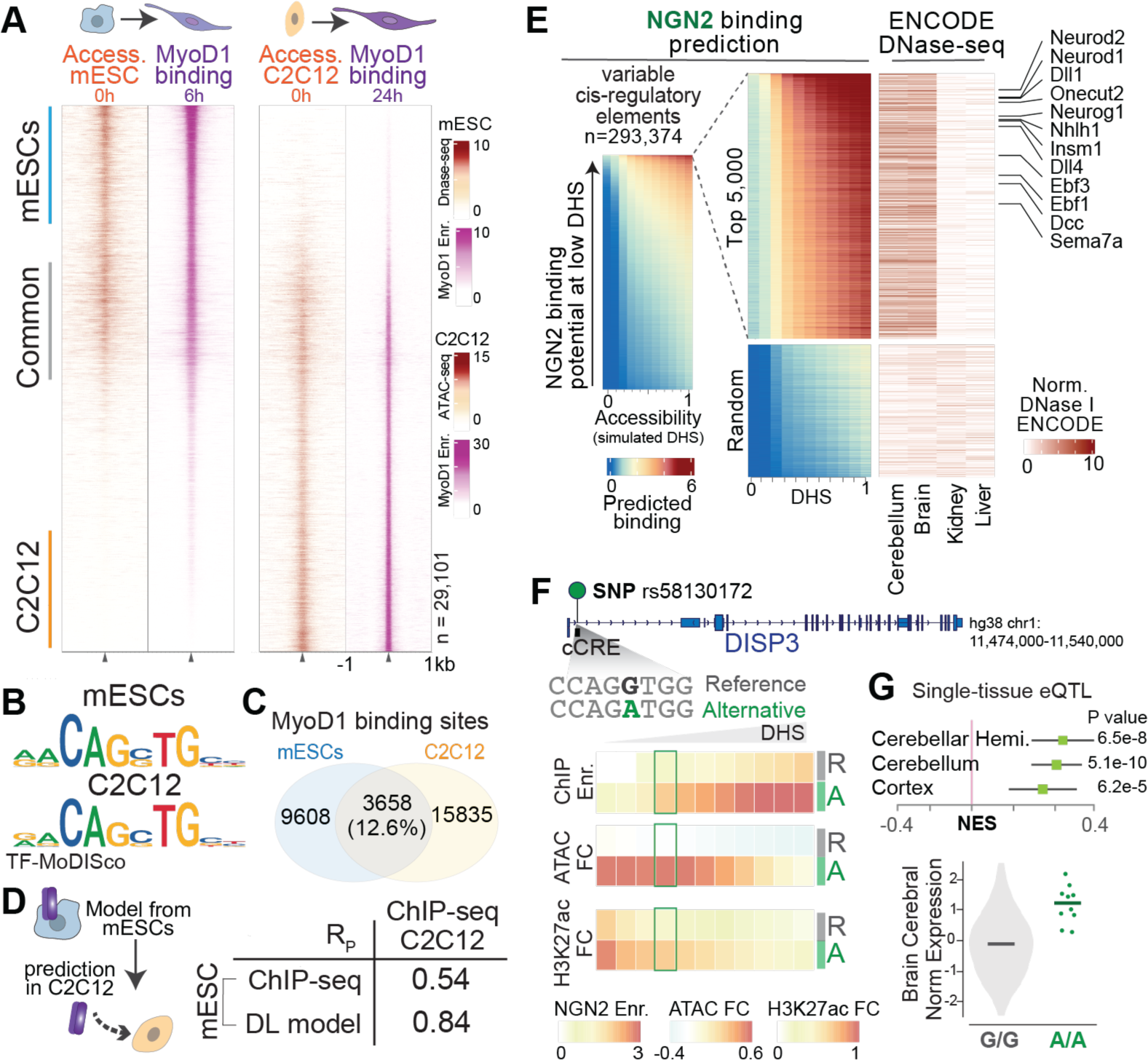
The derived model predicts TF binding and activity in other cellular contexts **(A)** Heatmap of MyoD1 ChIP-seq in mESCs (6h) and C2C12 (24h) cells upon induction, visualized on merged set of MyoD1 binding sites from both cell types, together with their pre-induction accessibility in mESCs (DNase-seq) and in C2C12 cells (ATAC-seq). **(B)** Enriched E-box seqlets identified in MyoD1 bound sites for mESCs and C2C12 cells, generated by TF-MoDISco. **(C)** Venn diagram contrasting MyoD1 ChIP-seq peaks in mESCs and C2C12 cells that show less than 15% overlap among combined binding sites between both cell types. **(D)** Scheme describing the CNN model transfer that is trained on mESCs and applied for predictions in another cell type (left panel). Correlation of the observed MyoD1 ChIP-seq enrichment in C2C12 cells with the observed enrichment values in mESCs versus the predicted enrichment values by the mESC trained CNN model (right panel). **(E)** Predicted NGN2 binding enrichment in all mouse ENCODE cCREs (rows) as a function of simulated initial DNA accessibility (columns), sorted based on NGN2 binding potential at low DHS (left panel). Top 5000 tissue specific cCREs with highest NGN2 binding potential at low DHS contrasted with 2000 randomly sampled tissue specific cCREs from the bottom 50% of the left panel (middle panel). DNase I accessibility counts corresponding to the cCREs in the middle panel, showing top two enriched and two closest to background tissue profiles available in ENCODE datasets (right panel). The specific DNase-seq profiles indicate that the cCREs with high NGN2 binding potential gain activity in cerebellum and brain samples. Example CREs proximal to the NGN2 upregulated genes, observed in mESC to neuronal differentiation model, are labeled. **(F)** rs58130172 as an example GWAS defined SNP, modifying an E-box variant that resides in a tissue specific human cCRE active in human brain. Scheme describing the SNP location at DISP3 gene and the effected sequence (top panel). Predicted NGN2 binding and activity change at the reference and alternative CRE sequence as a function of initial accessibility (bottom panel). **(G)** Measured effect of the rs58130172 SNP on DISP3 gene expression retrieved from eQTL data.

Despite the low similarity in genomic binding between the two cell types, the CNN model, trained only on mESCs, accurately predicts the binding in C2C12 cells using as input the accessibility in these cells prior to MyoD induction. The model performs comparably to its performance in mESCs (**Figure 7D, Figure S3A, Figure S7C**) illustrating that the rules of MyoD1 binding learned in mESC can readily be transferred to other cell states with their unique accessibility landscape.

At sites where NGN2 or MyoD1 can open chromatin, our model predicts a motif syntax that is less dependent on the existing accessibility for binding. Therefore, genomic sequences bearing such syntax are expected to be targeted in all cell types where the respective TF is expressed. To test this, we took a large set of cis-regulatory elements (CREs) identified within ENCODE consortium ^49^(see Methods) and used the model to predict binding for a range of different DNA accessibility values for each region **(Figure 7E, Figure S7D).** Top ranking regions that provide high NGN2 binding potential at low accessibility in this simulation indeed show accessibility in tissues (i.e. brain and cerebellum) where NGN2 is expressed. These regions included CREs of genes known to be critical for neuronal differentiation, which we also observed as being activated during the induced differentiation of mESCs arguing that this is a developmentally conserved feature. A comparable analysis applied to MyoD1 revealed its activity in muscle, but also its established but less well-known activity in thymus development ^50,51^.

Given its high predictability we wondered if the CNN model could guide the interpretation of sequence variations that are linked to gene expression variation as eQTLs^52^ reported in GWAS studies. To test this, we predicted binding of NGN2 and its impact at SNPs that modify an NGN2 motif variant in a tissue specific CRE that is detectable in the respective tissues (i.e. cerebellum, brain) **(Figure 7E).** This is exemplified by the variant rs58130172 (GWAS: GCST90105038 ^53^), at the DISP3 locus, a gene reported to influence neural fate decisions in a dose-sensitive manner ^54^ **(Figure 7F).** While the predictions on the reference enhancer sequence foresee low binding with no activation potential, the alternative sequence with the SNP displayed higher expected binding and activation. Consistent with the prediction, this variant had been identified as an eQTL and the alternative allele showed increased DISP3 expression in the brain (**Figure 7G)**.

Taken together, these results show that the identified chromatin-and sequence-dependent motif syntax is transferrable to other cellular origins, providing predictive power and deepening our understanding of how TFs that recognize such short abundant motifs can achieve specific regulatory outcomes and drive cellular differentiation.

## DISCUSSION

Here we demonstrate how distinct DNA sequence syntax, chromatin accessibility, and interaction partners accounts for shared and factor specific binding of two transcription factors with distinct lineage driving potential, despite operating on seemingly similar cognate motifs of low sequence complexity. This was made possible by monitoring chromatin landscape before and following TF induction combined with measurements of binding, transcriptional response, factor dimerization and in vitro DNA binding. The analysis of these datasets combined with convolutional neuronal networks generated a highly predictive model of TF binding and activity that is broadly applicable across cellular contexts.

Previous approaches mostly used chromatin accessibility to explain binding in a cell state where the TF under study is already expressed, thus mainly relying on the correlation between binding and local chromatin state. Our study illustrates the benefits of incorporating the chromatin landscape prior to TF presence and measuring acutely after TF induction. This enables the careful dissection of where these factors bind opportunistically in response to open chromatin, where they work cooperatively, and where they open chromatin themselves, acting as pioneer factors. This uncovered a TF-specific syntax that can readily predict binding and activity as a function of DNA sequence and preexisting accessibility (Figure 3-4).

Most eukaryotic TFs recognize short motifs and bind only to a minor fraction of potential genomics sites. It is generally accepted that the majority of these sites are masked by chromatin, consequently predicting the preferential binding to open chromatin, as seen here for many sites bound by NGN2 and MyoD1. We observe that the engagement with accessible sites is largely overlapping between both factors, though it remains to be sequence dependent, i.e. requiring E-box motifs (Figure 2). This type of opportunistic binding depends on the cell-type-specific accessibility landscape which is also exemplified by the changes observed during further lineage transitions (Figure 6). Newly activated TFs promote chromatin opening, which in turn creates new binding opportunities for both MyoD1 and NGN2. Since the newly activated TFs are lineage specific, the gained binding sites by NGN2 and MyoD1 can be assumed to contribute to the observed differentiation trajectories instead of being a mere by-product of other TFs’ activity. This model of combinatorial action would readily account for the frequently observed cell-type dependent binding patterns of many mammalian TFs, that are chromatin sensitive. It also explains how these factors function in very distinct cell types as illustrated by the fact that Neurogenin/Neurod/Onecut together regulate neural, pancreatic and intestinal development, and similarly MyoD1/Myogenin/Mef2 regulate muscle as well as thymus development ^51,55^.

In this context, it is important to note that the majority of motifs residing in open chromatin regions remain unbound. Nevertheless, binding is correctly predicted by the CNN model, likely because the accessibility measure provides local information on the nucleosome presence and position ^37^. Since bHLH TFs are not able to engage with motifs embedded in the nucleosome core in vitro ^33^, it seems plausible that NGN2 and MyoD1 are sensitive to positioned nucleosomes in vivo (Figure 3G). Such nucleosome sensitivity could also explain why opportunistic TF binding adjacent to another TF motif occurs within a variable but limited distance.

Given the strong dependence on local chromatin accessibility, it seems counterintuitive and unexpected that we can locate genomic sites where MyoD1 or NGN2 bind to inaccessible sites and cause chromatin opening. This suggests that they can also act as “pioneer factors”, which contrasts with their chromatin sensitivity. However, such activity is largely limited to sites characterized by multiple occurrences of TF-specific motif variants. This results in binding and chromatin opening at a complex composite motif syntax that occurs rarely in a genome by a TF, which normally recognizes a low complexity and highly abundant motif (Figure 2-4). This model of chromatin-insensitive binding at a composite-motif predicts that these sites should be commonly targeted in all cell-types that express the respective factor, which we indeed observe for MyoD1 in C2C12 cells and for NGN2 in brain tissues (Figure 7). Furthermore, the related gene targets (e.g. Neurod, Onecut, Ebf) exhibit specificity for the respective transcription factors, when enriched exclusively with their preferred TF motif variants (Figure 6, Figure 7E).

Monitoring the initial binding’s impact using CNN model further identified clear differences between motifs regarding their activation potential. In general, stronger binding correlates with higher activity, as expected, but there are numerous cases where activation and binding potential linked to a motif variant differ (Figure 4).

Although “context dependent activity” of TFs is commonly observed, it mostly remains molecularly undefined. Here we show that these slight motif variations lead to differential activation of a chromatinized reporter and thus act autonomously. We further link these to the in vitro binding preferences of different bHLH homo- and hetero-dimers, with resulting functional predictions readily validated by TF depletion, as evidenced for TCF3 (Figure 5). In addition to the direct regulation by NGN2 and MyoD1, we observed changes in motif variant regulation over time, which correlated with the presence of downstream activated E-box binders such as NEUROD1^56^, ZEB2^57^, ZBTB18^58,59^ and MYOGENIN^60^ (Figure S6C-F and H), showcasing the complicated functional regulatory diversity achieved through short motifs, which however can be traced with the applied approach. The combination of comprehensive modeling of epigenomic profiles with in vitro and in vivo validations used in this study can thus approximate, in unprecedented detail, how members of a large TF family (i.e. bHLH) with similar motifs can define specific differentiation trajectories. It further provides a rationale of how preferential interactions operating on single base changes increase the regulatory diversity, allowing targets to be controlled with distinct dynamics.

Combined, our findings expose the expected complexity of genomic binding and lineage driving potential of short motif binders such as MyoD1 and NGN2. Despite its complexity, we demonstrate a roadmap showing how binding behavior and regulatory impact can be decoded and integrated into highly predictive models.

These models largely rely on preexisting chromatin accessibility, co-expressed TFs, and DNA sequence, all of which can be precisely measured using current techniques at the cellular and tissue level. According to the resulting model, binding to less complex and more frequent motifs depends on cooperativity with other TFs and thus is largely cell type specific. In contrast, sites with a composite motif syntax are rare in the genome but are bound and critical for the lineage driving potential. This further exemplifies the breadth of chromatin sensitivities of individual transcription factors functioning through both opportunistic and deterministic binding events, which collectively drive specific genetic and epigenetic programs for cellular differentiation.

## Contributions

S.D. and D.S. designed the study, interpreted the results and wrote the manuscript with input from the co-authors. S.D. performed the experiments. S.D. and M.I. analyzed the data. M.I. established the CNN model, tools for its interpretability and performed the predictions with input from S.D.. L.I. established the DAP-seq protocol with input from S.D.. L.H. assisted in experiments with cloning and cell-line generation. C.W. generated HA36Tet3G parental cell-lines. L.B. contributed to data analysis. V.I. performed TMT labeling and mass-spectrometry for the nuclear proteome and D.H. performed the mass spectrometry for affinity purification mass-spectrometry.

## Acknowledgements

We would like to acknowledge FMI Facilities for excellent technical support, especially Sebastian Smallwood, Sirisha Aluri and Stephane Thiry (Functional Genomics), Hubertus Kohler (Cell Sorting), Laurent Gelman (Microscopy and Imaging) and Jan Seebacher (Protein Analysis). We would like to acknowledge Ralph Grand for open discussions throughout the study, Michael Stadler for his input on differential motif discovery and feedback on the manuscript, and Luca Giorgetti, Lisa Baumgartner, Ana Petracovici and Francesca Masoni for feedback on the manuscript.

D.S. acknowledges support from the Novartis Research Foundation, the Swiss National Science Foundation (310030B_176394 to D.S.) and the European Research Council under the European Union’s (EU) Horizon 2020 research and innovation programme grant agreements (ReadMe-667951 and DNAaccess-884664). S.D. acknowledges support from the European Molecular Biology Organization Long-Term fellowship (ALTF 1101) and the Christiane-Nüsslein-Volhard Stiftung. L.I. acknowledges the National Health and Medical Research Council CJ Martin Fellowship (APP1148380) and EU Horizon 2020 Research and Innovation Program under the Marie Sklodowska-Curie grant (748760).

The funders had no role in study design, data collection and analysis, decision to publish or preparation of the manuscript.

## SUPPLEMENTAL FIGURES

**Figure S1.**
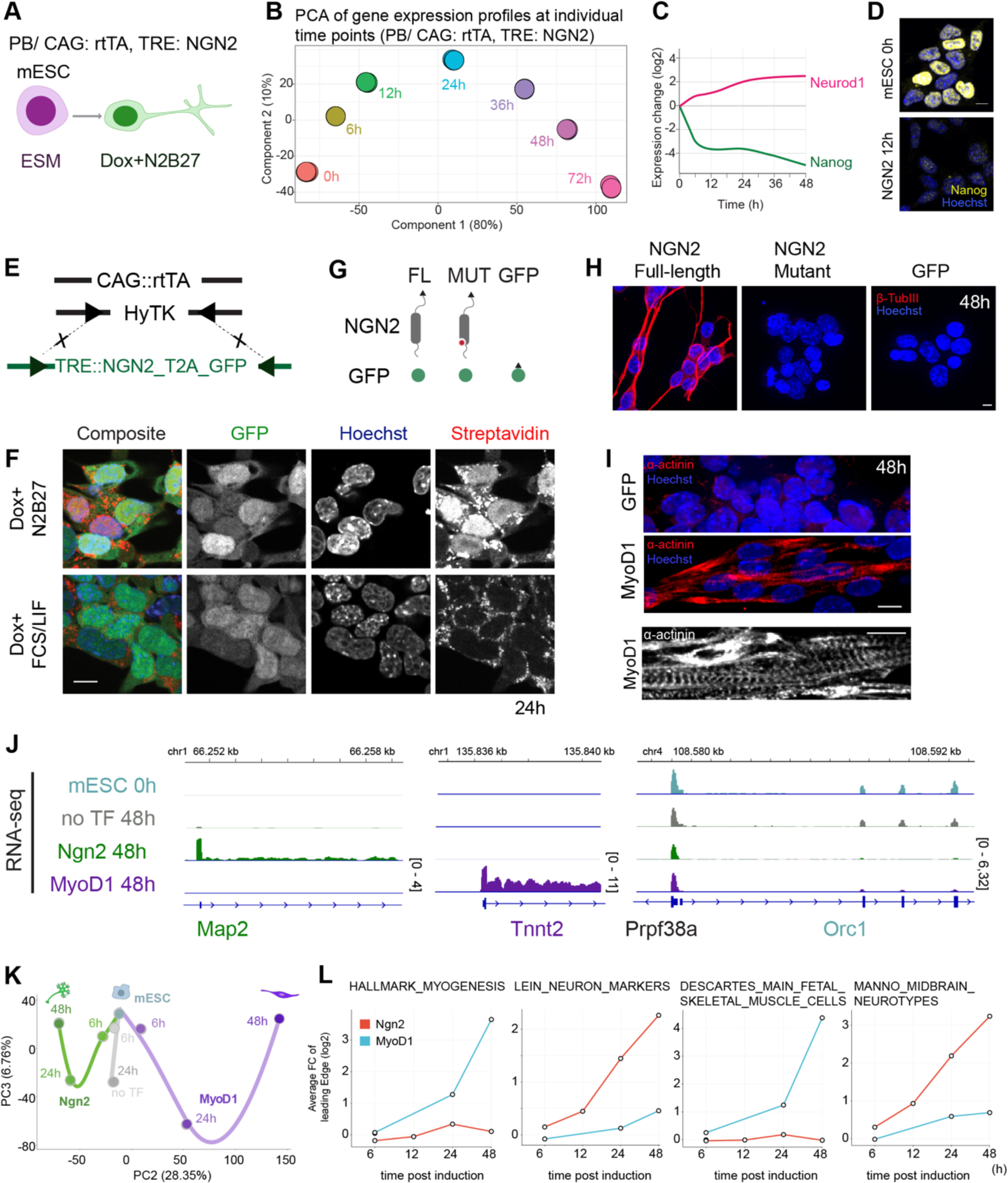
Characterization of NGN2 and MyoD1 induced differentiation, related to Figure 1 **(A)** Scheme of neurogenesis by NGN2 induction, utilizing mESCs with multiple, random integration of a piggyback CAG:rtTA/tetO:NGN2 construct. **(B)** Principal component analysis (PCA) of mRNA-seq from four replicates of NGN2 induction at consecutive time-points. **(C)** Gene expression change of neuronal marker Neurod1 and pluripotency marker Nanog upon NGN2 induced differentiation. **(D)** Immunofluorescence staining for NANOG (yellow) in mESCs and 12 hours post NGN2 induction, illustrating its depletion from nuclei (Hoechst, blue). Scale bar, 10 µm. **(E)** Description of separate, single-locus integration of CAG:rtTA and tetO:ORF, facilitating stable, controlled and comparable gene expression. **(F)** Immunofluorescence images reporting nuclear NGN2 protein levels (streptavidin staining for NGN2 in red, Hoechst staining for nuclei in blue) post-doxycycline induction in two different media: mESC (FCS/LIF) and N2B27 (w/o VitA), noting that while gene expression is induced comparably (as reported by t2a-GFP in green), NGN2 protein is degraded in mESC media, necessitating the removal of pluripotency signals for robust NGN2 induction and cellular differentiation. Scale bar, 10 µm. **(G)** Description of constructs expressed upon doxycycline induction: full-length TF and E-box binding mutant TF followed by t2a-GFP expression reporter, and GFP expression alone serving as noTF induction control. **(H)** Immunofluorescence staining of TUBB3 neuronal marker (red) at 48 hours post NGN2 induction, NGN2 E-box binding mutant, and GFP induction. Nuclei/Hoechst in blue. Scale bar, 10 µm. **(I)** Upper panels: α-Actinin myocyte marker staining (red) at 48 hours post-induction of MyoD1 and GFP. Bottom panel: Single-plane slice of α-Actinin staining (white) in differentiated myocytes at 48 hours post MyoD1 induction, demonstrating typical muscle sarcomere structure. Nuclei/Hoechst in blue. Scale bar, 10 µm. **(J)** Genome browser tracks showing total RNA-seq of mESC, noTF, NGN2, and MyoD1 at 48 hours post-induction at neural (Map2), myocyte (Tnnt2), and cell proliferation (Orc1) marker genes, exemplifying up-regulation of cell-type specific genes upon differentiation and common loss of cell proliferation. **(K)** Principal component 2 and 3 of the PCA analysis in Figure 1B for total RNA-seq data at consecutive time-points, demonstrating the differentiation trajectories of NGN2 induction in green, MyoD1 in purple, no-TF control induction in gray and mESCs in blue. **(L)** Gene-set enrichment analysis of differentially regulated genes between mESC and NGN2 or MyoD1 induction at multiple time-points, illustrating the cell-type specification of neurogenesis or myogenesis respectively.

**Figure S2.**
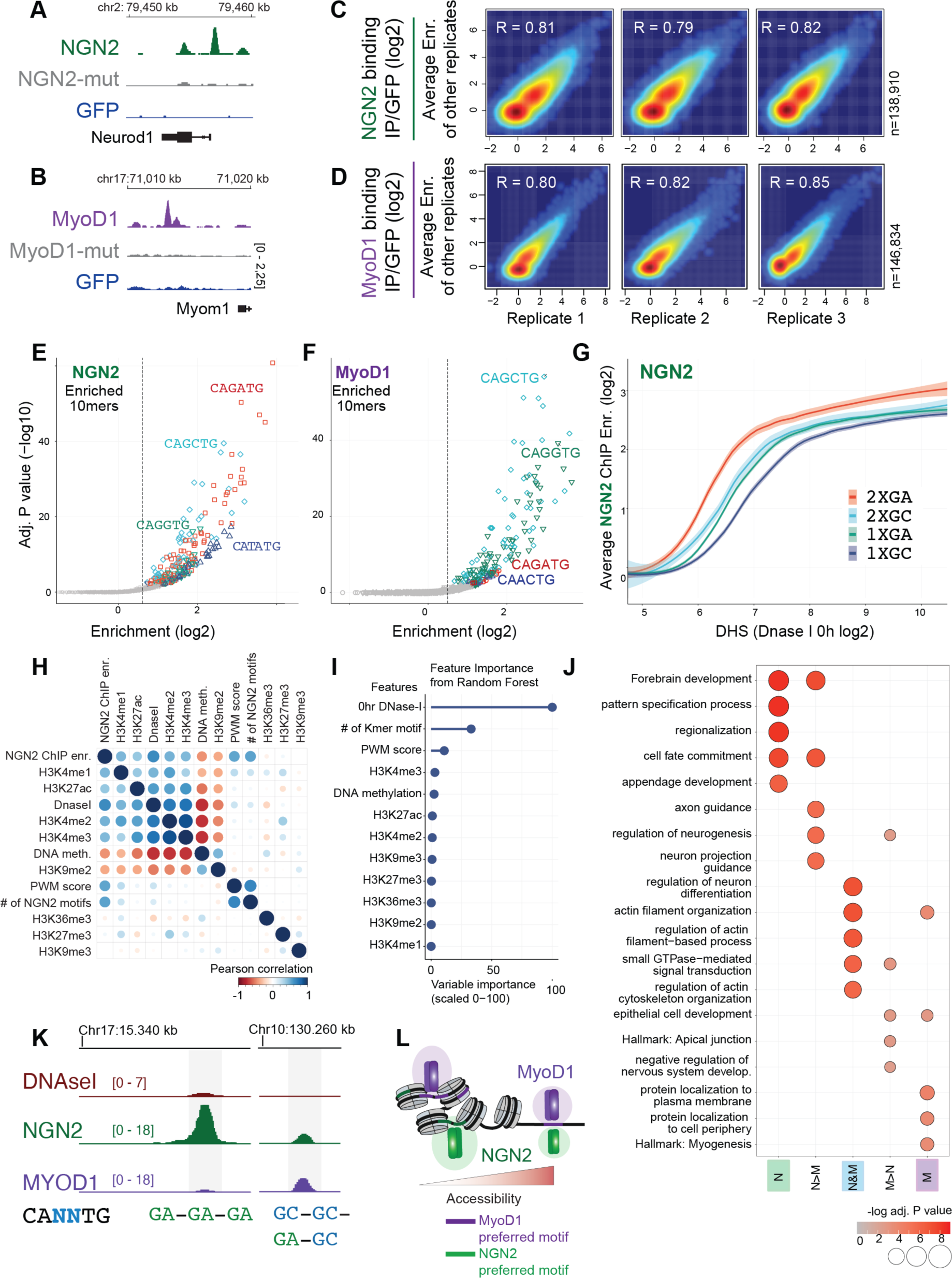
Characterization of NGN2 and MyoD1 binding to mESC chromatin, related to Figure 2 **(A** and **B)** Genome browser tracks of NGN2 and MyoD1 ChIP-seq with corresponding DNA binding mutants and GFP as ChIP-seq controls at known target genes: Neurod1 and Myom1, respectively. **(C** and **D)** ChIP-seq enrichments of NGN2 and MyoD1 replicates (6h) across MACS2-called peaks and the same number of random background regions. **(E** and **F)** K-mer (10bp, NNCANNTGNN) enrichment analysis on consensus peaks (14116 NGN2, 13266 MyoD1) using monaLisa for NGN2 and MyoD1 cognate motif sequences. E-box k-mers are colored according to their central nucleotide variations of the E-box sequence (GA, TA, GC, GG, AC). **(G)** Fitted line plot of average NGN2 ChIP-seq enrichment versus initial DNA accessibility in NGN2 consensus peaks with matched background regions grouped for motif number and motif variant composition (sites with one GA motif, with one GC motif, with two GA motifs, with two GC motifs), illustrating the difference in binding potentials depending on motif variant, motif number and prior accessibility. **(H)** Correlations of chromatin marks (DNase-seq, ATAC-seq, H3K4me1, H3K4me2, H3K4me3, H3K27ac, H3K27me3, H3K9me2 ChIP-seq log2 enrichment of IP/input) and NGN2 binding (NGN2 ChIP-seq log2 enrichment of IP/GFP and number of NGN2 motifs). **(I)** Feature importance of random forest prediction model for NGN2 binding across NGN2 peaks and randomly sampled genomic regions, using the occurrences of NGN2 motifs and chromatin features (accessibility and histone marks). **(J)** GO enrichment analysis on genes, corresponding to the peaks (linked by closest distance to the TSS) grouped as in Figure 2I based on NGN2 versus MyoD1 binding. **(K)** Example genomic regions of N>M and N&M at low accessibility, as extension of Figure 1J. **(L)** Schematic model of preferential NGN2 and MyoD1 binding patterns, showing the effect of chromatin barrier, motif preference and multi motif occurrence.

**Figure S3.**
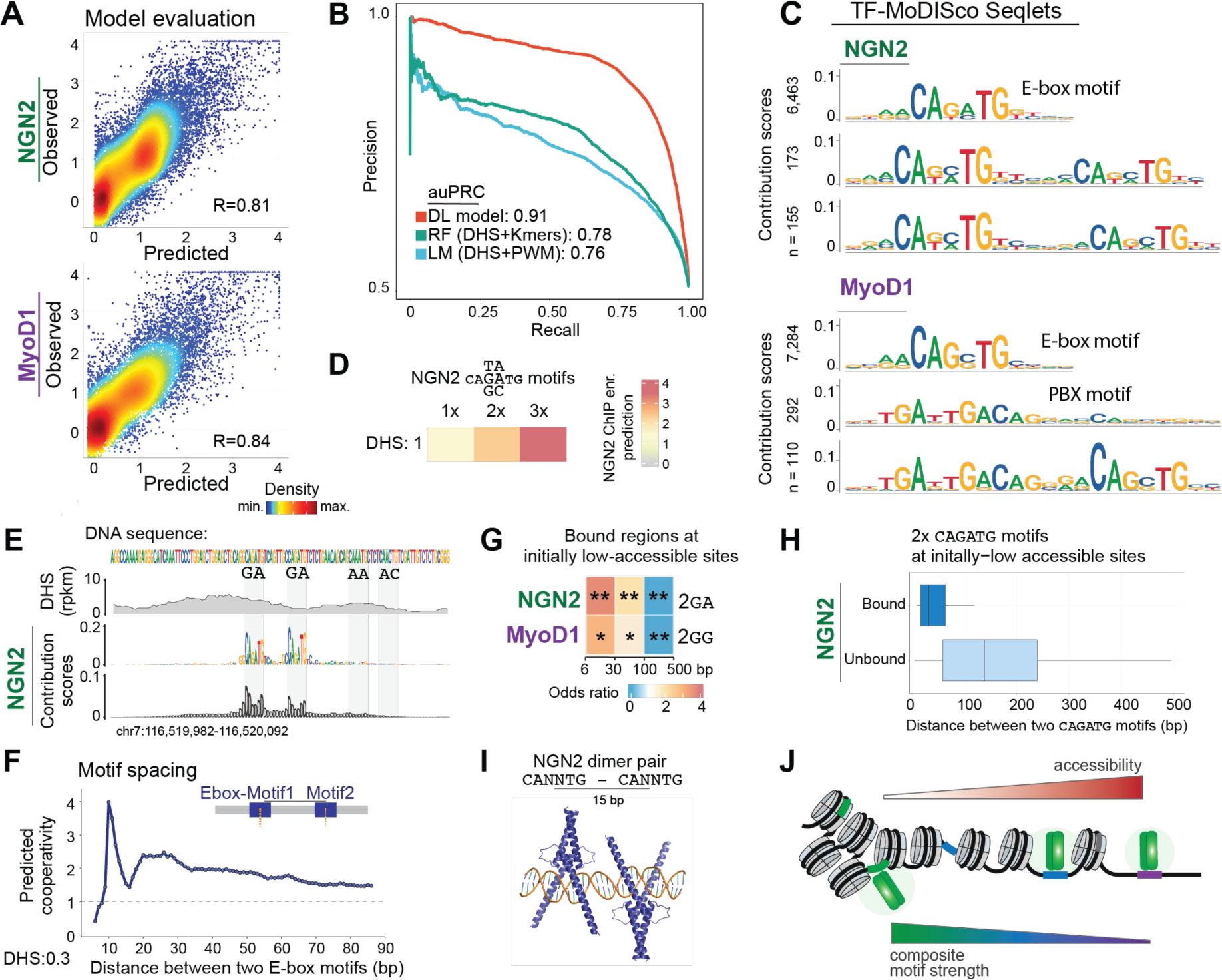
CNN model predicts ChIP-seq enrichments, underlined by a chromatin-dependent motif syntax, related to Figure 3 **(A)** Evaluation of the CNN model on the held-out test set (regions that the model was not trained on) by comparing measured versus predicted signal. **(B)** Precision recall calculation for evaluating the prediction of NGN2 binding across genomic sites with comparable initial accessibility distributions using DL model (DHS and DNA sequence), random forest (DHS and enriched k-mers) or linear model (DHS and enriched PWMs). **(C)** Three seqlets from TF-MoDISco results for NGN2 and MyoD1 binding predictions that contains E-box motif(s). E-box motifs detected as singles, in pairs or in combination with an interacting motif such as PBX for MyoD1. Numbers represent the occurrence of the seqlets with high contribution scores in the consensus peaks. **(D)** The effect of number of motifs on the predicted NGN2 binding strength is illustrated for NGN2 motifs using synthetic sequences. **(E)** Single-locus example of NGN2 bound regions. Upper panels visualize DNA sequence and measured accessibility, that are given as inputs to the model. Lower panels show their importance per base for the NGN2 binding prediction. **(F)** Predicted cooperativity of two NGN2 motifs at low DHS for enhancing binding potential across various distances between them (bp measured center to center). **(G)** Odds ratio for presence of the second NGN2 or MyoD1 motif at certain distances among bound low DHS regions with two motifs. **(H)** Distance distribution between motifs in NGN2 bound versus unbound regions with low initial accessibility and two CAGATG motifs. **(I)** Modeled NGN2 binding (AlphaFold 3) on two NGN2 motifs with 15 bp center to center distance, which gives the highest binding potential. **(J)** Model summary, highlighting the composite motif strength and its accessibility as the principal determinants of TF binding patterns.

**Figure S4.**
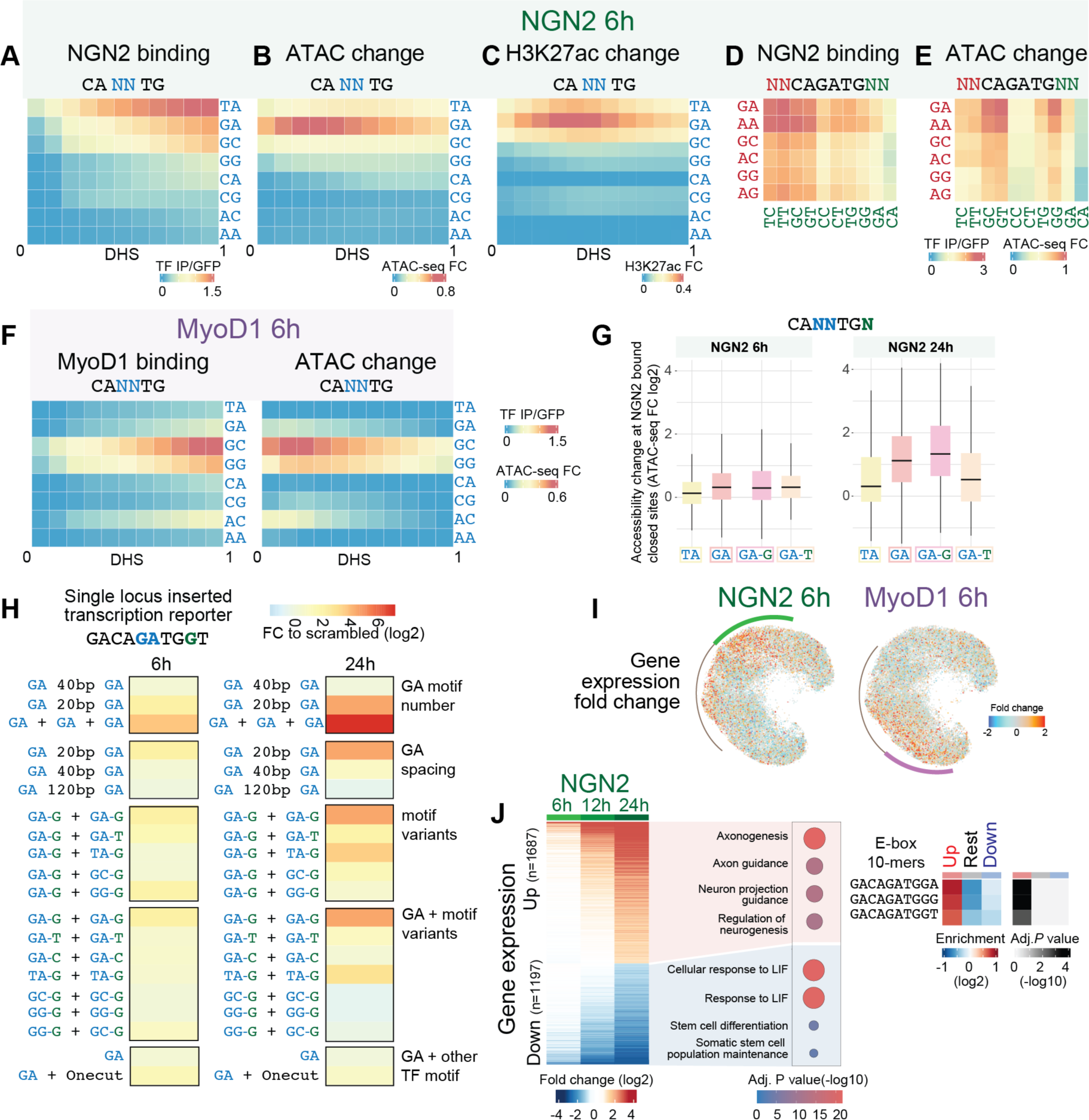
Impact of NGN2 and MyoD1 binding on chromatin and gene expression, related to Figure 4 **(A**-**C).** Predicted binding, accessibility (ATAC-seq) change and H2K27Ac ChIP-seq change upon NGN2 induction (6h) at NGN2 motif variants depending on simulated initial accessibility. **(D** and **E)** Predicted binding and accessibility change upon 6h NGN2 induction for lowly accessible (DHS 0.3) CAGATG motif depending on the flanking-bases **(F)** Predicted binding and accessibility (ATAC-seq) change upon MyoD1 induction (6h) at MyoD1 E-box motif variants depending on simulated initial accessibility. **(G)** Boxplots of measured ATAC-seq change (6h and 24h) at NGN2 peaks with low initial accessibility and indicated motif composition (TA: peaks that contain TA motif variant but not GA, GA: peaks that contain GA motif variant but not TA, GA-G: subset of GA, with three prime G flanking nucleotide, GA-T: subset of GA, with three prime T flanking nucleotide). **(H)** Single locus inserted stable transcription reporters designed for measuring the effect of NGN2 motif variations on gene expression, as described in Figure 4F (Supplementary Table 5). **(I)** Expression change of genes assigned based on proximity to the genomic sites displayed on the UMAP (as in Figure 4A) upon 6h NGN2 and MyoD1 induction. **(J)** Heatmap of expression changes of differentially expressed genes (absolute FC>0.5, p.val<0.001) upon NGN2 induction (left panel) and their top enriched GSEA terms (middle panel). Differential motif enrichment analysis (E-box 10mers, Enr>0.5, adj.p<0.01) among NGN2 peaks assigned to the group of genes that were consistently up-regulated, down-regulated and the rest. The G flanking CAGATG motif is enriched for continuously up-regulated genes (right panel). In (A-H), central nucleotides of the E-box motif are colored in blue, flanking nucleotides in red for 5’, in green for 3’.

**Figure S5.**
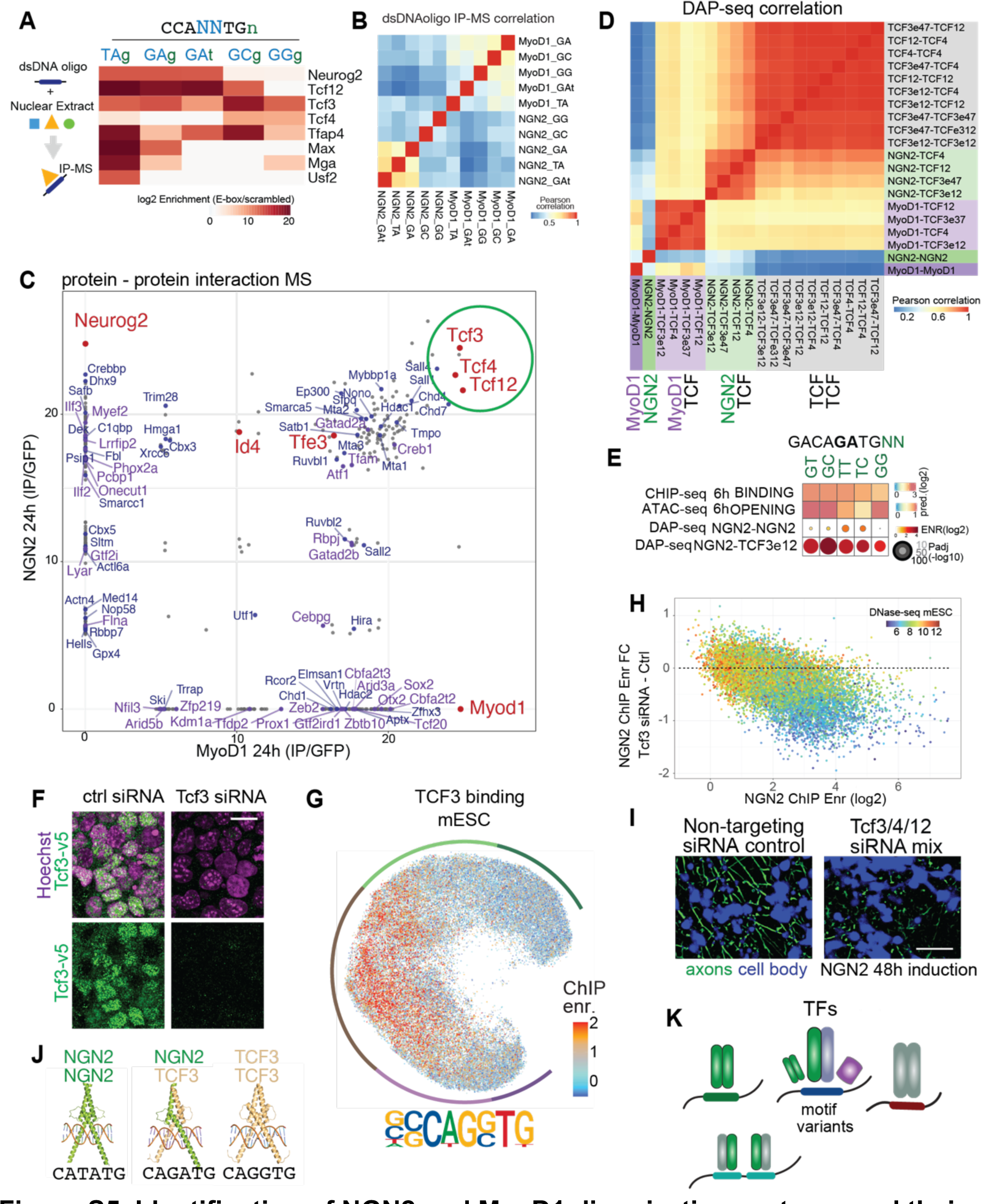
Identification of NGN2 and MyoD1 dimerization partners and their relevance on the differential activity observed at distinct motif variants, related to Figure 5 **(A)** Mass spectrometric detection of proteins interacting with E-box variants using synthesized double-stranded DNA-oligomers as bait in nuclear extracts of TF induced cells. Enrichment of E-box binding TFs relative to control oligos in NGN2 24h nuclear extracts (Supplementary Table 10). **(B)** Correlation heatmap of DNA oligo pull-down mass-spectrometry data in NGN2 and MyoD1 nuclear extracts, with motif variants labeled as in Figure S5A. **(C)** Mass spectrometry detection of protein interaction partners of NGN2 and MyoD1 at 24h. bHLH domain containing proteins are labeled in red, TFs with sequence specific DNA binding activity are labeled in purple and other proteins with chromatin associated functions are labeled in blue (Supplementary Table 9). **(D)** Correlation heatmap of DAP-seq samples, described in Figure 5B. **(E)** Binding versus opening at CAGATG motif flanking with G or T (predicted NGN2 6h ChIP-seq enrichment and NGN2 6h ATAC-seq fold change as in Figure 4C-D), contrasted with the motif enrichments in NGN2-NGN2 DAP-seq peaks and NGN2- TCF3E12 DAP-seq peaks against genome. **(F)** Immunofluorescence images of endogenously tagged Tcf3-v5 mESCs upon Tcf3 and non-targeting siRNA treatments, showing depletion of TCF3-V5 protein for testing the V5 tagging and response to knock-down treatment. Scale bar, 20 µm. **(G)** Top enriched motif in V5 tagged TCF3 ChIP-seq peaks in mESCs (top) and TCF3 ChIP-seq enrichments displayed on the UMAP (as in Figure 4A) **(H)** Scatter plot of NGN2 ChIP-seq enrichments in untreated cells (x axis) and NGN2 binding (6h) change upon decreasing TCF3 levels in mESCs in contrast to control treatment (y axis), color coded based on untreated mESC DNase-seq (DHS) signal. **(I)** Images of 48h NGN2 induced cells for inspection of cellular morphology (cytoplasmic GFP signal segmented for axons in green and cell bodies in blue) treated with control (non-targeting siRNA) and mix of Tcf3, Tcf4, Tcf12 siRNA. Scale bar, 50 µm. **(J)** AlphaFold3 models of NGN2-NGN2, NGN2-TCF3 and TCF3-TCF3 dimer pairs on CATATG, CAGATG and CAGGTG motifs. **(K)** Scheme describing preferential binding of several TFs to the E-box motif variants

**Figure S6.**
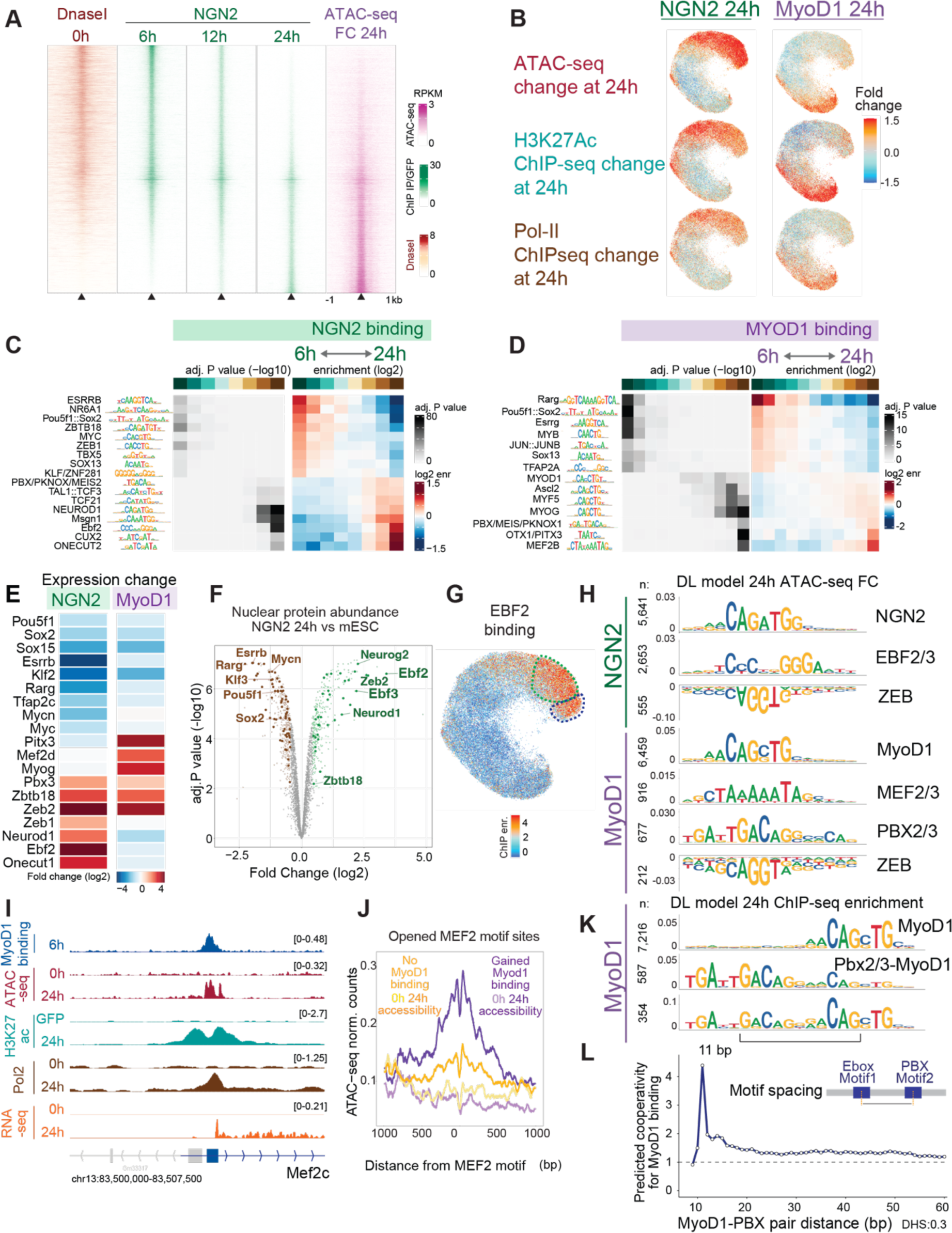
NGN2 and MyoD1 binding patterns change through differentiation as the chromatin landscape is reshaped by the downstream regulated TFs, related to Figure 6 (A) NGN2 binding illustrated as heatmap plots of NGN2 ChIP-seq enrichments at 6-, 12- and 24h NGN2 induction across combined peaks from all time-points, visualized together with initial accessibility and ATAC-seq change at 24h post induction. (B) NGN2 and MyoD1 ATAC-seq, H3K27Ac ChIP-seq and Pol2 ChIP-seq enrichment fold changes upon 24hour post induction visualized on UMAP of genomic regions. **(C** and **D)** Differentially enriched TF motifs contrasting 6h and 24h binding enrichments for NGN2 and MyoD1, together with adjusted p values and log2 enrichments identified with monaLisa motif enrichment analysis tool. **(E)** Gene expression change of TFs corresponding to motifs identified in C-D, upon 24h induction. **(F)** Volcano plot of differential nuclear protein abundances between mESCs and 24h NGN2 induction, highlighting increase in EBF2 and EBF3 abundance and decrease in pluripotency associated TFs. Proteins with -log10 adj.*P* >2 and log2 FC >0.4 are colored in transparent brown for mESC and transparent green for NGN2 (TFs indicated with solid color dots) against all detected proteins in grey. (Supplementary Table 6). **(G)** Visualizing EBF2 binding (in embryoid bodies, GSE114176^16^) on the UMAP as in Figure 6. **(H)** Top positive contributing TF motifs, identified with TF-MoDISco on CNN predictions for NGN2 and MyoD1 24h ATAC-seq fold-changes. **(I)** Genome browser tracks of MyoD1 ChIP-seq, ATAC-seq, H3K27Ac ChIP-seq, Pol2 ChIP-seq and RNA-seq at Mef2c locus as an example of a downstream activated TF expression upon MyoD1 induction. **(J)** Metaplots showing ATAC-seq signal pre and post MyoD1 induction at 24h across genomic regions with a Mef2 motif that gain accessibility, sub-grouped according to MyoD1 binding at 24h (bound purple, unbound yellow). The increase in ATAC-seq signal shows gained accessibility in regions with a Mef2 motif upon induction, and the regions with a Mef2 motif and MyoD1 motif gain MyoD1 binding and higher accessibility. **(K)** Top contributing TF motifs, identified with TF-MoDISco based on the contribution scores derived from the CNN model of MyoD1 24h binding. **(L)** Predicted cooperativity between PBX and E-box motifs for MyoD1 24h binding as a function of distance, highlighting optimal 11 bp distance and higher cooperativity up to 15bp.

**Figure S7.**
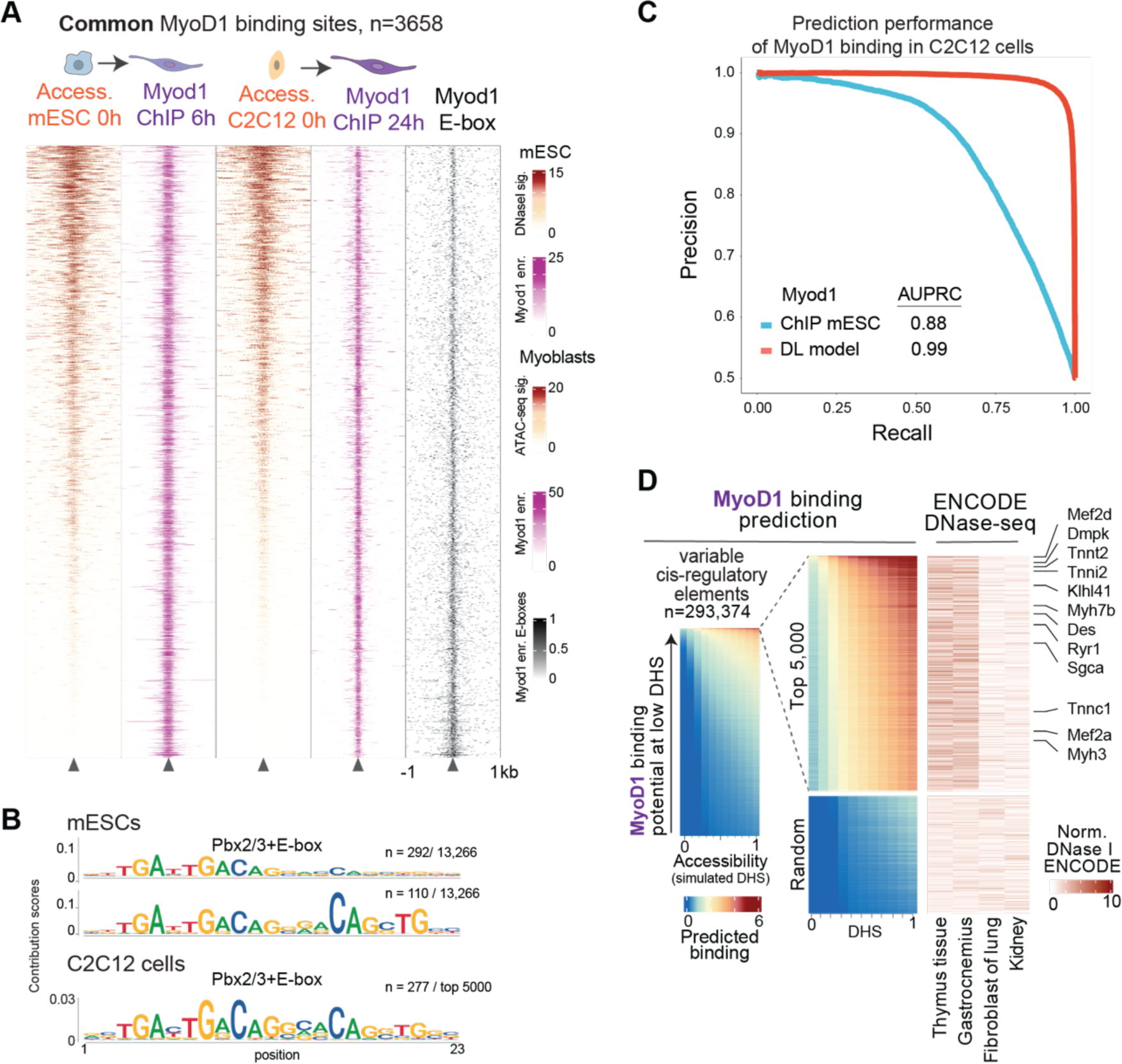
Transfer of predictive features to other cell origins, related to Figure 7 **(A)** Heatmap of overlapping MyoD1 peaks between mESCs and C2C12 cells (subset of Figure 7A) showing pre-induction DNA accessibility, MyoD1 ChIP-seq and MyoD1 E-box motifs. Overlapping regions largely exhibit common high accessibility or high motif density. **(B)** TF-MoDISco identified motifs in addition to E-box motifs alone in MyoD1 peaks across mESCs and C2C12 cells, highlighting heterotypic motif of Pbx and E-box motif pairs. **(C)** Comparing precision recall curves of CNN model, linear model with PWM motif and MyoD1 ChIP-seq measurements in mESCs for predicting MyoD1 binding enrichments in C2C12 cells **(D)** Predicted MyoD1 binding enrichment in all mouse ENCODE cCREs (rows) as a function of simulated initial DNA accessibility (columns), sorted based on MyoD1 binding potential at low DHS (left panel). Top 5000 tissue specific cCREs with highest MyoD1 binding potential at low DHS contrasted with 2000 randomly sampled tissue specific cCREs (middle panel). DNase I accessibility counts corresponding to the cCREs in the middle panel, showing top two highest and two lowest correlating tissue profiles available in ENCODE datasets (right panel). The specific DNase-seq profiles indicate that the cCREs with high MyoD1 binding potential gain activity in thymus and gastrocnemius (muscle) tissues. Example CREs proximal to the MyoD1 upregulated genes, observed in mESC to myocyte differentiation model, are labeled.

## METHODS

### Mouse embryonic stem cells (mESCs) culture maintenance

mESCs were cultured as described previously^1^. Briefly, cells were grown for several passages at 37°C 7% CO2, seeded on plates coated with 0.2% gelatin (Sigma) and maintained in serum/LIF (Dulbecco’s modified Eagle medium (DMEM, Invitrogen), supplemented with 15% fetal calf serum (Invitrogen), L-glutamine (Gibco), nonessential amino acids (Gibco), 0.001% 2-Mercaptoethanol (Sigma) and leukemia inhibitory factor (produced in-house).

### RMCE (recombinase-mediated cassette exchange) system

mESCs containing an RMCE cassette (detailed in a prior study by Lienert *et al.*^2^) were subjected to hygromycin selection (250 μg/ml, Roche) for a period of 10 days. Subsequently, four million cells were electroporated using the Amaxa Nucleofection system (Lonza) in 100-µl volumes, which included 95 µl of Nucleofector solution and 5 µl of a targeting plasmid mixture (comprising 25 μg of the L1-insert-1L plasmid and 15 µg helper construct pIC-Cre). Two days post-transfection, negative selection with 3 μM Ganciclovir (Roche) was carried out for 10 days. Individual clones were then analyzed for insertion.

### Single locus inserted inducible transcription factor (TF) expression

To achieve stable, regulated, and inducible transcription factor expression with an epitope tagging suitable for effective chromatin immunoprecipitation, we engineered mESC cells utilizing the tetON3G (Clontech 631346) and RAMBIO systems.

mESCs (129-C57Bl/6 strain) carrying a single stable RMCE acceptor cassette^3^ (L1-HygR-TK-1L) insertion and homogenous BirA-V5 expression were used as a parental cell line (HA36CB1, described in Baubec *et al.*^3^) which has shown stable and uniform expression and epitope tagging of various reporter constructs inserted by RMCE. To establish the tetON system with a well-controlled and effective expression, we first modified the cells to report on the doxycycline inducible activity. HA36CB1 cells were recombined with TRE:Luciferase (TRE3G promoter from Clontech, driving expression of Luciferase) and subjected to RMCE selection. pEF1a-Tet3G plasmid (Clontech, carrying the engineered reverse tetracycline transactivator (rtTA) with ubiquitous promoter), was randomly integrated and subjected to Puromycin selection. Single clones were then tested for Luciferase expression with and without doxycycline induction to provide the highest non-leaky inducibility. The TRE:Luciferase cassette was replaced back with L1-HygR-TK-1L through CRE-mediated recombination followed by hygromycin selection to recreate the open RMCE cassette, resulting in the HA36CB1Tet3G cells.

HA36CB1Tet3G mESCs were recombined with individual L1-TRE-biotag-TRE-T2A-GFP plasmids (where the tetracycline responsive element (TRE) promoter drives the expression of BioTag, a transcription factor, T2A in-frame cleavage peptide, and GFP). Single clones were grown for one week, split into two, and selected based on their uniform GFP expression and subsequent differentiation upon doxycycline induction in induction media (DMEM/F12/Glutamax (LifeTech, no. 31331-028), 1× B27 without vitamin A (LifeTech, no. 12587-010), 1× N2 supplement (LifeTech, no. 17502-048), 1 ug/ml doxycycline (Sigma, no. D989)).

### Neuronal and myocyte differentiation

Previously established HA36 mESCs containing random integration of the pTRE-*Ngn2* construct (as described in Kaluscha *et al.*^4^) were trypsinized and seeded on poly-D-lysine/laminin-coated plates containing induction media supplemented with 10 ng/ml human epidermal growth factor (LifeTech, no. PHG0315) and 10 ng/ml human fibroblast growth factor (LifeTech, no. CTP0261) as described in Kaluscha *et al*.^5^, which was modified from Thoma *et al.*^6^ (Figure S1A-D).

For controlled TF induced differentiation with minimal change of the culture components, HA36/birA mESCs stably expressing rtTA and single locus inserted TRE driven TFs (followed by in frame cleavage signal and GFP) were treated with mESC media (as in Casey, 2018), containing 1 μg/ml doxycycline (Figure S1F). Despite consistent expression (detected by GFP fluorescence), the protein levels of NGN2 (detected by immunofluorescence and proteomics) were low and heterogenous in serum/LIF, potentially due to instability of this protein as previously described^7^ resulting in only few differentiated cells up to 72 hours. To circumvent this, HA36 mESCs, stably expressing rtTA and single locus inserted TRE driven expression of TF constructs (tre:ngn2-t2a-gfp, tre:myod1-t2a-gfp, tre:ngn2mutant-t2a-gfp, tre:myod1mutant-t2a-gfp) were treated with induction media (as described above: DMEM/F12/Glutamax, B27 without vitamin A, N2 and doxycycline) up to three days. The parallel treatment of tre:gfp cell-line in induction media was carried out along all experiments as induction control. Cells were harvested with TrypLE at multiple time-points for downstream profiling., HA36 mESCs, stably expressing rtTA and single locus inserted TRE driven expression of TF constructs (tre:ngn2-t2a-gfp, tre:myod1-t2a-gfp, tre:ngn2mutant-t2a-gfp, tre:myod1mutant-t2a-gfp) were treated with induction media (as described above: DMEM/F12/Glutamax, B27 without vitamin A, N2 and doxycycline) up to three days. The parallel treatment of tre:gfp cell-line in induction media was carried out along all experiments as induction control. Cells were harvested with TrypLE at multiple time-points for downstream profiling.

### Light microscopy

For immunofluorescence imaging, mESCs, seeded on poly-L-lysine-coated eight-well μ-Slides (Ibidi), were fixed with 4% formaldehyde in PBS for 15 min, washed with PBS, permeabilized in PBS with 1% bovine serum albumin (BSA) and 0.1% Triton X-100 for 30 minutes, then incubated overnight with primary antibody in PBS with 1% BSA and 0.1% Tween-20. Samples were washed with PBS with 0.1% Tween-20 and incubated with secondary antibody (unless primary antibody is directly coupled with a fluorophore) in PBS with 1% BSA and 0.1% Tween-20 for 1 hour at room temperature. Nuclei were counterstained with Hoechst 33342 (1:1000 dilution, 134406, Thermo Fisher Scientific) in PBS incubated for 10 minutes and then washed with PBS. Cells were imaged with a Visitron Spinning Disk W1 microscope with a ×40 objective. Images were processed with Fiji/ImageJ^8^ and exported in RGB format. Within experiments, the imaging settings were kept the same and the visualization settings were propagated to each image.

For inspection of cellular morphology upon NGN2 induction, cells were imaged with Bio-Rad ZOE Fluorescent Cell imager utilizing cytoplasmic GFP expression of the tre:ngn2-t2a-gfp cells. Images were processed with Fiji/ImageJ^8^. Composite images of the cells together with segmented axonal projections (Fiji Tubeness Plug-in^9^) were exported.

For live-cell imaging, mESCs were seeded on poly-L-lysine-coated eight-well μ-Slides (Ibidi). mESCs were induced with doxycycline (tre:ngn2-t2a-gfp and tre:myod1-t2a-gfp cell lines in induction media for imaging the cellular differentiation and tre:gfp cell line in FCS/LIF media for imaging the mESC maintenance). Samples were imaged with a Visitron Spinning Disk W1 microscope using a 488-nm laser and 20x air objective in 12-minute intervals while being kept at 37 °C and 7% CO2 in an incubation chamber. Time-lapse images were processed with ImageJ^10^ and exported in movie format.

Primary antibodies for immunofluorescence: anti-Nanog (1:100, abcam, EPR20694), anti-BetaIII Tubulin (1:1000, Millipore, AB9354), anti-a-Actinin (1:500, Sigma, A7811), anti-V5 (1:500, Invitrogen, ma5-32053), Alexa 568 coupled Streptavidin (1:1000, Invitrogen, s11226) Secondary antibodies for immunofluorescence: Alexa Fluor 568 anti-chicken (1:500, A11041 invitrogen), Alexa Fluor 647 anti-mouse (1:500, A21235 invitrogen), Alexa Fluor 488 anti-rabbit (1:500, A11008 invitrogen)

### Chromatin Immunoprecipitation Sequencing (ChIP-seq)

ChIP was carried out as previously described^11^ with the following modifications: (1) 10 million cells were fixed in suspension, (2) chromatin was sonicated using a Diagenode Bioruptor Pico for 22 cycles of 30s on, 30s off, (3) protein A magnetic Dynabeads (Thermo Fisher, 10008D) were used.

BioTag ChIP was carried out as previously described in Baubec *et al.*^3^ with the following modifications: (1) 10 million cells cells were fixed in suspension for 8 minutes (2) nuclei were lysed in 50 mM HEPES, 1 mM EDTA, 1% Triton X-100, 0.1% deoxycholate, 0.1% SDS, and 500 mM NaCl for 1 hr on ice, (3) crosslinked chromatin was sonicated using a Diagenode Bioruptor Pico for 22 cycles of 30s on, 30s off.

Antibodies for pull-down of the target proteins for ChIP-seq were anti-Pol2 antibody (Santa-Cruz, sc899x), anti-H3K27ac antibody (Active Motif, 39133), anti-V5 antibody (Invitrogen, MA5-32053) and in case of biotin as a bait Streptavidin (Thermo Fisher, 11205D).

Immunoprecipitated DNA was subjected to library preparation using NEBNext Ultra II DNA Library Prep Kit for Illumina and amplified with 12 cycles. Libraries were sequenced on an Illumina HiSeq (50 cycles) or NovaSeq (paired-end, 100 cycles).

ChIP–seq datasets were aligned to mouse genome mm10 assembly using Bowtie2 aligner^12^ accessible from the Bioconductor package, QuasR^13^ with default settings.

Peak calling on TF ChIP-seq samples was performed with MACS2^14^ (v.2.1.3.3) using the callpeak argument with default settings using the *P* value threshold of 1e-3. Genomic peaks with low mappability^15^ (less than 80%) or that overlapped with blacklisted regions^16^ were removed from further analysis.

To define consensus peak sets for NGN2 and MyoD1 binding at 6 hours, MACS2 peaks across three replicates were combined using R package DiffBind^17,18^ (v.3.14.0). The DiffBind peak summits were further processed in R^19^. TF ChIP enrichments at 500 bp (average ChIP-seq peak length) genomic regions centered at the peak summits were quantified with QuasR^13^ package (v1.44.0) with default parameters and shifting the reads by half the estimated size of average ChIP-seq library fragments. Counts were log2 transformed with a pseudo-count of 8, to reduce noise at low read counts. Normalization was performed according to library size for each sample (multiplying counts by a scaling factor making the total counts equal). Enrichments of log2 ChIP–seq read counts were calculated by subtracting the experiment matched log2 counts of the GFP pull-down. To define a reliable set of consensus peaks, irreproducible discovery rate^20^ (IDR) analysis (as described in the Encode project, v1.3) was used, with a threshold of IDR < 0.01, log2 normalized ChIP-seq counts >6 in at least two replicates, average ChIP-seq enrichment relative to corresponding DNA binding mutant >1, average ChIP-seq enrichment relative to GFP >1.5 and a minimum ChIP-seq enrichment of 0.5 in each replicate, following an approach similar to Isbel *et al.*^21^.

To generate unified regions of interest for various analysis, peaks from different samples and treatments were pooled. Peaks with more than 70% overlap were merged and resized from the center using GenomicRanges^22^ (v.1.56.1). This allows to have a unified peak set without losing relevant resolution for potential differential signal among TFs or other chromatin features. Read counts were generated over the peak regions using qCount function of the QuasR ^13^ package with default parameters and shifting the reads by half the size of ChIP-seq library fragments. Counts were log2 transformed and nEnrichments of log2 ChIP-seq read counts were calculated by subtracting the matched log2 counts from the corresponding control datasets (matched input sequencing for public data, IgG ChIP-seq for antibody pull-down and GFP ChIP-seq for BioTag pull-down).

### RNA sequencing (RNA-seq)

For each sample, RNA from 50,000 cells was isolated using Norgen Single Cell RNA Purification Kit following the manufacturer’s instructions. Sequencing libraries were prepared using Illumina TruSeq Stranded Total RNA Library Preparation Kit except for Figure S2B-C which was prepared using the mRNA Library Preparation. Libraries were single-end sequenced on a HiSeq 2500 platform with 50 cycles. lllumina RTA 1.18.64 (HiSeq 2500) and bcl2fastq2 v.2.17 was used for base calling and demultiplexing.

Using STAR aligner ^23^ (v2.5.2b), RNA-seq reads were aligned to the mm10 reference genome with default parameters. The aligned files were subsequently sorted and indexed with SAMtools ^24^ (v1.9). Protein-coding genes were quantified employing the Rsubread ^25,26^ (v2.18.0) tool from the Bioconductor ^27^ suite. Specifically, the featureCounts function in Rsubread was executed using standard settings along with gene annotations from the M25 GENCODE ^28^ release, setting GTF.attrType to ‘gene_name’ for grouping gene elements, such as exons. In each pairwise comparison, the count matrix was subsetted for the samples under comparison and the scaling factors were calculated using edgeR’s ^29^ calcNormFactors function with the default parameters. Differentially expressed genes were identified using the edgeR^29^ package with the glmQLFit and glmQLFTest functions. *P* values were corrected for multiple testing using the Benjamini–Hochberg^30^ method. Log2 fold change estimates were derived using the predFC function in edgeR ^29^ (v4.2.0). For each differential analysis comparison, genes were considered to be differentially expressed with a false discovery rate (FDR) < 1e-3, an absolute log2 fold change of at least 1 and average logCPM of > -2. Gene expression profiles were characterized by the gene set enrichment analysis^31^ (GSEA) using the mouse MSigDB^32^ collection as the reference gene set database (excluding M2 positional gene sets and gene sets names containing MIR:MIR Legacy or MIR:MIRDB). The pre-ranked approach (log2 fold change) was employed from R fgsea^33^ package (v1.30.0) to perform GSEA with the default parameters. Libraries were normalized using the cpm function in edgeR package following the filtering step for low expression (filterByExpr). Principal component analysis (PCA, prcomp from R) of the gene expression profiles was performed using the top 5000 variable genes across samples derived from distinct in vitro differentiation trajectories.

To uncover enriched motif variants associated with genes that are upregulated along the NGN2-induced gene expression trajectory, we conducted a comprehensive analysis over multiple time points (6, 12, and 24 hours). We identified genes exhibiting progressively increasing or decreasing expression by comparing their log2 fold changes to those of mESCs at 0 hours, ensuring each sequential time point demonstrated an equal or higher expression level than its predecessor. We utilized the clusterProfiler^34^ package (v4.12.0) to perform gene annotation enrichment analysis on 1,687 upregulated and 1,197 downregulated genes identified for NGN2-driven neurogenesis, employing GO:BP^35^ gene sets as the gene annotation resource (Figure S4J). Additionally, NGN2 binding sites, gathered from the same 6, 12 and 24-hour post-induction time points, were compiled and merged using the reduce function from GenomicRanges, with a criterion of ChIP IP/GFP enrichment greater than 1 at any of the time points. These sites, located within 25kb of the gene TSS, were mapped to the progressively-regulated genes using the annotatePeak function from the ChIPseeker^36^ package (v1.40.0) based on the gene annotation from TxDb.Mmusculus.UCSC.mm10.knownGene (v3.10.0). We divided the mapped TF binding sites into three bins—corresponding to upregulated genes, downregulated genes, and others. An enrichment analysis of 10-mer E-box motifs (NNCANNTGNN) was subsequently performed for each bin using the monaLisa^37^ Bioconductor package (v1.10.0), with “otherBins” as the background and a minimum score of 10 to ensure a perfect match for 10-mer motifs. Motifs showing an enrichment greater than 0.5 and an adjusted *P* value below 0.01 were highlighted in Figure S4J.

The differential gene expression analysis for NGN2 (6h, 12h, 24h) and MyoD1 (6h, 24h) is available in Supplementary Table 7.

### ATAC–seq

ATAC–seq was performed as previously described^38^. Briefly, 50,000 cells were resuspended in lysis buffer to extract nuclei. Nuclear pellets were incubated with transposition reaction buffer for 30 min at 37 °C. DNA was purified using the PCR Purification Kit (Qiagen). Eluted transposed DNA was amplified with 7 cycles of PCR using Q5 High-Fidelity Polymerase (NEB). Libraries were sequenced paired-end with 75 cycles on Illumina NextSeq platform.

Illumina RTA 2.4.1 (NextSeq 500) and bcl2fastq2 v.2.17 were used for base calling and demultiplexing.

ATAC–seq reads were trimmed using cutadapt^39^ (v.1.18) with parameters -a CTGTCTCTTATACACA -A CTGTCTCTTATACACA -m 10–overlap = 1 and then mapped to mm10 reference genome using Bowtie 2 aligner^12^ via QuasR with default settings. ATAC-seq peaks were called with MACS2^14^ for each sample with the paired-end analysis mode (BAMPE) and the *P* value threshold of 1e-3. Peaks overlapping mm10 blacklisted^16^ regions or having low mappability^15^ (<80%) were removed. Peaks across replicates were merged if the resized peaks (±250 bp) were overlapping more than 70% from the peak center.

Read counts at all ATAC-seq peaks were quantified using qCount function from QuasR package with default parameters. To ensure comparability across samples, we normalized these counts using the cyclicLoess method^40,41^ available from the limma^42^ package (v3.60.3). This normalization step is crucial as it adjusts for variations in library size and TN5 reaction efficiency, with the assumption that the majority of data points, consisting predominantly of ubiquitously active regulatory and structural elements, do not change. Normalized signal was further log2 transformed with a pseudo count of 8, to account for noise at low read counts and the resulting normalized matrix was used to calculate log2 fold changes between samples. Subsequently, for each pairwise sample comparison, we identified differentially accessible regions using the edgeR^29^ package (v4.2.0) by the glmQLFit and glmQLFTest functions.

### Differential Affinity Purification Sequencing (DAP-seq)

DAP-seq protocol was adapted from Bartlett *et al.*^43^. To prepare a stock of fragmented genomic DNA, 500 million mESCs devoid of DNA methylation (i.e. DNMT TKO cells^44^) were harvested and lysed in Lysis buffer (20 mM Tris pH 8, 4 mM EDTA, 20 mM NaCl, 1% SDS) with 1 ml ProtK (10 mg/ml) at 55°C overnight. Genomic DNA was isolated with phenol/chloroform extraction. DNA pellet was resuspended in 4 ml elution buffer (10 mM Tris-Cl, pH 8) with 50ug/ml RNAseA and incubated at 37°C overnight. DNA was fragmented using sonication with Covaris S220 to reach 150-250bp length. Fragmented DNA was ethanol precipitated and resuspended in elution buffer (10ug/ul).

To produce transcription factor proteins, Pgex6.1 plasmid constructs encoding GST tagged Ngn2, Myod1 and bHLH domains of Tcf3e12, Tcf3e47, Tcf4 and Tcf12 were introduced to Rosetta (DE3) competent cells and single clones were grown in LB with Chloramphenicol (0.25 µl/ml) and Carbenicillin (0.5 µl/ml). 4 ml of the liquid culture were transferred to MgSO4 containing 100 ml LB media with antibiotics (Carbenicillin and Chloramphenicol) and grown until OD 0.5 was reached. Protein production was induced with IPTG (40 µl in 100 ml) and the culture was grown at 37°C 3 hours for NGN2 and MyoD1, and at 16°C 3 hours for TCF bHLHs (since these conditions gave the highest soluble concentrations with limited degradation products). Cultures were spun at 6000 rpm for 10 minutes at 4°C and the pellets were frozen at -80°C overnight. Pellets were resuspended in cold lysis buffer 16 ml lysis buffer (50mM Tris pH8, 0.2% Tween20, 300 mM NaCl, 3 mM MgCl2, 1 mM TCEP) with lysozyme and incubated at 4°C for 1 hour. Lysates were sonicated in 15ml falcons using Bioruptor (15 s ON 15 s OFF for 1min (total 2 minutes) at 50% duty cycle) in ice water, preventing heating, and centrifuged at 13000 g for 45 minutes at 4°C.

To pair and isolate the proteins, supernatants were mixed according to desired dimer pairs and incubated rotating for 10 minutes at room temperature. Five dilutions of the lysate mix were generated by adding lysis buffer (1x, 0.5x,0.25x, 0.13x, 0.06x). To bind the transcription factors on beads, 10 µl of MagneGST beads (Promega) per sample were washed with lysis buffer, added to the lysate mix dilutions and incubated rotating for 1 hour at room temperature. Supernatants were removed on the magnetic racks and the beads were washed with lysis buffer. Beads bound by TFs were resuspended in 100 µl lysis buffer and transferred to 96 well plates. Lysis buffer is replaced on the magnetic rack and the beads were resuspended in 80 µl PBS.

To probe transcripiton factors for preferentially bound DNA fragments, PBS was replaced with 10ug fragmented genomic DNA diluted in 80 µl PBS. Samples were incubated rotating horizontally at room temperature for 1 hour. During the following steps beads were handled with minimum physical disruption (i.e. letting the beads fall naturally within the buffer during washes without pipetting directly or shaking and not letting the beads dry). Supernatant was removed on the magnetic rack and the beads were washed five times with 200 µl of PBS with 0.005% NP40 (each wash for 2 minutes and on the magnet for 1 minute). For the final wash, the samples were transferred to new tubes and washed with PBS twice. To recover bound DNA, beads were resuspended in 30 µl of elution buffer with 2 µl 10 mg/ml proteinase K and incubated at 50°C for 3 hours. Beads were removed and DNA was purified with 2X AmpureXP beads, eluted in 30 µl elution buffer. To determine the protein dilution range that gives the highest specific DNA enrichment, samples were tested with qPCR using primers corresponding to a region in the Dll1 locus that contains multiple E-box motifs and a control region devoid of E-box motifs. Sample replicates were subjected to library preparation with 12 cycles of amplification (NEBNext Ultra II DNA Library Prep Kit, Illumina). Libraries were sequenced on Illumina HiSeq (50 cycles) or NovaSeq (paired-end, 100 cycles).

Pull-down of fragmented genomic DNA with magneGST beads alone (without TFs) were used as a background enrichment control. Reads were mapped to mouse mm10 assembly with QuasR^13^ package. Peaks were called with MACS2 using default parameters. Peaks across samples were pooled and peaks overlapping more than 70% were merged. DAP-seq enrichments at 500 bp (average ChIP-seq peak length) genomic regions centered at the peak summits were quantified with the qCount function in QuasR package with default parameters shifting the reads by half of the fragment size. Counts were log2 transformed with a pseudo-count of 8, controlling for noise at low read counts. Normalization was performed according to the library size of each sample (multiplying counts by a scaling factor making the total counts equal). Enrichments of log2 ChIP–seq read counts were calculated by averaging the replicates and subtracting the log2 counts of GST alone pull-down. To perform differential E-box motif enrichment analysis between TF dimer pairs, top 15000 enriched peaks of the unique dimer pairs were analyzed using monaLisa^37^ Bioconductor package (v1.10.0). Library of 8-mer E-box motifs (NCANNTGN) with parameters set to “otherBins” for comparison and min.score=8 for ensuring perfect match to the 8-mers were used. E-box motifs, with enrichment > 0.2, -log10(adjusted p value) > 5, were visualized in Figure 5C as dotplots where the size of the dots represents the *P* values, and the color represents log2 enrichments. For motif enrichment against genomic background, the parameter “genome” was used as illustrated in Figure S5E.

### qPCR primers

Positive control forward: AAGCTTTTAGTGCTCCAGGC Positive control reverse: AAGCCTCTCATTGTGCCACG Negative control forward: TCACACTTGTCTTCTGGTCTCAC Negative control reverse: CAAGGTGACCTAGAAACTTGC

### Stably inserted transcription reporters

Plasmid constructs that contain LoxP flanking inserts composed of E-box motifs embedded in an in-vitro-derived background sequence (background sequence is as described in Grand *et al.*^15^ for the transcription reporter of BANP motifs) followed by TATA box, Nano-Luciferase open reading frame, construct specific barcodes and polyA were synthesized by TWIST bioscience. Mixed plasmid library was amplified by cloning equally mixed plasmid constructs in E.Coli.

To generate mESCs with NGN2 induction system and an open RMCE site characterized for providing neutral background, Piggybac plasmid with TetON system carrying doxycycline inducible NGN2-T2A-GFP expression (CAGS:rtTA/TRE:NGN2-T2A-GFP/NeoR) was randomly integrated to TC-1 mESCs (background 129S6/SvEvTac, carrying an RMCE selection cassette^2^) with dual helper plasmid as described previously^4^. Briefly, 4 million cells were electroporated with 4.5 µg plasmid mix (3.8 µg Piggybac plasmid and 0.7 µg Dual helper construct). Starting 2 days after, cells were subjected to G418 (Neomycin) selection for 2 weeks. Single clones were then tested for doxycycline inducible NGN2-T2A-GFP expression and neuronal differentiation.

The expression reporter constructs were stably integrated to the mESCs via RMCE. After 10 days of Ganciclovir negative selection, cells were expanded for 2 days and induced with induction media to express NGN2. Genomic DNA was extracted via cell lysis followed by AMPURE XP beads with isopropanol. RNA was extracted using Norgen Single Cell RNA purification kit. Isolated RNA was reverse transcribed using Superscript IV reverse transcriptase with a specific primer targeting the downstream sequence of the barcodes. cDNA and genomic DNA were amplified with same primer pair for 8 cycles. The resulting DNA was subjected to Nextera XT DNA Library Preparation and sequenced with MiSeq Micro v2.

Reads were mapped to the construct library using QuasR, where each construct has a specific barcode sequence. RNA and genomic DNA samples were scaled to normalize to the same total number of counts. Enrichments of barcodes in the RNA samples were calculated by dividing the RNA counts with the genomic DNA counts, representing the proportion of RNA counts per cell (as described in Hartl *et al.*^45^). Enrichments of each construct relative to the control construct (with scrambled motifs) were calculated and averaged across two replicates and visualized as heatmaps (Figure 4F, Figure S4H, Supplementary Table 5).

Reverse transcription primer: GTCTCGTGGGCTCGGAGATGT Amplification forward primer: GCACCCTGTGGAACGGCAAC Amplification reverse primer: GGGCTCGGAGATGTCCTAGG E-box motifs used: GACAGATGGT (referred as GA or GAg), GACAGATGTT (GAt), GACATATGGT (TA), GACAGCTGGT (GC), GACAGGTGGT (GG), GACAGATGCT (GAc), GCCAGATGGT (cGA)

### Generation of Tcf3-V5

TC-1 mESCs were modified with CRISPR to insert coding sequence of V5 tag to the C terminus of endogenous TCF3 gene just before the stop codon using IDT Alt-R CRISPR-Cas9 system. Custom guide RNA (cRNA and tracrRNA) and single stranded DNA oligo (ssODNs) for homologous recombination were synthesized by IDT. cRNA (Alt-R® CRISPR-Cas9 crRNA) and tracRNA (Alt-R® CRISPR-Cas9 tracrRNA, 1072533) were annealed to obtain 100 µM gRNA, by mixing one to one, heating 95°C for 5 minutes and cooling to room temperature. To assemble the active Cas9 complex, 1 µl of gRNA (100pmol) and 0.8 µl Cas9 protein (50 pmol, Alt-R S.p. Cas9 Nuclease V3, 1081058) were mixed and incubated for 15 minutes at room temperature. 2 nmol electroporation enhancer (Alt-R Cas9 Electroporation Enhancer) and 2nmol ssODN (Alt-R HDR Donor Oligo) and 17 µl Lonza P3 primary cell nucleofection buffer were added to the mix. 60,000 mESCs were resuspended in this 20 µl of the mix and nucleofected using Amaxa 4D nucleofection system with CA-120 program (as described in Dewari *et al.*^46^). Cells were transferred to 24-well culture plates containing 500 µl Serum/LIF media supplemented with homologous recombination enhancer (Alt-R HDR Enhancer) and grown for 12 hours. Media was replaced with Serum/LIF and cells were subjected to clonal selection. Colonies originating from single cells were screened by Sanger sequencing of the tagged locus and anti-V5 immuno-fluorescence staining. Cells containing V5 tagged Tcf3 allele were expanded. Cells were further verified with V5 staining upon 24-hour Tcf3 siRNA treatment.

Guide RNA seed: GTCCAACGAAGAAGCTGTGA ssODN: CCGGGCCTGGGTGAGGCCCACAACCCAGCCGGGCACCTGGGTAAGCCTATCCCTAAC CCTCTCCTCGGTCTCGATTCTACGTGAG<u>G</u>CG<u>TCACAGCTTCTTCGTTGGAC</u>CAGGGAC CACCATA

*to prevent cutting after insertion, underlined single base was modified from CC to GC to mutate the PAM site for the corresponding guide RNA

Genotyping primers:

Forward: AGGAGGAGAAGGTGTCTGGC Reverse: AGCTGTTCAGTGCCACCCTC

### Protein affinity-purification mass spectrometry (IP-MS)

Affinity purification for LC–MS/MS was performed based on Ostapcuk *et al.*^47^ with modifications. mESCs expressing NGN2, MyoD1 or GFP (6h and 24h) in triplicates of 10 million cells per sample were harvested. Cells were resuspended in 1ml B150 (10 mM Tris-HCl pH7.5, 2 mM MgCl2, 150 mM NaCl, 0.5% Triton X-100) with AG (50 mM L-Arg (Sigma A5006), 50 mM L-Glu (Sigma G1251)) and PIC (1xPIC Protease Inhibitor Cocktail). Cells were lysed by vortexing, spun for 5 minutes at 4°C 13000g and the supernatant was transferred to a new tube. 10 µl M280 Streptavidin Dynabeads per sample were washed with B150+AG+PIC and added to the lysate to bind the target proteins to the beads. Samples were rotated for 4 hours at 4°C and washed three times with 1ml B150. To remove the detergent, beads were resuspended in 250µl B150ND (10mM Tris-HCl pH7.5, 2mM MgCl2, 150mM NaCl) and transferred to new tube. Samples were washed with 1ml B150ND and supernatant was removed. Beads were resuspended in 5 µl digestion buffer (3 M guanidine-HCl, 20 mM EPPS pH 8.5, 10 mM CAA, 5 mM TCEP) and 1 µl Lys-C and incubated at room temperature for 4 hours. 17 µl 50 mM HEPES pH 8.5 and 1 µl 0.2 µg/µl trypsin was added to the samples, incubating overnight at 37 °C. For complete digestion, additional 1 µl of 0.2 µg/µl trypsin was added, incubating for 5 hours at 37 °C. Samples were acidified by adding 1 µl of 20% trifluoroacetic acid (TFA) and sonicated in an ultrasound bath.

Peptides were analysed by capillary liquid chromatography tandem mass spectrometry with an EASY-nLC 1000 using the two column set-up (Thermo Fisher Scientific). The peptides were loaded with 0.1% formic acid, 2% acetonitrile in H2O onto a peptide trap (Acclaim PepMap 100, 75um x 2cm, C18, 3um, 100Å) at a constant pressure of 800 bar. Peptides were separated, at a flow rate of 150 nl/min with a linear gradient of 2–6% buffer B in buffer A in 3 minutes followed by an linear increase from 6 to 22% in 40 minutes, 22-28% in 9 min, 28-36% in 8min, 36-80% in 1 min and the column was finally washed for 14 min at 80% B (Buffer A: 0.1% formic acid in water, buffer B: 0.1% formic acid in acetonitrile) on a 50um x 15cm ES801 C18, 2um, 100Å column mounted on an EASY-Spray™ source connected to a Orbitrap Fusion (Thermo Fisher Scientific). The data were acquired using 120000 resolution for the peptide measurements in the Orbitrap and a top T (3s) method with HCD fragmentation for each precursor and fragment measurement in the ion trap according the recommendation of the manufacturer (Thermo Fisher Scientific).

Thermo RAW files were processed using MaxQuant 1.5.3.8 software (Max Planck Institute of Biochemistry) with Andromeda search engine with label-free quantification (protein and peptide FDR values set to 1% and 0.1% respectively) against the Mus musculus UniProt database v.2017_04 supplemented with common contaminating proteins. The combined intensities of peptides of proteins were imported into R and normalized between samples to the same total count of proteins. Protein counts were log2 transformed and TF pull-down samples were compared with GFP pull-downs. The significance estimates were calculated using the eBayes function in the limma R package. Adjusted *P* values and log2 fold enrichments of proteins identified in TF pull-downs were visualized in Figure 5A with names of the chromatin associated proteins (GO term(s) from Biomart-BioConductor package^48,49^) showing -log10 (adjusted *P* value)>5 and log2 enrichment>5. Enrichment of proteins detected in NGN2 6h, NGN2mut 6h, NGN2 24h and NGN2mut 24h relative to GFP IPs were normalized to bait protein and listed in Supplementary Table 8. Highly abundant proteins detected across many samples (e.g. ribosomal proteins, mitochondrial proteins) in IPs are labelled as IP gray list. Enrichment of proteins detected in NGN2 24h and MyoD1 24h IP-MS were normalized to the bait protein (i.e. NGN2 and MyoD1 respectively) and visualized together in Figure S5J and listed in Supplementary Table 9.

### Nuclear protein mass spectrometry

Proteome quantification was done as described previously in Grand, 2021^15^, with the following modifications: i) isolated nuclei were used as a starting material rather than whole cells to enable detection of transcription factors which are generally less abundant proteins in the cell, i) nine channels from a TMTpro16 kit were used. Briefly, mESCs, NGN2 24h induced cells and MyoD1 24h induced cells were harvested in triplicates of 10 million per sample. Nuclei were isolated by suspending in 1.25 ml of nuclear extraction buffer (10 mM HEPES pH 7.9, 10 mM KCl, 10 mM EDTA, 0.5% NP-40, 1 mM DTT and complete protease inhibitor (Roche)) and incubating on ice for 10 min with vortexing every 2 minutes. Nuclei were collected by centrifugation (800*g*, 10 min, 4 °C), washed with ice cold PBS and resuspended in 250 µl of nuclear disruption buffer (20 mM HEPES pH 7.9, 400 mM NaCl, 1 mM EDTA, 1% glycerol, 1 mM DTT, complete protease inhibitor (Roche) and incubated for 2 hours at 4 °C rotating, with vortexing every 30 minutes. Samples were centrifuged at 3000RPM for 10 minutes at 4 °C and sodium deoxycholate (final concentration to 1%) was added to the supernatants. The lysates were further disrupted with sonication. Proteins were reduced, alkylated and LysC/Trypsin digested.

Peptides (50µg) from each biological replicate were labelled using TMT 16plex isobaric labelling kit (Thermo Fisher Scientific). High-pH reversed-phase chromatography was employed using an Agilent 1100 HPLC system with an autosampler, a YMC Triart C18 0.5 x 250 mm column, and a fraction collector. The peptides were separated using the binary buffer system: high-pH Buffer A (20 mM ammonium formate in water, pH 10) and high-pH Buffer B (20 mM ammonium formate pH 10 in 90% acetonitrile) at a flow rate of 12 μL/minute. Samples were acidified with TFA. Peptides, reconstituted in 2% acetonitrile, 0.1% formic acid, were on-line separated on a C18 50 cm μPAC column using a EASYnLC-1000 system (Thermo) mounted on a Digital PicoView nanospray source (New Objective), connected to an Orbitrap Fusion Lumos mass spectrometer (Thermo). Peptide elution was achieved using a linear gradient of increasing concentration of acetonitrile in 0.1% formic acid at a flow rate of 500 nl/min. For MS data acquisition in a data-dependent mode, ions for survey MS1 scans were recorded in profile mode in the Orbitrap detector at a resolution of 120,000 in the range of 400-1400 m/z. MS2 HCD spectra were recorded in the in centroid mode, starting at 100 m/z at 50,000 resolution.

MS2 fragment ion spectra were searched in Proteome Discoverer^50^ v2.5 (Thermo) with the Sequest ^51^ HT search engine against the Mouse Uniprot protein database and the MaxQuant ^52^ contaminant database. The Proteome Discoverer proteins table was further analyzed using einprot ^53^ R package (v0.9.5). Contaminants were removed, master proteins were log2 transformed. Samples were median normalized and imputed using MinProb method.

Samples were compared using limma ^42, 54^ Bioconductor package. The differential analysis for mESCs versus NGN2 24h and MyoD1 24h is listed in Supplementary Table 6.

### Oligonucleotide affinity-purification mass spectrometry

Protein detection from nuclear extracts using DNA double stranded DNA oligomers (Oligonucleotide affinity purification) was performed as described previously in Grand *et al.*^55^ (which was adapted from Makowski *et al.*^55^). Briefly, 20 bp ssDNA oligomers of ACTGTAC-CANNTGn-TCAGTA with E-box motifs replacing Ns: GAg, GAt, GCg, GGg were synthesized by Microsynth. Oligos were annealed in a thermocycler and diluted in DNA binding buffer (DBB: 1 M NaCl, 10mM Tris (pH 8.0), 1mM EDTA, 0.05% NP-40) to a 4 μM concentration in 200-μl per replicate. Oligos were bound to 5 μl bead slurry (Dynabeads, Thermo Fisher Scientific, 11205D) per sample and washed with DBB. 150 million mESCs, expressing NGN2 or MyoD1 (24h induction), were harvested with TrypLE, resuspended in the induction media and pelleted (600*g*, 5 min). Nuclei were extracted by suspending in Buffer A (10 mM HEPES pH 7.9, 10 mM KCl, 10 mM EDTA, 0.5% NP-40, 1 mM DTT and complete protease inhibitor (Roche)) and disrupted in Buffer B (20 mM HEPES pH 7.9, 400 mM NaCl, 1 mM EDTA, 1% glycerol, 1 mM DTT, complete protease inhibitor (Roche)). For each sample, 80 μg nuclear extract (Qubit Q33211) in triplicates was diluted in 200 μl final protein binding buffer (150 mM NaCl, 50 mM Tris, pH 8, 0.25% NP-40, 1 mM Tris(2-carboxyethyl)phosphine (TCEP) and complete protease inhibitor (Roche)) and added to the oligo-bound magnetic beads for 2 hours at 4 °C. Samples were washed with washing buffer (150 mM NaCl, 100 mM ammonium bicarbonate). The proteins were digested with LysC (Wako Chemicals)/Trypsin (Promega) and acidified with TFA.

Peptides were analyzed by capillary liquid chromatography tandem mass spectrometry with an EASY-nLC 1000 using the two-column set-up (Thermo Fisher Scientific). The peptides were loaded with 0.1% formic acid, 2% acetonitrile in H2O onto a peptide trap (Acclaim PepMap 100, 75um x 2cm, C18, 3um, 100Å). Peptides were separated, at a flow rate of 500 nl/min with a linear gradient of 3–6% buffer B in buffer A in 4 minutes followed by a linear increase from 6 to 22% in 55 minutes, 22-40% in 4 min, 40-80% in 1min and the column was finally washed for 10 min at 80% B (Buffer A: 0.1% formic acid in water, buffer B: 0.1% formic acid in acetonitrile) on a C18 50 cm μPAC column (Thermo Fisher Scientific) mounted on a Digital PicoView nanospray source (New Objective), connected to an Orbitrap Fusion Lumos mass spectrometer (Thermo Fisher Scientific). The data were acquired using 120000 resolution for the peptide measurements in the Orbitrap and a top T (3s) method with HCD fragmentation for each precursor and fragment measurement in the ion trap according to the recommendation of the manufacturer (Thermo Fisher Scientific).

Thermo RAW files were processed using MaxQuant software^56^ (v1.5.3.8, Max Planck Institute of Biochemistry) with Andromeda search engine with label-free quantification (peptide FDR values set to 0.1%) against the Mus musculus UniProt database v.2017_04 supplemented with common contaminating proteins. Results were filtered for reverse hits and contaminants. Peptide counts were imported to R, log2 transformed and subtracted from the corresponding control sample (scrambled E-box motifs) and summarized to unique protein groups by the highest enrichment. Proteins, detected in NGN2 24h nuclear extracts, corresponding to a mouse transcription factor^57,58^ with known E-box binding activity were visualized in a heatmap (Figure 5SA). Correlations (R cor function, pearson) between samples were plotted as a heatmap (Figure S5B). Enriched proteins in either NGN2 or MyoD1 induction are listed in Supplementary Table 10. Sequence specific DNA binding proteins are labeled as TF list.

### siRNA treatments

siRNAs targeting Tcf3 (Entrez Gene 21423), Tcf4 (Entrez Gene 21413), Tcf12 (Entrez Gene 21406), as well as non-targeting control pool were purchased from Horizon Discovery in ON-TARGETplus siRNA SMARTpool format.

For testing TCF3-V5 cell line with immunofluorescence, 100,000 TCF3-V5 mESCs were seeded on Ibidi 8 well slide in FCS/LIF media containing Tcf3 siRNA or control siRNA together with RNAimax reagents (300 µl media, 34 µl Opti-MEM, 1 µl Lipofectamine and 1 µl of 4µM siRNA stock) following the manufacturer’s guidelines. 24 hours later, cells were subjected to immunofluorescence staining.

For testing the effect of decrease on TCF proteins during neuronal differentiation, 100,000 tre:ngn2-t2a-gfp mESCs were seeded on 24 well plates in 300µl FCS/LIF media containing same amount of siRNA as described above for Tcf3, Tcf4, Tcf12, mix of Tcf3/4/12 and non-targeting siRNA. 24 hours later, cells were induced in induction media, containing the same amount of siRNA. 48 hours later, GFP expressing cells were imaged.

For testing the impact of decreasing TCF protein levels on initial NGN2 engagement upon its expression in mESCs, 4 million tre:ngn2-t2a-gfp and the control tre:ngn2mutant-t2a-gfp mESCs were seeded on 10 cm plates in 11 ml FCS/LIF media with siRNAs (40µl of 4µM siRNA, 40 µl Lipofectamine, 1360 µl Opti-MEM) for Tcf3, Tcf4, Tcf12 and non-targeting siRNA control. After 24 hours, the media was replaced with induction media. 6 hours after induction, NGN2 ChIP-seq was performed. NGN2 ChIP enrichments were quantified in NGN2 6h consensus peaks. Log2 fold changes of siRNA treated samples over non targeting siRNA control was calculated and the peaks were divided in 3 bins based on their change in NGN2 enrichment. Differential motif enrichment analysis was performed using monaLisa^37^ Bioconductor package (v1.10.0). Library of 8-mer E-box motifs (NCANNTGN) with parameters set to “otherBins” for comparison and min.score=8 for ensuring perfect match to the 8-mers were used. E-box motifs, with enrichment>0.5, -log10vadjusted P value>5, were visualized (see Figure 5C) as dotplots where the size of the dots represents the p values, and the color represents enrichments.

### Downstream Data Analysis

#### Motifs for TF binding

HOMER known-motif analysis using JASPAR motif database^59,60^ and HOMER^61^ de novo motif analysis was performed for the top 500 most enriched NGN2 or MyoD1 consensus peaks that resulted in PWM of motifs containing an E-box sequence where the top enriched known motif (PWM) was found in 95% of these regions. The top PWM motifs identified by Homer de novo motif analysis (200bp around peak center)^15^ were visualized as sequence logo^62^ in bits in Figure 1A. Since PWMs do not contain dependency information between bases, to identify potential cognate motifs for NGN2 and MyoD1, we performed K-mer motif enrichment analysis (as frequently preferred for short sequence motifs) in all NGN2 or MyoD1 consensus peaks using monaLisa^37^ package (v1.10.0), with 10-mer NNCANNTGNN motif library, GC matched mouse genome background and min.score=10 to ensure a perfect match for the motif sequence. 222 out of 2080 E-box motifs (NNCANNTGNN) were found to be overrepresented for NGN2 by the criteria of adjusted *P* value < 0.01 and log2 enrichment >0.5, and similarly revealed 185 overrepresented E-box motif variants for MyoD1 (Figure S2E-F). According to these criteria, 91% of the consensus peaks had at least one enriched E-box motif (referred as cognate motif sequences) both for NGN2 and MyoD1 while 99% of the peaks had at least one match to any E-box motif.

To define all the potential binding sites for NGN2 or MyoD1, the positions of all the cognate motifs for the corresponding TF (exact match to one of the cognate motif 10-mers, described above) in Mus Musculus mm10 genome assembly were identified. TF ChIP-seq enrichments and DNAse-seq signal (DHS) were counted at these cognate motif sites (±250bp from the center of the motif). Motifs that are represented more than once due to closer distance to each other (less than 250 bp to another cognate motif) were resolved by keeping the motif that covers all other nearby motifs within this 500 bp region. This set of genomic regions are referred as cognate motif regions as potential binding sites for NGN2 or MyoD1 in the mouse genome (Fig 2A-D). Fraction of bound sites described in Figure2C and D were calculated by taking the threshold for bound enrichment>1.5 versus all regions within the DHS bin indicated (DHS bins: less than 6, 6 to 7, 7 to 10, and more than 10).

For the analysis shown in Figure 2E and F, the numbers of cognate motif occurrences were contrasted between the consensus peaks and all other cognate motif regions with at least one cognate motif within the same DHS bins. In Figure 2G and H, consensus peaks and same number of background genomic regions with a single cognate motif were pooled and average TF ChIP enrichments on motifs grouped by the central nucleotide types (TA, GA, GC and GG central nucleotides as the most enriched types) were plotted across low to high initial DHS signal using loess curve and its standard error. This combines the ChIP enrichments and motif distributions across different initial accessibility on single motifs, revealing the relative binding potentials on the motif types as a function of the regions’ accessibility. The effect of motif strength for binding can also be seen by the effect of multiple occurrences, where comparison of two GC motifs versus single GA motif would give a similar curve relative to their respective backgrounds across different accessibilities (Figure S2G).

#### Identifying Key Chromatin Features for NGN2 Binding Prediction

To identify potential critical factors impacting the initial binding by NGN2, we sought to determine whether any other chromatin features from mESCs in addition to the chromatin accessibility are crucial for predicting NGN2 genome-wide binding. To this end, we generated a training set using NGN2 6h TF-binding peaks (MACS2) together with the same size background sequences from randomly selected 500 bp genomic tiles. Chromatin features for these genomic regions were quantified using qCount function from QuasR, including chromatin accessibility measured by DNase-seq, average DNA methylation, active histone marks (H3K4me3, H3K4me2, H3K4me1, H3K27ac, H3K36me3), and repressive histone marks (H3K9me3, H3K9me2, H3K27me3) from undifferentiated mESCs (publicly available datasets ^44,63–66^ listed in Supplementary Table 2). Furthermore, we analyzed genomic sequences for NGN2 motif features, calculating PWM log-odds scores and counting NGN2-enriched 10mers individually for each E-box motif core (CANNTG). We then used corrplot^67^ (v0.92) to visualize correlation patterns between these chromatin and sequence features and the NGN2 IP/GFP enrichment value where the features were grouped and ordered according to hierarchical clustering with the ward.D method. These chromatin features served as inputs for the training of the random forest^68^ model (R caret^69^, v6.0) with the parameters of 100 trees and 10 variables per tree to predict the continuous NGN2 IP/GFP enrichment (log2) values. We extracted feature importance scores using varImp function (scale=true) from caret^69^ package to reveal the most informative chromatin features for predicting NGN2 binding.

#### CNN training and model interpretation

Training of CNN models was performed using DNA sequences and DNaseI profiles at peak sites (i.e. TF binding sites) to predict the continuous values of ChIP-seq enrichment signal (log2 IP/GFP) for TF binding. Basset^70^ CNN architecture was implemented and further tailored based on design and hyperparameters from DeepSTARR^71^. The CNN architecture included four initial convolutional layers (1D, with filter counts of 512, 512, 512, and 256; kernel sizes of 12, 3, 5, and 3, respectively), each succeeded by ReLU activation and by max-pooling operations with pool size of 2. Following the convolutional sequence, our network architecture comprised two dense layers with 512 and 256 neurons, respectively. Two dense layers were both activated by ReLU and a dropout rate of 0.4 was applied to each. A linear activation function in the output layer was utilized to predict the TF ChIP–seq enrichment 6 hours post-induction. The model was developed within the Keras^72^ environment with the help of the KerasR^73^ package (v2.2.5.0) powered by a TensorFlow^74^ (v2.0.0) backend. Optimization was carried out using the Adam^75^ optimizer with a mean squared error loss function and a training batch size of 64. Early stopping was monitored, using a 20% validation split from the training dataset and a patience threshold of 15 epochs. Chromosomes 17, 18, and 19 were assigned for testing, omitted from the training and validation steps to assess model performance. TF binding sites from chromosomes 14, 15 and 16 were kept as the validation set to optimize the hyperparameters. As input to the CNN model, one-hot-encoded representation of 500 bp-long DNA sequence(s) were combined with pre-existing DNaseI^44^￼ that were generated with the qProfile function from QuasR package. DNaseI profiles were smoothed using a rolling mean with a window size of 21 bp and scaled from 0 to 1 to match the one-hot encoded range of the DNA sequence. For modelling the TF binding, we initially used a looser criterion to assemble a larger positive set (IDR < 0.2, average ChIP-seq enrichment over GFP >0.5 and over DNA binding mutant >0 for replicates at 6h). The foreground set (NGN2: 64,640, MyoD1: 50,964 sites) was supplemented with a background set of the same size, containing regions from the mouse genome with a similar DNaseI distribution. These background regions were selected using the matchRanges^76^￼ package (v1.10.0) in an iterative process to prevent redundant genomic matches. The CNN model was trained and evaluated on both the positive and negative strands of the genomic sites.

The same CNN architecture was adapted to predict i) accessibility changes as measured by ATAC-seq, ii) fold changes in H3K27ac, and iii) fold changes in Pol II signal intensities at multiple time points, specifically at 6 and 24 hours after TF induction. For NGN2 and MyoD1, and for each time point (6h and 24h) separately, we assembled a peak space that included the corresponding TF binding sites, ATAC-seq peaks and the GFP controls. Fold changes were calculated using cyclicLoess^40,41^ normalized values from the limma package^42^ (v3.60.3). For ATAC-seq changes, the CNN model was trained to predict both increases and decreases in accessibility regions, along with a balanced background showing no change in accessibility. For H3K27ac and Pol II, the CNN model was trained to predict only increasing sites together with a balanced set of background regions with no change, while regions with decreased signal intensity were discarded.

The models were evaluated on the held-out test set (chromosomes 17-19) using Pearson correlation and precision-recall curve metrics (area under the Precision-Recall Curve) calculated with R ROCR^77^ package (v1.0). The performance of the CNN model was compared to two alternative models: i) a linear model and ii) a random forest^68^ (number of trees:100, the number of variables: 8) approach both implemented using the R caret^69^ package (v6.0). Both alternatives utilized a single value of DNase I signal, and E-box motif counts based on overrepresented 10mers (separated by the core of the motif CANNTG) for each genomic region of 500 bp.

To interpret the CNN model^78^, we utilized the shap.DeepExplainer function (enhanced implementation of DeepLIFT^79^ algorithm) available from the SHAP library ^80,81^. This tool was used to assign contribution scores for each nucleotide in the sequences (±250 bp) for the reliable set of consensus TF binding sites. To serve as background sequence for the ‘DeepExplainer’, we generated 100 dinucleotide-shuffled versions for each TF binding site and all the values were set to zero for the background DNase I profile. To cluster and summarize the subsequences extracted from CNN model, hypothetical/contribution scores were fed into the TF-MoDISco^82^ tool using the parameters of sliding window size of 15 bp with additional flank size of bp 5 and a cut-off of target seqlet FDR of 0.15. TF-MoDISco consensus seqlets (metaclusters) with more than 100 instances were retained. For this TF-MoDISco step, contribution scores assigned to DNase I values were excluded and only the hypothetical/contribution scores for the one-hot encoded DNA sequence were considered. As TF-MoDISco revealed E-box motifs as the most important feature, we decided to compute “contribution weight matrix” (CWM) for E-box sites by taking the average of the contribution scores of the nucleotides present in the input sequence as similarly done by BPNet^83^ and affinity distillation^84^ approaches. From the consensus TF binding sites, 26,128 E-box instances were shown to have total contribution score greater than 0.1 for NGN2, and 25,327 E-box instances for MyoD1. At the same E-box sites, the contribution scores (i.e. also called as hypothetical scores) calculated for DNaseI profile were also summarized at base-pair resolution and shown as the letter ‘D’ in the sequence logo^62^. For exactly the same E-box sites, positional weight matrix was generated by calculating the frequency of each nucleotide for each position and visualized with the help of ggseqlogo^62^ (v0.2).

#### Estimation of marginal effects of E-box motif variants on TF binding strength and chromatin activation potential using in-silico mutagenesis

To isolate the impact of sequence features independent of the genomic sequence context and initial chromatin accessibility, affinity distillation approach was adapted from Alexandari *et al.*^84^ to estimate the binding strengths of E-box motif variants using synthetic sequences. First, we generated 200 random background sequences (500 bp) without any E-box motifs by dinucleotide shuffling a randomly selected set of consensus NGN2 binding sites. Next, we inserted variants of the E-box motif into these random backgrounds around the center (±10 bp). Trained CNN models were then applied to predict binding to the synthetic sequences and measure the relative increase in the binding strength for a specific motif variant by comparing to ‘no insert’ control. To ensure accuracy, all synthetic sequences were designed to ensure a single E-box motif site per sequence while eliminating synthetic sequences ending up with multiple E-box sites per sequence. Since the trained CNN model requires DNase I as input, affinity distillation method was repeated for various levels of uniform DNase I profiles ranging from 0 to 1 in increments of 0.1. To obtain a reliable estimate for the core variant of the CANNTG E-box motif, the core motif variants were tested in combination with the top 12 flank sequences selected for their relative binding strength derived from affinity distillation. The relative binding strength of the core sequence variants was then calculated by averaging the binding strengths of all the combinations of flanking motif variants. The same approach was also applied to interpret the CNN models trained for the prediction of ATAC-seq, H3K27ac and Pol-II change(s) during NGN2-and MyoD1-driven in vitro differentiation. Using the NGN2-binding CNN model, the impact of multiple E-box motif instances were evaluated on the TF binding prediction. For this analysis, a consensus NGN2 motif was inserted near the center (250±10 bp) of the sequence with up to two additional consensus NGN2 motifs would be placed on either side of the central motif (±10-20 bp).

#### In silico motif interaction analysis

In silico motif interaction analysis was conducted using the detailed methodology of Avsec *et al.*^83^ to investigate the influence of E-box motif pairs on the cooperative binding of NGN2 and MyoD1 interaction with PBX motif. Briefly, in the case of NGN2, two consensus E-box motifs were inserted into 200 random background sequences. One E-box motif (Motif A) was positioned near the center of each sequence, while the second motif (Motif B) was placed downstream at varying distances (d). The CNN TF binding model was used to predict binding enrichment (as in total signal intensity) for the motif pair (Motif-AB), for Motif A alone (Motif-A), for Motif B alone (Motif-B), and for sequences with no motifs (Ø). The cooperative impact of the motif pair was calculated by subtracting the marginal effect of Motif B and dividing by the binding prediction for Motif A alone, summarized as (MotifAB – (MotifB - Ø)) / MotifA. We then averaged the cooperativity estimates across 200 random background sequences. In the case of PBX-MyoD1 heterotypic interaction, MyoD1 consensus motif was embedded in the center and the PBX consensus motif was placed along the sequence at a range of distances.

To complement DL-based in-silico motif interaction analysis, we systematically analyzed NGN2 motif pairs in the genome to investigate the impact of inter-motif distance on binding probability. Specifically, E-box motif pairs were extracted from the mouse reference genome, focusing on NGN2-enriched 10-mers of CAGATG, CAGCTG, and CATATG. To ensure a robust analysis, pairs containing more than two motifs within a 200 bp window were excluded. The analysis targeted initially low accessible sites (log2 DNase-I signal <6.5), retaining 185,326 NGN2 motifs with neighboring motifs closer than 200 bp (center-to-center). The bound fraction of motifs (IP/GFP ChIP enrichment bound>1.5, unbound<0.5) was quantified as a function of distance between motif pairs to study how varying distances between NGN2 motif pairs influence their binding probabilities (Figure S3H). In addition, the enriched co-occurrence of TF motif pairs (enriched 10mers) within specified distances in consensus NGN2/MyoD1 binding sites was analyzed. To this end, we compared the frequency of E-box motif pairs in TF binding sites against random genomic tiles of 500 bp from the mouse genome. We aimed to determine if these motif pairs are overrepresented within specific distances (6-30 bp, 30-100 bp and 100-500 bp) in TF binding sites compared to random genomic regions. Statistical significance was assessed with Fisher’s exact test.

#### Nucleosome positions versus cognate motifs

The observations on the contribution scores in the CNN model for NGN2 binding in open chromatin regions indicate that the shape of the accessibility signal relative to the E-box may have an informative role in predicting the binding. Thus, we probed the underlying information that could help explain the presence of unbound strong motifs in open chromatin regions. First, NGN2 ChIP enrichments (IP/GFP) on NGN2 cognate motif sites (±250 bp) residing in mESC DHS>9 were quantified. Sites with enrichment above 1.5 and below 0.5 were taken as bound and unbound, respectively. MNase-seq and DNase-seq data in mESCs were quantified for these two groups centered on the NGN2 motifs using qProfile function from QuasR package with 41 bp sliding windows along (±500 bp) and no additional shifting of the reads, thus corresponding to the cut sites by these enzymes. Values of each region were normalized by median-centering and averaged across all regions. The resulting average MNase and DNase-seq shapes for these two groups were visualized as line plots in Figure 3G.

#### Structure modelling with AlphaFold3

The structure model predictions were performed using AlphaFold 3^85^ (available at https://alphafoldserver.com/ as of 2024-07-09) to model the structures of bHLH DNA-binding domains of NGN2 and TCF3 in complex with their respective DNA sequences. The following DNA sequences were used for the predictions: i) NGN2 homodimer: AAAAACATATGTTTTT, ii) TCF3 homodimer: AAAAACAGGTGTTTTT, iii) NGN2-TCF3 heterodimer: AAAAACAGATGTTTTT, iv) E-box motif pair separated by 11 bp (2xNGN2 homodimer): AAAAAACATATGTTTAACATATGTTTTTT and v) E-box motif pair separated by 11 base pairs separated by 15 bp (2xNGN2 homodimer): AAAAAACATATGTTTTTAAAACATATGTTTTTT.

#### Dimensionality reduction with UMAP

To integrate multiple measurements of the genome made in this study allowing visualization on the same set, we generated a combined peak set across experiments including i) ChIP-seq peaks of NGN2 and MyoD1 from 6h to 24h and ii) ATAC-seq peaks collected from mESCs and upon 6h and 24h NGN2 and MyoD1 induction. Genomic regions overlapping more than 70% were merged. Within these regions TF ChIP-seq (±250 bp), ATAC-seq (±500 bp) and histone marks (±1 kbp) were counted. ATAC-seq and histone mark counts were normalized with cyclicLoess^40,41^ available from the limma^42^ package (v3.60.3), allowing comparable fold change calculation across treatments. To visualize the relevant genomic regions of action, we retained genomic regions that met either of the following criteria: i) NGN2 or MyoD1 IP/GFP ChIP-seq enrichment is higher than 1.5 at any time point, or ii) a dynamic change in absolute accessibility FC greater than 0.5 at any time point upon NGN2 or MyoD1 induction.

We then carried out dimensionality reduction on these genomic regions using Uniform Manifold Approximation and Projection (UMAP) technique^86^ available via R umap package (v0.2.10.0) (with parameters: metric=”correlation”, min_dist=0.001, n_neighbors=201).

UMAP enables us to capture the continuous progression of genomic regions by grouping similar regions based on their chromatin dynamics. UMAP was used to color the genomic regions based on the signal intensity of TF ChIP-seq enrichments, initial DNase-seq accessibility from undifferentiated mESCs and fold changes of ATAC-seq, H3K27ac ChIP-seq, Pol-II ChIP-seq and RNA-seq along the differentiation.

#### Differential motif enrichment on changing TF binding patterns

To investigate the basis of change in NGN2 and MyoD1 binding patterns through differentiation, we conducted a motif enrichment analysis contrasting bound regions. For NGN2 and MyoD1 individually, 6h and 24h ChIP-seq peaks were pooled and peaks overlapping more than 70% were merged. ChIP-seq enrichments (IP/GFP) at 6h and 24 h inductions were calculated in the merged peak-set. 6h ChIP-seq enrichment values (log2 fold changes over GFP control) were subtracted from 24h ChIP enrichments. Subsequently, the peaks were categorized into bins based on the magnitude of change in transcription factor enrichment between these two time points. Differential motif enrichment analysis across these bins was performed with the monaLisa package using “otherBins” as the background and a minimum score of 10 matching to the JASPAR motif library. To refine the motif logos for visualization, motifs that were subsequences of each other or exhibited high similarity (based on the motifSimilarity score depending on the length of the motif) were consolidated with the motif logo displaying the highest differential enrichment at either time- point. (Figure S6C-D). The top two differentially enriched unique motif logos in 6h vs 24h were displayed in Figure 6C.

To investigate the relation between NGN2 binding and newly opened sites at EBF motifs, position of EBF motifs (JASPAR: MA0154.1/2/3 with matching score>10) overlapping with the newly opened regions (initial DHS<7 and 24h NGN2 vs 24h GFP ATAC-seq FC>0.5) upon NGN2 24h induction were identified. Distribution of NGN2 cognate motifs centered around the EBF motifs were visualized as density plots for the two subgroups of these newly opened EBF sites: the ones that gained new NGN2 binding (NGN2 24h enrichment>1.5 and NGN2 6h enrichment<0.5) versus the ones that are not bound by NGN2 (both NGN2 24h and 6h enrichment<0.5). For these subgroups, ATAC-seq meta-profiles of these regions centered around EBF motifs (±1kb) were generated using qProfile function in QuasR with 21 bp sliding windows (Figure 6E-F).

#### MyoD1 datasets from in vitro differentiated myocyte cells

MyoD1 binding sites (ENCODE identifier: ENCFF628LCC) derived from in vitro differentiated myocyte cells originated from mouse C2C12 cells were obtained from the ENCODE^66,87,88^ portal (https://www.encodeproject.org/) together with the alignment ChIP-seq datasets with the following identifiers: ENCFF816WPP, MyoD1 ChIP-seq at 24 hours and ENCFF204HMY as the corresponding control. ATAC-seq datasets^89^ for the undifferentiated C2C12 cells were downloaded from NCBI GEO^90^ from the identifiers GSM4889011 and GSM4889012. As similarly described for CNN training, the foreground set of MyoD1 binding sites (n: 29,477) were resized to 500 bp as the requirement of the CNN model and the test set was extended with a random background set of the same size. Quantile normalization was performed to align the DNaseI distribution of the test set from differentiated myocytes with that of the training set of MyoD1 binding sites derived from mESCs. Subsequently, the MyoD1-binding CNN model derived from mESCs was utilized to predict the enrichment levels of MyoD1 binding sites in in vitro myocyte cells differentiated from mouse C2C12 cells.

#### Enriched activity of the predicted NGN2/MyoD1 binding sites on ENCODE cCREs across tissues

ENCODE candidate cis-Regulatory Elements (cCREs, n = 343,731) combined from all cell types^66^ (ENCODE 2012) were downloaded from UCSC table browser^91^ (http://genome.ucsc.edu). cCREs were filtered for mappability and resized to 500 bp to feed into the CNN models. The cCRE sequences were analyzed using CNN models to predict the TF binding potential values at 6h induction of NGN2 and MyoD1 across various DNase I levels. A total of 302 DNase-seq datasets derived from mouse ‘primary cells’ or ‘tissues’ were collected from the ENCODE server^87^ in normalized bigwig format as listed in Supplementary Table 2. These datasets represent 52 different tissues including cerebellum, embryo, immune cells and many others. Constitutively present cCREs that showed an average signal greater than 7 and standard deviation lower than 1.5 were excluded from further analysis to focus on variable cCREs across tissues (n = 293,374). To identify cell types with high activity at the predicted TF binding sites, we followed these steps: i) all cCREs were ranked based on their potential TF binding at low accessibility ([0-0.3]) as predicted by CNN models, ii) in each ENCODE sample, the average ENCODE DNaseI signal at the top 5,000 cCRE sites was divided by the average signal of the bottom half of cCREs, iii) tissues were sorted based on their DNase I signal enrichment compared to the background with two top distinct tissues visualized as heatmaps (rightmost heatmap panels in Figure 7E and Figure 7D). As a reference, two out of 302 datasets closest to the background value of 1 were included in the heatmap for background comparison.

#### Predicting regulatory potential of GWAS variant(s) using CNN models

We systematically searched for GWAS SNPs to identify candidate E-box variant(s) with potential regulatory effects on gene expression. A comprehensive GWAS dataset (*P* value < 1e-5) was obtained from the NHGRI-EBI GWAS Catalog^92^ on 02/02/2024. After obtaining SNPs located within human cCREs, we overlapped these SNPs with cerebellum-specific eQTLs from the GTEX database^93^, given NGN2’s predicted activity in this tissue. Then, the systematic search was restricted to SNPs that alter the core of the E-box motif, which could affect NGN2 binding strength. Our analysis identified a single significant SNP, rs58130172, associated with ’Educational attainment’ trait^94,95^. This alternative variant, located near the TSS, was associated with increase in the expression of the DISP3 gene, acting as an eQTL with a q-value less than 4e−6 and a log2FC of 0.49^93^. Using our NGN2-associated CNN models derived from mouse ESCs, we analyzed the 500 bp region around rs58130172 to study the impact of single nucleotide variation on NGN2 binding, chromatin accessibility and H3K27ac levels.

#### Visualization of genomic enrichment signal using IGV and ComplexHeatmap

Genome-wide coverage tracks (tdf) were generated with IGVtools^96^ (Robinson 2011) count function in windows (-w) of 10 after the extension (-e) of reads to 150 for ChIP-seq and 0 as no extension for the rest of data types. Integrative genomics viewer^96,97^ (v2.11.3) was employed to visualize the normalized genome-wide tracks. To visualize the NGN2 and MyoD1 binding sites together in a heatmap, we merged the consensus TF binding sites if they overlapped by more than 70%. Genomic signal profiles for these TF binding sites were generated by counting the 5′ positions of mapped reads using the qProfile function of QuasR^13^ with the default parameters except for the ChIP-seq samples where reads were shifted based on the estimated size of the ChIP–seq library fragments. Normalized RPKM counts were then calculated for ±1 kb regions surrounding the TF binding sites, with each site smoothed using a running mean of 6 bp. For ChIP-seq datasets, GFP control was subtracted, and the replicates were averaged. Regions were ordered based on the differential ranking of the compared TF ChIP-seq enrichments. The resulting profiles were visualized with the EnrichedHeatmap package^98–100^ (v1.34.0). Enriched 10mers detected for NGN2 and MyoD1 were pooled together and all the motif instances from the mouse genome were assembled. For the target regions of interest (i.e. TF binding sites), the genomic bins overlapping with the motif instances were highlighted in black in the additional EnrichedHeatmap column. For the EnrichedHeatmap visualization in Figure 2I, NGN2 6h and MyoD1 6h consensus peaks were merged. TF binding sites were categorized into five groups based on the ChIP enrichment values of NGN2 (Nenr) and MyoD1: (i) ’2 N>M’ where (Nenr-Menr)> 0.75, (ii) ’4 M>N’ where (Menr - Nenr) > 0.75, (iii) ’3 N&M’ where Nenr > 1, Menr > 1, and |Nenr - Menr| < 0.75, (iv) ’1 N’ with Nenr > 1.5 and Menr < 0.75, and (v) ’5 M’ with Menr > 1.5 and Nenr < 0.75. Regions with enrichment values lower than 1 for both NGN2 and MyoD1 were excluded. These TF binding sites were further mapped to the nearest gene using the annotatePeak function from the ChIPseeker^36^ package (v1.40.0) based on the gene annotations from TxDb.Mmusculus.UCSC.mm10.knownGene (v3.10.0). Next, we characterized the resulting gene sets using gene annotation enrichment analysis via the clusterProfiler^34^ package (v4.12.0) using GO:BP^35^ and Hallmark gene sets^101,102^ from the mouse MsigDB collection^32^ (Figure S2J).

For the EnrichedHeatmap visualization in Figure S6A, NGN2 ChIP-seq peaks showing greater than 1.5-fold enrichment at any time-point of 6h, 12h and 24h NGN2 induction within a time-course experiment were merged and the genomic sites were ordered in the heatmap based on the differential ranking between 6h and 24h enrichments. For the EnrichedHeatmap visualization in Figure 7A, MyoD1 6h mESC consensus peaks and MyoD1 24h C2C12 consensus peaks were merged. Subset of these regions, showing enrichment above 1 in both cell-types, were visualized in Figure S7A.

## REFERENCES

1. Wunderlich, Z., and Mirny, L.A. (2009). Different gene regulation strategies revealed by analysis of binding motifs. Trends Genet 25, 434–440. 10.1016/J.TIG.2009.08.003.

2. Ephrussi, A., Church, G.M., Tonegawa, S., and Gilbert, W. (1985). B lineage--specific interactions of an immunoglobulin enhancer with cellular factors in vivo. Science 227, 134–140. 10.1126/SCIENCE.3917574.

3. Longo, A., Guanga, G.P., and Rose, R.B. (2008). Crystal structure of E47- NeuroD1/beta2 bHLH domain-DNA complex: heterodimer selectivity and DNA recognition. Biochemistry 47, 218–229. 10.1021/BI701527R.

4. De Martin, X., Sodaei, R., Santpere, G., and Margaglione, M. (2021). Molecular Sciences Mechanisms of Binding Specificity among bHLH Transcription Factors. 10.3390/ijms22179150.

5. Guo, J., Li, T., Schipper, J., Nilson, K.A., Fordjour, F.K., Cooper, J.J., Gordân, R., and Price, D.H. (2014). Sequence specificity incompletely defines the genome-wide occupancy of Myc. Genome Biol 15, 482. 10.1186/S13059-014-0482-3.

6. Casey, B.H., Kollipara, R.K., Pozo, K., and Johnson, J.E. (2018). Intrinsic DNA binding properties demonstrated for lineage-specifying basic helix-loop-helix transcription factors. 10.1101/gr.224360.117.

7. Srivastava, D., and Mahony, S. (2020). Sequence and chromatin determinants of transcription factor binding and the establishment of cell type-specific binding patterns. Biochim Biophys Acta Gene Regul Mech 1863. 10.1016/j.bbagrm.2019.194443.

8. Sönmezer, C., Kleinendorst, R., Imanci, D., Barzaghi, G., Villacorta, L., Schübeler, D., Benes, V., Molina, N., and Krebs, A.R. (2021). Molecular Co-occupancy Identifies Transcription Factor Binding Cooperativity In Vivo. Mol Cell 81, 255–267.e6. 10.1016/J.MOLCEL.2020.11.015.

9. Swinstead, E.E., Miranda, T.B., Paakinaho, V., Baek, S., Goldstein, I., Hawkins, M., Karpova, T.S., Ball, D., Mazza, D., Lavis, L.D., et al. (2016). Steroid Receptors Reprogram FoxA1 Occupancy through Dynamic Chromatin Transitions. Cell 165, 593–605. 10.1016/J.CELL.2016.02.067.

10. Isbel, L., Iskar, M., Durdu, S., Weiss, J., Grand, R.S., Hietter-Pfeiffer, E., Kozicka, Z., Michael, A.K., Burger, L., Thomä, N.H., et al. (2023). Readout of histone methylation by Trim24 locally restricts chromatin opening by p53. Nat Struct Mol Biol 30, 948–957. 10.1038/S41594-023-01021-8.

11. Hansen, J.L., and Cohen, B.A. (2022). A quantitative metric of pioneer activity reveals that HNF4A has stronger in vivo pioneer activity than FOXA1. Genome Biol 23. 10.1186/S13059-022-02792-X.

12. Isbel, L., Grand, R.S., and Schübeler, D. (2022). Generating specificity in genome regulation through transcription factor sensitivity to chromatin. Nat Rev Genet 23, 728–740. 10.1038/S41576-022-00512-6.

13. Slattery, M., Zhou, T., Yang, L., Dantas Machado, A.C., Gordân, R., and Rohs, R. (2014). Absence of a simple code: how transcription factors read the genome. Trends Biochem Sci 39, 381. 10.1016/J.TIBS.2014.07.002.

14. Kumar, N., Tsai, Y.H., Chen, L., Zhou, A., Banerjee, K.K., Saxena, M., Huang, S., Toke, N.H., Xing, J., Shivdasani, R.A., et al. (2019). The lineage-specific transcription factor CDX2 navigates dynamic chromatin to control distinct stages of intestine development. Development 146. 10.1242/DEV.172189.

15. Saotome, M., Poduval, D.B., Grimm, S.A., Nagornyuk, A., Gunarathna, S., Shimbo, T., Wade, P.A., and Takaku, M. (2024). Genomic transcription factor binding site selection is edited by the chromatin remodeling factor CHD4. Nucleic Acids Res 52, 3607–3622. 10.1093/nar/gkae025.

16. Aydin, B., Kakumanu, A., Rossillo, M., Moreno-Estellés, M., Garipler, G., Ringstad, N., Flames, N., Mahony, S., and Mazzoni, E.O. (2019). Proneural factors Ascl1 and Neurog2 contribute to neuronal subtype identities by establishing distinct chromatin landscapes. Nat Neurosci 22, 897–908. 10.1038/S41593-019-0399-Y.

17. Soufi, A., Donahue, G., and Zaret, K.S. (2012). Facilitators and impediments of the pluripotency reprogramming factors’ initial engagement with the genome. Cell 151, 994–1004. 10.1016/j.cell.2012.09.045.

18. Lee, Q.Y., Mall, M., Chanda, S., Zhou, B., Sharma, K.S., Schaukowitch, K., Adrian- Segarra, J.M., Grieder, S.D., Kareta, M.S., Wapinski, O.L., et al. (2020). Pro-neuronal activity of Myod1 due to promiscuous binding to neuronal genes. Nat Cell Biol 22, 401–411. 10.1038/S41556-020-0490-3.

19. Thoma, E.C., Wischmeyer, E., Offen, N., Maurus, K., Sirén, A.L., Schartl, M., and Wagner, T.U. (2012). Ectopic expression of neurogenin 2 alone is sufficient to induce differentiation of embryonic stem cells into mature neurons. PLoS One 7. 10.1371/JOURNAL.PONE.0038651.

20. Weintraub, H., Tapscott, S.J., Davis, R.L., Thayer, M.J., Adam, M.A., Lassar, A.B., and Miller, A.D. (1989). Activation of muscle-specific genes in pigment, nerve, fat, liver, and fibroblast cell lines by forced expression of MyoD. Proc Natl Acad Sci U S A 86, 5434–5438. 10.1073/PNAS.86.14.5434.

21. Davis, R.L., Weintraub, H., and Lassar, A.B. (1987). Expression of a single transfected cDNA converts fibroblasts to myoblasts. Cell 51, 987–1000. 10.1016/0092-8674(87)90585-X.

22. Baubec, T., Ivánek, R., Lienert, F., and Schübeler, D. (2013). Methylation-dependent and -independent genomic targeting principles of the MBD protein family. Cell 153, 480–492. 10.1016/J.CELL.2013.03.011.

23. Machlab, D., Burger, L., Soneson, C., Rijli, F.M., Schübeler, D., and Stadler, M.B. (2022). monaLisa: an R/Bioconductor package for identifying regulatory motifs. Bioinformatics 38, 2624–2625. 10.1093/BIOINFORMATICS/BTAC102.

24. Heinz, S., Benner, C., Spann, N., Bertolino, E., Lin, Y.C., Laslo, P., Cheng, J.X., Murre, C., Singh, H., and Glass, C.K. (2010). Simple combinations of lineage- determining transcription factors prime cis-regulatory elements required for macrophage and B cell identities. Mol Cell 38, 576–589. 10.1016/J.MOLCEL.2010.05.004.

25. Domcke, S., Bardet, A.F., Adrian Ginno, P., Hartl, D., Burger, L., and Schübeler, D. (2015). Competition between DNA methylation and transcription factors determines binding of NRF1. Nature 2015 528:7583 *528*, 575–579. 10.1038/nature16462.

26. Xin, B., and Rohs, R. (2018). Relationship between histone modifications and transcription factor binding is protein family specific. Genome Res 28, 321–333. 10.1101/GR.220079.116.

27. Kelley, D.R., Snoek, J., and Rinn, J.L. (2016). Basset: learning the regulatory code of the accessible genome with deep convolutional neural networks. Genome Res 26, 990–999. 10.1101/GR.200535.115.

28. Shrikumar, A., Greenside, P., and Kundaje, A. (2017). Learning Important Features Through Propagating Activation Differences. 34th International Conference on Machine Learning, ICML 2017 *7*, 4844–4866. arXiv.1605.01713.

29. Novakovsky, G., Dexter, N., Libbrecht, M.W., Wasserman, W.W., and Mostafavi, S. (2023). Obtaining genetics insights from deep learning via explainable artificial intelligence. Nat Rev Genet 24, 125–137. 10.1038/S41576-022-00532-2.

30. Shrikumar, A., Tian, K., Avsec, Ž., Shcherbina, A., Banerjee, A., Sharmin, M., Nair, S., and Kundaje, A. (2018). Technical Note on Transcription Factor Motif Discovery from Importance Scores (TF-MoDISco) version 0.5.6.5. 10.48550/arXiv.1811.00416.

31. Fong, A.P., Yao, Z., Zhong, J.W., Johnson, N.M., Farr, G.H., Maves, L., and Tapscott, S.J. (2015). Conversion of MyoD to a neurogenic factor: binding site specificity determines lineage. Cell Rep 10, 1937–1946. 10.1016/J.CELREP.2015.02.055.

32. Alexandari, A.M., Horton, C.A., Shrikumar, A., Shah, N., Li, E., Weilert, M., Pufall, M.A., Zeitlinger, J., Fordyce, P.M., and Kundaje, A. (2023). De novo distillation of thermodynamic affinity from deep learning regulatory sequence models of in vivo protein-DNA binding. bioRxiv. 10.1101/2023.05.11.540401.

33. Michael, A.K., Stoos, L., Crosby, P., Eggers, N., Nie, X.Y., Makasheva, K., Minnich, M., Healy, K.L., Weiss, J., Kempf, G., et al. (2023). Cooperation between bHLH transcription factors and histones for DNA access. Nature 619, 385–393. 10.1038/s41586-023-06282-3.

34. Barisic, D., Stadler, M.B., Iurlaro, M., and Schübeler, D. (2019). Mammalian ISWI and SWI/SNF selectively mediate binding of distinct transcription factors. Nature 569, 136–140. 10.1038/S41586-019-1115-5.

35. Kornberg, R.D., and Stryer, L. (1988). Statistical distributions of nucleosomes: nonrandom locations by a stochastic mechanism. Nucleic Acids Res 16, 6677–6690. 10.1093/NAR/16.14.6677.

36. D’Oliveira Albanus, R., Kyono, Y., Hensley, J., Varshney, A., Orchard, P., Kitzman, J.O., and Parker, S.C.J. (2021). Chromatin information content landscapes inform transcription factor and DNA interactions. Nat Commun 12. 10.1038/S41467-021-21534-4.

37. Zhong, J., Luo, K., Winter, P.S., Crawford, G.E., Iversen, E.S., and Hartemink, A.J. (2016). Mapping nucleosome positions using DNase-seq. Genome Res 26, 351–364. 10.1101/GR.195602.115.

38. Avsec, Ž., Weilert, M., Shrikumar, A., Krueger, S., Alexandari, A., Dalal, K., Fropf, R., McAnany, C., Gagneur, J., Kundaje, A., et al. (2021). Base-resolution models of transcription-factor binding reveal soft motif syntax. Nat Genet 53, 354–366. 10.1038/S41588-021-00782-6.

39. Abramson, J., Adler, J., Dunger, J., Evans, R., Green, T., Pritzel, A., Ronneberger, O., Willmore, L., Ballard, A.J., Bambrick, J., et al. (2024). Accurate structure prediction of biomolecular interactions with AlphaFold 3. Nature 630, 493–500. 10.1038/S41586-024-07487-W.

40. Calo, E., and Wysocka, J. (2013). Modification of enhancer chromatin: what, how and why? Mol Cell 49, 825–837. 10.1016/J.MOLCEL.2013.01.038.

41. Hartl, D., Krebs, A.R., Grand, R.S., Baubec, T., Isbel, L., Wirbelauer, C., Burger, L., and Schübeler, D. (2019). CG dinucleotides enhance promoter activity independent of DNA methylation. Genome Res 29, 554–563. 10.1101/GR.241653.118.

42. 42. Pan, Y., van der Watt, P.J., and Kay, S.A. (2023). E-box binding transcription factors in cancer. Front Oncol 13. 10.3389/FONC.2023.1223208.

43. Rauluseviciute, I., Riudavets-Puig, R., Blanc-Mathieu, R., Castro-Mondragon, J.A., Ferenc, K., Kumar, V., Lemma, R.B., Lucas, J., Chèneby, J., Baranasic, D., et al. (2024). JASPAR 2024: 20thãnniversary of the open-access database of transcription factor binding profiles. Nucleic Acids Res 52, D174–D182. 10.1093/NAR/GKAD1059.

44. Jones, S. (2004). An overview of the basic helix-loop-helix proteins. Genome Biol 5. 10.1186/GB-2004-5-6-226.

45. Bartlett, A., O’Malley, R.C., Huang, S.S.C., Galli, M., Nery, J.R., Gallavotti, A., and Ecker, J.R. (2017). Mapping genome-wide transcription-factor binding sites using DAP-seq. Nat Protoc 12, 1659–1672. 10.1038/NPROT.2017.055.

46. 46. De Souza, N. (2012). The ENCODE project. Nature Methods 2012 9:11 *9*, 1046– 1046. 10.1038/nmeth.2238.

47. Asp, P., Blum, R., Vethantham, V., Parisi, F., Micsinai, M., Cheng, J., Bowman, C., Kluger, Y., and Dynlacht, B.D. (2011). Genome-wide remodeling of the epigenetic landscape during myogenic differentiation. Proc Natl Acad Sci U S A 108. 10.1073/pnas.1102223108.

48. 48. Esteves de Lima, J., Bou Akar, R., Machado, L., Li, Y., Drayton-Libotte, B., Dilworth, F.J., and Relaix, F. (2021). HIRA stabilizes skeletal muscle lineage identity. Nat Commun 12. 10.1038/S41467-021-23775-9.

49. Kagda, M.S., Lam, B., Litton, C., Small, C., Sloan, C.A., Spragins, E., Tanaka, F., Whaling, I., Gabdank, I., Youngworth, I., et al. (2023). Data navigation on the ENCODE portal. 10.21203/rs.3.rs-3088639/v1.

50. Grounds, M.D., Garrett, K.L., and Beilharz, M.W. (1992). The transcription of MyoD1 and myogenin genes in thymic cells in vivo. Exp Cell Res 198, 357–361. 10.1016/0014-4827(92)90391-K.

51. Ragazzini, R., Boeing, S., Zanieri, L., Green, M., D’Agostino, G., Bartolovic, K., Agua- Doce, A., Greco, M., Watson, S.A., Batsivari, A., et al. (2023). Defining the identity and the niches of epithelial stem cells with highly pleiotropic multilineage potency in the human thymus. Dev Cell 58, 2428–2446.e9. 10.1016/J.DEVCEL.2023.08.017.

52. Aguet, F., Barbeira, A.N., Bonazzola, R., Brown, A., Castel, S.E., Jo, B., Kasela, S., Kim-Hellmuth, S., Liang, Y., Oliva, M., et al. (2020). The GTEx Consortium atlas of genetic regulatory effects across human tissues. Science 369, 1318–1330. 10.1126/SCIENCE.AAZ1776.

53. Lee, J.J., Wedow, R., Okbay, A., Kong, E., Maghzian, O., Zacher, M., Nguyen-Viet, T.A., Bowers, P., Sidorenko, J., Karlsson Linnér, R., et al. (2018). Gene discovery and polygenic prediction from a 1.1-million-person GWAS of educational attainment. Nat Genet 50, 1112. 10.1038/S41588-018-0147-3.

54. Konířová, J., Oltová, J., Corlett, A., Kopycińska, J., Kolář, M., Bartůněk, P., and Zíková, M. (2017). Modulated DISP3/PTCHD2 expression influences neural stem cell fate decisions. Sci Rep 7. 10.1038/SREP41597.

55. 55. Arntfield, M.E., and van der Kooy, D. (2011). β-Cell evolution: How the pancreas borrowed from the brain: The shared toolbox of genes expressed by neural and pancreatic endocrine cells may reflect their evolutionary relationship. Bioessays 33, 582–587. 10.1002/BIES.201100015.

56. Pataskar, A., Jung, J., Smialowski, P., Noack, F., Calegari, F., Straub, T., and Tiwari, V.K. (2016). NeuroD1 reprograms chromatin and transcription factor landscapes to induce the neuronal program. EMBO J 35, 24–45. 10.15252/embj.201591206.

57. 57. Birkhoff, J.C., Korporaal, A.L., Brouwer, R.W.W., Nowosad, K., Milazzo, C., Mouratidou, L., van den Hout, M.C.G.N., van IJcken, W.F.J., Huylebroeck, D., and Conidi, A. (2023). Zeb2 DNA-Binding Sites in Neuroprogenitor Cells Reveal Autoregulation and Affirm Neurodevelopmental Defects, Including in Mowat-Wilson Syndrome. Genes (Basel) 14. 10.3390/GENES14030629/S1.

58. Wang, R., Bhatt, A.B., Minden-Birkenmaier, B.A., Travis, O.K., Tiwari, S., Jia, H., Rosikiewicz, W., Martinot, O., Childs, E., Loesch, R., et al. (2023). ZBTB18 restricts chromatin accessibility and prevents transcriptional adaptations that drive metastasis. Sci Adv 9. 10.1126/sciadv.abq3951.

59. Lu, C., Garipler, G., Dai, C., Roush, T., Salome-Correa, J., Martin, A., Liscovitch- Brauer, N., Mazzoni, E.O., and Sanjana, N.E. (2023). Essential transcription factors for induced neuron differentiation. Nature Communications 2023 14:1 *14*, 1–14. 10.1038/s41467-023-43602-7.

60. Londhe, P., and Davie, J.K. (2011). Sequential association of myogenic regulatory factors and E proteins at muscle-specific genes. Skelet Muscle 1, 1–18. 10.1186/2044-5040-1-14.

## Method References

1. Mohn, F., Weber, M., Rebhan, M., Roloff, T.C., Richter, J., Stadler, M.B., Bibel, M., and Schübeler, D. (2008). Lineage-specific polycomb targets and de novo DNA methylation define restriction and potential of neuronal progenitors. Mol Cell 30, 755–766. 10.1016/J.MOLCEL.2008.05.007.

2. Lienert, F., Wirbelauer, C., Som, I., Dean, A., Mohn, F., and Schübeler, D. (2011). Identification of genetic elements that autonomously determine DNA methylation states. Nat Genet 43, 1091–1097. 10.1038/NG.946.

3. Baubec, T., Ivánek, R., Lienert, F., and Schübeler, D. (2013). Methylation-dependent and - independent genomic targeting principles of the MBD protein family. Cell 153, 480–492. 10.1016/J.CELL.2013.03.011.

4. Kaluscha, S., Domcke, S., Wirbelauer, C., Stadler, M.B., Durdu, S., Burger, L., and Schübeler, D. (2022). Evidence that direct inhibition of transcription factor binding is the prevailing mode of gene and repeat repression by DNA methylation. Nat Genet 54, 1895– 1906. 10.1038/S41588-022-01241-6.

5. Kaluscha, S., Domcke, S., Wirbelauer, C., Stadler, M.B., Durdu, S., Burger, L., and Schübeler, D. (2022). Evidence that direct inhibition of transcription factor binding is the prevailing mode of gene and repeat repression by DNA methylation. nature.comPaperpileS Kaluscha, S Domcke, C Wirbelauer, MB Stadler, S Durdu, L Burger, D SchübelerNature genetics, 2022•nature.comSign in *54*, 1895–1906. 10.1038/s41588-022-01241-6.

6. Thoma, E.C., Wischmeyer, E., Offen, N., Maurus, K., Sirén, A.L., Schartl, M., and Wagner, T.U. (2012). Ectopic expression of neurogenin 2 alone is sufficient to induce differentiation of embryonic stem cells into mature neurons. PLoS One 7. 10.1371/JOURNAL.PONE.0038651.

7. Wilkinson, G., Dennis, D., and Schuurmans, C. (2013). Proneural genes in neocortical development. Neuroscience 253, 256–273. 10.1016/J.NEUROSCIENCE.2013.08.029.

8. Schindelin, J., Arganda-Carreras, I., Frise, E., Kaynig, V., Longair, M., Pietzsch, T., Preibisch, S., Rueden, C., Saalfeld, S., Schmid, B., et al. (2012). Fiji: an open-source platform for biological-image analysis. Nat Methods *9*, 676–682. 10.1038/NMETH.2019.

9. Sato, Y., Nakajima, S., Shiraga, N., Atsumi, H., Yoshida, S., Koller, T., Gerig, G., and Kikinis, R. (1998). Three-dimensional multi-scale line filter for segmentation and visualization of curvilinear structures in medical images. Med Image Anal 2, 143–168. 10.1016/S1361-8415(98)80009-1.

10. Schneider, C.A., Rasband, W.S., and Eliceiri, K.W. (2012). NIH Image to ImageJ: 25 years of image analysis. Nat Methods 9, 671–675. 10.1038/NMETH.2089.

11. Weber, M., Hellmann, I., Stadler, M.B., Ramos, L., Pääbo, S., Rebhan, M., and Schübeler, D. (2007). Distribution, silencing potential and evolutionary impact of promoter DNA methylation in the human genome. nature.comPaperpileM Weber, I Hellmann, MB Stadler, L Ramos, S Pääbo, M Rebhan, D SchübelerNature genetics, 2007•nature.comSign in. 10.1038/ng1990.

12. Langmead, B., and Salzberg, S.L. (2012). Fast gapped-read alignment with Bowtie 2. Nat Methods 9, 357–359. 10.1038/NMETH.1923.

13. Gaidatzis, D., Lerch, A., Hahne, F., and Stadler, M.B. (2015). QuasR: quantification and annotation of short reads in R. Bioinformatics 31, 1130–1132. 10.1093/BIOINFORMATICS/BTU781.

14. Zhang, Y., Liu, T., Meyer, C.A., Eeckhoute, J., Johnson, D.S., Bernstein, B.E., Nussbaum, C., Myers, R.M., Brown, M., Li, W., et al. (2008). Model-based analysis of ChIP-Seq (MACS). Genome Biol 9. 10.1186/GB-2008-9-9-R137.

15. Grand, R.S., Burger, L., Gräwe, C., Michael, A.K., Isbel, L., Hess, D., Hoerner, L., Iesmantavicius, V., Durdu, S., Pregnolato, M., et al. (2021). BANP opens chromatin and activates CpG-island-regulated genes. Nature 596, 133–137. 10.1038/S41586-021-03689-8.

16. Amemiya, H.M., Kundaje, A., and Boyle, A.P. (2019). The ENCODE Blacklist: Identification of Problematic Regions of the Genome. Sci Rep 9. 10.1038/S41598-019-45839-Z.

17. Stark, R., and Brown, G. (2011). DiffBind: differential binding analysis of ChIP-Seq peak data. R package.

18. Ross-Innes, C.S., Stark, R., Teschendorff, A.E., Holmes, K.A., Ali, H.R., Dunning, M.J., Brown, G.D., Gojis, O., Ellis, I.O., Green, A.R., et al. (2012). Differential oestrogen receptor binding is associated with clinical outcome in breast cancer. Nature 481, 389–393. 10.1038/NATURE10730.

19. R Core Team. (2023). R: A language and environment for statistical computing. R Foundation for Statistical Computing, Vienna, Austria. Open J Stat 13.

20. Li, Q., Brown, J.B., Huang, H., and Bickel, P.J. (2011). Measuring reproducibility of high-throughput experiments. Ann Appl Stat 5, 1752–1779. 10.1214/11-AOAS466.

21. Isbel, L., Iskar, M., Durdu, S., Weiss, J., Grand, R.S., Hietter-Pfeiffer, E., Kozicka, Z., Michael, A.K., Burger, L., Thomä, N.H., et al. (2023). Readout of histone methylation by Trim24 locally restricts chromatin opening by p53. Nat Struct Mol Biol 30, 948–957. 10.1038/S41594-023-01021-8.

22. Lawrence, M., Huber, W., Pagès, H., Aboyoun, P., Carlson, M., Gentleman, R., Morgan, M.T., and Carey, V.J. (2013). Software for Computing and Annotating Genomic Ranges. PLoS Comput Biol 9, 1003118. 10.1371/JOURNAL.PCBI.1003118.

23. Dobin, A., Davis, C.A., Schlesinger, F., Drenkow, J., Zaleski, C., Jha, S., Batut, P., Chaisson, M., and Gingeras, T.R. (2013). STAR: ultrafast universal RNA-seq aligner. Bioinformatics 29, 15–21. 10.1093/BIOINFORMATICS/BTS635.

24. Li, H., Handsaker, B., Wysoker, A., Fennell, T., Ruan, J., Homer, N., Marth, G., Abecasis, G., and Durbin, R. (2009). The Sequence Alignment/Map format and SAMtools. Bioinformatics 25, 2078. 10.1093/BIOINFORMATICS/BTP352.

25. Liao, Y., Smyth, G.K., and Shi, W. (2019). The R package Rsubread is easier, faster, cheaper and better for alignment and quantification of RNA sequencing reads. Nucleic Acids Res 47, e47. 10.1093/NAR/GKZ114.

26. Liao, Y., Smyth, G.K., and Shi, W. (2014). featureCounts: an efficient general purpose program for assigning sequence reads to genomic features. Bioinformatics 30, 923–930. 10.1093/BIOINFORMATICS/BTT656.

27. Gentleman, R.C., Carey, V.J., Bates, D.M., Bolstad, B., Dettling, M., Dudoit, S., Ellis, B., Gautier, L., Ge, Y., Gentry, J., et al. (2004). Bioconductor: open software development for computational biology and bioinformatics. Genome Biol 5, R80. 10.1186/GB-2004-5-10-R80.

28. Frankish, A., Diekhans, M., Jungreis, I., Lagarde, J., Loveland, J.E., Mudge, J.M., Sisu, C., Wright, J.C., Armstrong, J., Barnes, I., et al. (2021). GENCODE 2021. Nucleic Acids Res 49, D916–D923. 10.1093/NAR/GKAA1087.

29. Robinson, M.D., McCarthy, D.J., and Smyth, G.K. (2010). edgeR: a Bioconductor package for differential expression analysis of digital gene expression data. Bioinformatics 26, 139. 10.1093/BIOINFORMATICS/BTP616.

30. Benjamini, Y., and Hochberg, Y. (1995). Controlling the False Discovery Rate: A Practical and Powerful Approach to Multiple Testing. J R Stat Soc Series B Stat Methodol 57. 10.1111/j.2517-6161.1995.tb02031.x.

31. Subramanian, A., Tamayo, P., Mootha, V.K., Mukherjee, S., Ebert, B.L., Gillette, M.A., Paulovich, A., Pomeroy, S.L., Golub, T.R., Lander, E.S., et al. (2005). Gene set enrichment analysis: a knowledge-based approach for interpreting genome-wide expression profiles. Proc Natl Acad Sci U S A 102, 15545–15550. 10.1073/PNAS.0506580102.

32. Castanza, A.S., Recla, J.M., Eby, D., Thorvaldsdóttir, H., Bult, C.J., and Mesirov, J.P. (2023). Extending support for mouse data in the Molecular Signatures Database (MSigDB). Nat Methods 20, 1619–1620. 10.1038/S41592-023-02014-7.

33. Korotkevich, G., Sukhov, V., Budin, N., Shpak, B., Artyomov, M.N., and Sergushichev, A. (2021). Fast gene set enrichment analysis. bioRxiv, 060012. 10.1101/060012.

34. Wu, T., Hu, E., Xu, S., Chen, M., Guo, P., Dai, Z., Feng, T., Zhou, L., Tang, W., Zhan, L., et al. (2021). clusterProfiler 4.0: A universal enrichment tool for interpreting omics data. Innovation (Cambridge (Mass.)) 2. 10.1016/J.XINN.2021.100141.

35. Aleksander, S.A., Balhoff, J., Carbon, S., Cherry, J.M., Drabkin, H.J., Ebert, D., Feuermann, M., Gaudet, P., Harris, N.L., Hill, D.P., et al. (2023). The Gene Ontology knowledgebase in 2023. Genetics 224. 10.1093/GENETICS/IYAD031.

36. Yu, G., Wang, L.G., and He, Q.Y. (2015). ChIPseeker: an R/Bioconductor package for ChIP peak annotation, comparison and visualization. Bioinformatics 31, 2382–2383. 10.1093/BIOINFORMATICS/BTV145.

37. Machlab, D., Burger, L., Soneson, C., Rijli, F.M., Schübeler, D., and Stadler, M.B. (2022). monaLisa: an R/Bioconductor package for identifying regulatory motifs. Bioinformatics 38, 2624–2625. 10.1093/BIOINFORMATICS/BTAC102.

38. Buenrostro, J.D., Giresi, P.G., Zaba, L.C., Chang, H.Y., and Greenleaf, W.J. (2013). Transposition of native chromatin for multimodal regulatory analysis and personal epigenomics. Nat Methods 10, 1213. 10.1038/NMETH.2688.

39. Martin, M. (2011). Cutadapt removes adapter sequences from high-throughput sequencing reads. EMBnet J 17, 10–12. 10.14806/EJ.17.1.200.

40. Ballman, K. V., Grill, D.E., Oberg, A.L., and Therneau, T.M. (2004). Faster cyclic loess: normalizing RNA arrays via linear models. Bioinformatics 20, 2778–2786. 10.1093/BIOINFORMATICS/BTH327.

41. Bolstad, B.M., Irizarry, R.A., Åstrand, M., and Speed, T.P. (2003). A comparison of normalization methods for high density oligonucleotide array data based on variance and bias. Bioinformatics 19, 185–193. 10.1093/BIOINFORMATICS/19.2.185.

42. Ritchie, M.E., Phipson, B., Wu, D., Hu, Y., Law, C.W., Shi, W., and Smyth, G.K. (2015). limma powers differential expression analyses for RNA-sequencing and microarray studies. Nucleic Acids Res 43, e47. 10.1093/NAR/GKV007.

43. Bartlett, A., O’Malley, R.C., Huang, S.S.C., Galli, M., Nery, J.R., Gallavotti, A., and Ecker, J.R. (2017). Mapping genome-wide transcription-factor binding sites using DAP-seq. Nat Protoc 12, 1659–1672. 10.1038/NPROT.2017.055.

44. Domcke, S., Bardet, A.F., Adrian Ginno, P., Hartl, D., Burger, L., and Schübeler, D. (2015). Competition between DNA methylation and transcription factors determines binding of NRF1. Nature 2015 528:7583 *528*, 575–579. 10.1038/nature16462.

45. Hartl, D., Krebs, A.R., Grand, R.S., Baubec, T., Isbel, L., Wirbelauer, C., Burger, L., and Schübeler, D. (2019). CG dinucleotides enhance promoter activity independent of DNA methylation. Genome Res 29, 554–563. 10.1101/GR.241653.118.

46. Dewari, P.S., Southgate, B., McCarten, K., Monogarov, G., O’duibhir, E., Quinn, N., Tyrer, A., Leitner, M.C., Plumb, C., Kalantzaki, M., et al. (2018). An efficient and scalable pipeline for epitope tagging in mammalian stem cells using Cas9 ribonucleoprotein. Elife 7. 10.7554/ELIFE.35069.

47. Ostapcuk, V., Mohn, F., Carl, S.H., Basters, A., Hess, D., Iesmantavicius, V., Lampersberger, L., Flemr, M., Pandey, A., Thomä, N.H., et al. (2018). Activity-dependent neuroprotective protein recruits HP1 and CHD4 to control lineage-specifying genes. Nature 557, 739–743. 10.1038/S41586-018-0153-8.

48. Durinck, S., Spellman, P.T., Birney, E., and Huber, W. (2009). Mapping identifiers for the integration of genomic datasets with the R/Bioconductor package biomaRt. Nat Protoc 4, 1184–1191. 10.1038/NPROT.2009.97.

49. 49. Durinck, S., Moreau, Y., Kasprzyk, A., Davis, S., De Moor, B., Brazma, A., and Huber, W. (2005). BioMart and Bioconductor: a powerful link between biological databases and microarray data analysis. Bioinformatics 21, 3439–3440. 10.1093/BIOINFORMATICS/BTI525.

50. Orsburn, B.C. (2021). Proteome Discoverer-A Community Enhanced Data Processing Suite for Protein Informatics. Proteomes 9. 10.3390/PROTEOMES9010015.

51. Eng, J.K., McCormack, A.L., and Yates, J.R. (1994). An approach to correlate tandem mass spectral data of peptides with amino acid sequences in a protein database. J Am Soc Mass Spectrom 5, 976–989. 10.1016/1044-0305(94)80016-2.

52. Cox, J., and Mann, M. (2008). MaxQuant enables high peptide identification rates, individualized p.p.b.-range mass accuracies and proteome-wide protein quantification. Nat Biotechnol 26, 1367–1372. 10.1038/NBT.1511.

53. Soneson, C., Iesmantavicius, V., Hess, D., Stadler, M.B., and Seebacher, J. (2023). einprot: flexible, easy-to-use, reproducible workflows for statistical analysis of quantitative proteomics data. J Open Source Softw 8, 5750. 10.21105/JOSS.05750.

54. Phipson, B., Lee, S., Majewski, I.J., Alexander, W.S., and Smyth, G.K. (2016). Robust hyperparameter estimation protects against hypervariable genes and improves power to detect differential expression. Ann Appl Stat 10, 946–963. 10.1214/16-AOAS920.

55. Makowski, M.M., Gräwe, C., Foster, B.M., Nguyen, N. V., Bartke, T., and Vermeulen, M. (2018). Global profiling of protein-DNA and protein-nucleosome binding affinities using quantitative mass spectrometry. Nat Commun 9. 10.1038/S41467-018-04084-0.

56. Tyanova, S., Temu, T., and Cox, J. (2016). The MaxQuant computational platform for mass spectrometry-based shotgun proteomics. Nat Protoc 11, 2301–2319. 10.1038/NPROT.2016.136.

57. Lambert, S.A., Jolma, A., Campitelli, L.F., Das, P.K., Yin, Y., Albu, M., Chen, X., Taipale, J., Hughes, T.R., and Weirauch, M.T. (2018). The Human Transcription Factors. Cell 172, 650– 665. 10.1016/J.CELL.2018.01.029.

58. Hammelman, J., Patel, T., Closser, M., Wichterle, H., and Gifford, D. (2022). Ranking reprogramming factors for cell differentiation. Nat Methods 19, 812–822. 10.1038/S41592-022-01522-2.

59. Sandelin, A., Alkema, W., Engström, P., Wasserman, W.W., and Lenhard, B. (2004). JASPAR: an open-access database for eukaryotic transcription factor binding profiles. Nucleic Acids Res 32, D91. 10.1093/NAR/GKH012.

60. 60. Fornes, O., Castro-Mondragon, J.A., Khan, A., Van Der Lee, R., Zhang, X., Richmond, P.A., Modi, B.P., Correard, S., Gheorghe, M., Baranašić, D., et al. (2020). JASPAR 2020: update of the open-access database of transcription factor binding profiles. Nucleic Acids Res 48, D87–D92. 10.1093/NAR/GKZ1001.

61. Heinz, S., Benner, C., Spann, N., Bertolino, E., Lin, Y.C., Laslo, P., Cheng, J.X., Murre, C., Singh, H., and Glass, C.K. (2010). Simple combinations of lineage-determining transcription factors prime cis-regulatory elements required for macrophage and B cell identities. Mol Cell 38, 576–589. 10.1016/J.MOLCEL.2010.05.004.

62. Wagih, O. (2017). ggseqlogo: a versatile R package for drawing sequence logos. Bioinformatics 33, 3645–3647. 10.1093/BIOINFORMATICS/BTX469.

63. 63. Tippmann, S.C., Ivanek, R., Gaidatzis, D., Schöler, A., Hoerner, L., Van Nimwegen, E., Stadler, P.F., Stadler, M.B., and Schübeler, D. (2012). Chromatin measurements reveal contributions of synthesis and decay to steady-state mRNA levels. Mol Syst Biol 8. 10.1038/MSB.2012.23.

64. Stadler, M.B., Murr, R., Burger, L., Ivanek, R., Lienert, F., Schöler, A., Wirbelauer, C., Oakeley, E.J., Gaidatzis, D., Tiwari, V.K., et al. (2011). DNA-binding factors shape the mouse methylome at distal regulatory regions. Nature 480, 490–495. 10.1038/NATURE10716.

65. 65. von Meyenn, F., Iurlaro, M., Habibi, E., Liu, N.Q., Salehzadeh-Yazdi, A., Santos, F., Petrini, E., Milagre, I., Yu, M., Xie, Z., et al. (2016). Impairment of DNA Methylation Maintenance Is the Main Cause of Global Demethylation in Naive Embryonic Stem Cells. Mol Cell 62, 848– 861. 10.1016/J.MOLCEL.2016.04.025.

66. 66. De Souza, N. (2012). The ENCODE project. Nature Methods 2012 9:11 *9*, 1046–1046. 10.1038/nmeth.2238.

67. Wei, T., Simko, V., Levy, M., Xie, Y., Jin, Y.J., and Zemla, J. (2017). Visualization of a Correlation Matrix. R package “corrplot”. Statistician 56.

68. Breiman, L. (2001). Random forests. Mach Learn 45, 5–32. 10.1023/A:1010933404324/METRICS.

69. Kuhn, M. (2008). Building Predictive Models in R Using the caret Package. J Stat Softw 28, 1–26. 10.18637/JSS.V028.I05.

70. Kelley, D.R., Snoek, J., and Rinn, J.L. (2016). Basset: learning the regulatory code of the accessible genome with deep convolutional neural networks. Genome Res 26, 990–999. 10.1101/GR.200535.115.

71. de Almeida, B.P., Reiter, F., Pagani, M., and Stark, A. (2022). DeepSTARR predicts enhancer activity from DNA sequence and enables the de novo design of synthetic enhancers. Nat Genet 54, 613–624. 10.1038/S41588-022-01048-5.

72. Chollet, F. (2015). keras. Preprint.

73. Arnold, T.B. (2017). kerasR: R Interface to the Keras Deep Learning Library. J Open Source Softw 2, 296. 10.21105/JOSS.00296.

74. Abadi, M., Agarwal, A., Barham, P., Brevdo, E., Chen, Z., Citro, C., Corrado, G.S., Davis, A., Dean, J., Devin, M., et al. (2016). TensorFlow: Large-Scale Machine Learning on Heterogeneous Distributed Systems. arXiv preprint arXiv:1603.04467 172.

75. 75. Kingma, D.P., and Ba, J.L. (2014). Adam: A Method for Stochastic Optimization. 3rd International Conference on Learning Representations, ICLR 2015 - Conference Track Proceedings.

76. Davis, E.S., Mu, W., Lee, S., Dozmorov, M.G., Love, M.I., and Phanstiel, D.H. (2023). matchRanges: generating null hypothesis genomic ranges via covariate-matched sampling. Bioinformatics 39. 10.1093/BIOINFORMATICS/BTAD197.

77. Sing, T., Sander, O., Beerenwinkel, N., and Lengauer, T. (2005). ROCR: visualizing classifier performance in R. Bioinformatics 21, 3940–3941. 10.1093/BIOINFORMATICS/BTI623.

78. Novakovsky, G., Dexter, N., Libbrecht, M.W., Wasserman, W.W., and Mostafavi, S. (2023). Obtaining genetics insights from deep learning via explainable artificial intelligence. Nat Rev Genet 24, 125–137. 10.1038/S41576-022-00532-2.

79. Shrikumar, A., Greenside, P., and Kundaje, A. (2017). Learning Important Features Through Propagating Activation Differences. 34th International Conference on Machine Learning, ICML 2017 *7*, 4844–4866. arXiv.1605.01713.

80. Lundberg, S.M., and Lee, S.I. (2017). A Unified Approach to Interpreting Model Predictions. Adv Neural Inf Process Syst 2017-December, 4766–4775.

81. Lundberg, S.M., Erion, G., Chen, H., DeGrave, A., Prutkin, J.M., Nair, B., Katz, R., Himmelfarb, J., Bansal, N., and Lee, S.I. (2020). From Local Explanations to Global Understanding with Explainable AI for Trees. Nat Mach Intell 2, 56–67. 10.1038/S42256-019-0138-9.

82. Shrikumar, A., Tian, K., Avsec, Ž., Shcherbina, A., Banerjee, A., Sharmin, M., Nair, S., and Kundaje, A. (2018). Technical Note on Transcription Factor Motif Discovery from Importance Scores (TF-MoDISco) version 0.5.6.5. 10.48550/arXiv.1811.00416.

83. Avsec, Ž., Weilert, M., Shrikumar, A., Krueger, S., Alexandari, A., Dalal, K., Fropf, R., McAnany, C., Gagneur, J., Kundaje, A., et al. (2021). Base-resolution models of transcription-factor binding reveal soft motif syntax. Nat Genet 53, 354–366. 10.1038/S41588-021-00782-6.

84. Alexandari, A.M., Horton, C.A., Shrikumar, A., Shah, N., Li, E., Weilert, M., Pufall, M.A., Zeitlinger, J., Fordyce, P.M., and Kundaje, A. (2023). De novo distillation of thermodynamic affinity from deep learning regulatory sequence models of in vivo protein-DNA binding. bioRxiv. 10.1101/2023.05.11.540401.

85. Abramson, J., Adler, J., Dunger, J., Evans, R., Green, T., Pritzel, A., Ronneberger, O., Willmore, L., Ballard, A.J., Bambrick, J., et al. (2024). Accurate structure prediction of biomolecular interactions with AlphaFold 3. Nature 630, 493–500. 10.1038/S41586-024-07487-W.

86. McInnes, L., Healy, J., and Melville, J. (2018). UMAP: Uniform Manifold Approximation and Projection for Dimension Reduction. ArXiv 1802.03426. 10.48550/arXiv.1802.03426.

87. Sloan, C.A., Chan, E.T., Davidson, J.M., Malladi, V.S., Strattan, J.S., Hitz, B.C., Gabdank, I., Narayanan, A.K., Ho, M., Lee, B.T., et al. (2016). ENCODE data at the ENCODE portal. Nucleic Acids Res 44, D726–D732. 10.1093/NAR/GKV1160.

88. Hitz, B.C., Jin-Wook, L., Jolanki, O., Kagda, M.S., Graham, K., Sud, P., Gabdank, I., Strattan, J.S., Sloan, C.A., Dreszer, T., et al. (2023). The ENCODE Uniform Analysis Pipelines. bioRxiv. 10.1101/2023.04.04.535623.

89. 89. Esteves de Lima, J., Bou Akar, R., Machado, L., Li, Y., Drayton-Libotte, B., Dilworth, F.J., and Relaix, F. (2021). HIRA stabilizes skeletal muscle lineage identity. Nat Commun 12. 10.1038/S41467-021-23775-9.

90. Clough, E., Barrett, T., Wilhite, S.E., Ledoux, P., Evangelista, C., Kim, I.F., Tomashevsky, M., Marshall, K.A., Phillippy, K.H., Sherman, P.M., et al. (2024). NCBI GEO: archive for gene expression and epigenomics data sets: 23-year update. Nucleic Acids Res 52, D138–D144. 10.1093/NAR/GKAD965.

91. Karolchik, D., Hinricks, A.S., Furey, T.S., Roskin, K.M., Sugnet, C.W., Haussler, D., and Kent, W.J. (2004). The UCSC Table Browser data retrieval tool. Nucleic Acids Res 32. 10.1093/NAR/GKH103.

92. Sollis, E., Mosaku, A., Abid, A., Buniello, A., Cerezo, M., Gil, L., Groza, T., Güneş, O., Hall, P., Hayhurst, J., et al. (2023). The NHGRI-EBI GWAS Catalog: knowledgebase and deposition resource. Nucleic Acids Res 51, D977–D985. 10.1093/NAR/GKAC1010.

93. Aguet, F., Barbeira, A.N., Bonazzola, R., Brown, A., Castel, S.E., Jo, B., Kasela, S., Kim-Hellmuth, S., Liang, Y., Oliva, M., et al. (2020). The GTEx Consortium atlas of genetic regulatory effects across human tissues. Science 369, 1318–1330. 10.1126/SCIENCE.AAZ1776.

94. Okbay, A., Wu, Y., Wang, N., Jayashankar, H., Bennett, M., Nehzati, S.M., Sidorenko, J., Kweon, H., Goldman, G., Gjorgjieva, T., et al. (2022). Polygenic prediction of educational attainment within and between families from genome-wide association analyses in 3 million individuals. Nat Genet 54, 437–449. 10.1038/S41588-022-01016-Z.

95. Schoeler, T., Speed, D., Porcu, E., Pirastu, N., Pingault, J.B., and Kutalik, Z. (2023). Participation bias in the UK Biobank distorts genetic associations and downstream analyses. Nat Hum Behav 7, 1216–1227. 10.1038/S41562-023-01579-9.

96. Thorvaldsdóttir, H., Robinson, J.T., and Mesirov, J.P. (2013). Integrative Genomics Viewer (IGV): high-performance genomics data visualization and exploration. Brief Bioinform 14, 178–192. 10.1093/BIB/BBS017.

97. Robinson, J.T., Thorvaldsdóttir, H., Winckler, W., Guttman, M., Lander, E.S., Getz, G., and Mesirov, J.P. (2011). Integrative genomics viewer. Nat Biotechnol 29, 24–26. 10.1038/NBT.1754.

98. Gu, Z., Gu, L., Eils, R., Schlesner, M., and Brors, B. (2014). circlize Implements and enhances circular visualization in R. Bioinformatics 30, 2811–2812. 10.1093/BIOINFORMATICS/BTU393.

99. Gu, Z., Eils, R., and Schlesner, M. (2016). Complex heatmaps reveal patterns and correlations in multidimensional genomic data. Bioinformatics 32, 2847–2849. 10.1093/BIOINFORMATICS/BTW313.

100. Gu, Z., Eils, R., Schlesner, M., and Ishaque, N. (2018). EnrichedHeatmap: an R/Bioconductor package for comprehensive visualization of genomic signal associations. BMC Genomics 19. 10.1186/S12864-018-4625-X.

101. Liberzon, A., Birger, C., Thorvaldsdóttir, H., Ghandi, M., Mesirov, J.P., and Tamayo, P. (2015). The Molecular Signatures Database (MSigDB) hallmark gene set collection. Cell Syst 1, 417. 10.1016/J.CELS.2015.12.004.

102. Liberzon, A., Subramanian, A., Pinchback, R., Thorvaldsdóttir, H., Tamayo, P., and Mesirov, J.P. (2011). Molecular signatures database (MSigDB) 3.0. Bioinformatics 27, 1739–1740. 10.1093/BIOINFORMATICS/BTR260.

103. 103. Carroll, T.S., Liang, Z., Salama, R., Stark, R., and de Santiago, I. (2014). Impact of artifact removal on ChIP quality metrics in ChIP-seq and ChIP-exo data. Front Genet 5. 10.3389/FGENE.2014.00075/ABSTRACT.

104. Okonechnikov, K., Conesa, A., and García-Alcalde, F. (2016). Qualimap 2: advanced multi-sample quality control for high-throughput sequencing data. Bioinformatics 32, 292–294. 10.1093/BIOINFORMATICS/BTV566.

